# Acyl Carrier Protein is Essential for Apicoplast Biogenesis in Malaria Parasites Independent of Fatty Acid Synthesis

**DOI:** 10.64898/2026.03.29.713301

**Authors:** Sage W. R. Geher, Seyi Falekun, Jessica N. Pita-Aquino, Russell P. Swift, Megan Okada, Yasaman Jami-Alahmadi, James A. Wohlschlegel, Sean T. Prigge, Paul A. Sigala

## Abstract

Acyl carrier protein (ACP) and its 4-phosphopantetheine prosthetic group canonically function as the soluble scaffold for acyl chain assembly and elongation during type-II fatty acid synthesis (FASII). *Plasmodium* malaria parasites retain a FASII pathway in the apicoplast organelle that has been the subject of considerable scrutiny and confusion. Although apicoplast FASII is essential for *P. falciparum* growth within mosquitoes and the human liver, this pathway is dispensable and largely inactive in blood-stage parasites that can scavenge host fatty acids. In contrast to FASII enzymes that can be disrupted without fitness defect, we report that knockout or ligand-dependent knockdown of apicoplast ACP is lethal to blood-stage *P. falciparum*, indicating an essential FASII-independent function. Loss of ACP impairs the biosynthesis of essential isoprenoid precursors and blocks apicoplast biogenesis. Using proximity biotinylation and biochemical interaction studies, we identified a key role for ACP in binding and stabilizing apicoplast pyruvate kinase II (PKII). This critical enzyme is the only known source of nucleoside triphosphates (NTPs) in this organelle and is required for isoprenoid synthesis and apicoplast biogenesis. Our work reveals that ACP knockdown results in destabilization and loss of PKII, which is sufficient to explain ACP essentiality in this stage. This work unveils essential ACP function at a key biochemical hub controlling broad apicoplast metabolism in malaria parasites that is independent of the canonical ACP role in FASII.

## INTRODUCTION

Malaria remains a pressing global health threat, with a disproportionate impact on under-resourced communities in Africa that is likely to worsen due to funding cutbacks by key donor countries.^1^ At the same time, the efficacy of current therapies is threatened by ongoing resistance development by virulent *Plasmodium falciparum* parasites.^2, 3^ These concerns highlight the critical need to deepen understanding of *Plasmodium* biology to optimize use of current medicines and to develop new therapeutic strategies.

*Plasmodium* parasites retain two critical organelles, the mitochondrion and a non-photosynthetic plastid termed the apicoplast, which carry out critical metabolic functions and are validated drug targets.^4–7^ Since human cells lack an apicoplast, this organelle and its prokaryotic-like pathways were considered to be fertile ground for discovering new parasite-specific drug targets. However, the identification of essential pathways within the apicoplast has been challenging, since many of its metabolic functions are dispensable during parasite growth within red blood cells (RBCs) and are only required in the mosquito and/or liver stages. For example, biosynthetic pathways in the apicoplast for making acetyl CoA, lipoate, fatty acids, and heme biosynthesis intermediates are all dispensable during blood-stage growth.^8–15^ Indeed, biosynthesis of the isoprenoid precursor, isopentenyl pyrophosphate (IPP), and the final step in production of coenzyme A are the only known essential apicoplast pathways whose metabolic outputs are required outside the organelle.^5, 16^ However, pathways that support apicoplast maintenance and IPP synthesis,^17, 18^ including protein and DNA synthesis,^19–21^ RNA transcription,^22, 23^ Fe-S cluster biogenesis,^24, 25^ and (d)NTP production^26^ are also essential for parasite viability.

Fatty acid synthesis in the apicoplast has been the subject of considerable confusion and controversy since the sequencing of the *P. falciparum* genome.^27–29^ The apicoplast encodes a type II fatty acid biosynthesis (FASII) pathway for making octanoic acid used in lipoate biosynthesis and for elongation into longer-chain fatty acids.^27^ FASII activity involves over seven enzymes of prokaryotic origin for initiation, modification, and elongation of a growing acyl chain tethered to the terminal thiol of a 4-phosphopantetheine group (4-PP) on the acyl carrier protein (ACP, Pf3D7_0208500), which serves as the soluble scaffold for fatty acid synthesis (Figure 1).^30^ This pathway was originally proposed to be essential for blood-stage parasites, based on the anti-*Plasmodium* activity of triclosan, an inhibitor of the FASII enoyl-ACP reductase enzyme (FabI) in bacteria.^31^ However, subsequent gene-expression and knockout studies of FabI and other FASII enzymes and isotope-tracing experiments clarified that this pathway is dispensable and largely inactive in blood-stage *Plasmodium*, which can scavenge host-derived fatty acids.^13, 14, 32^ Indeed, FASII enzymes only appear to be critical for the highly proliferative stages of *P. falciparum* growth in mosquitoes and the human liver.^14, 15^

**Figure 1.**
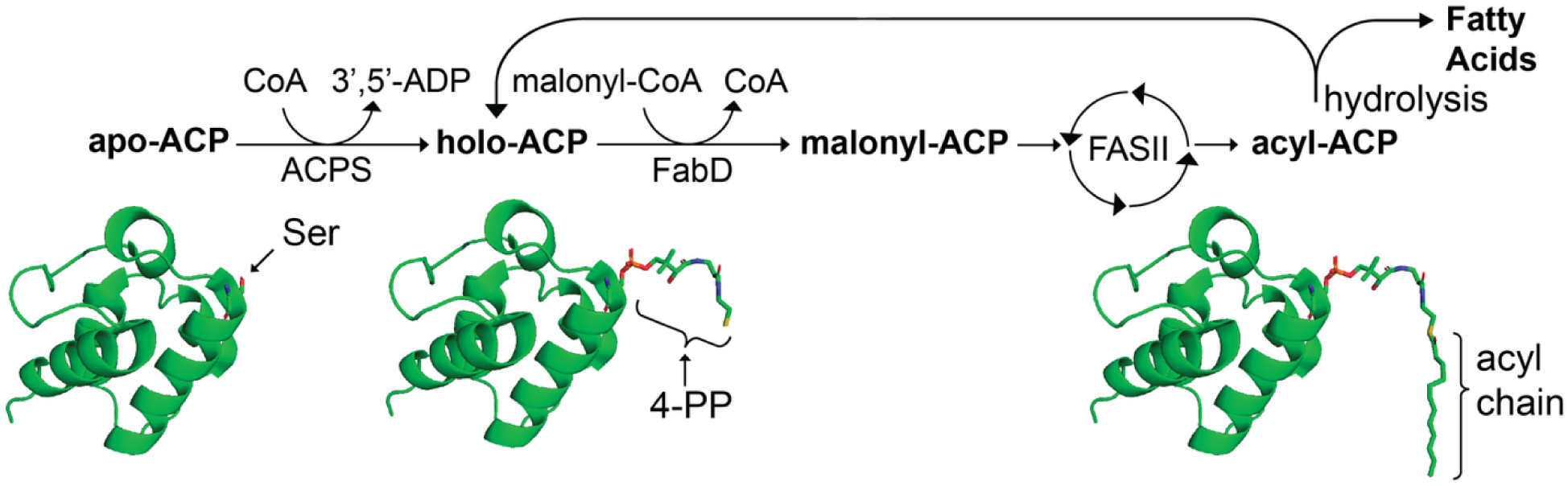
Schematic depiction of apicoplast ACP and the FASII pathway in *P. falciparum*. CoA = coenzyme A, ACPS = holo-ACP synthase, FabD = malonyl-CoA:ACP S-malonyltransferase, 4-PP = 4-phosphopantetheine. The ACP structural models are based on the X-ray structures of *P. falciparum* holo-aACP (PDB 3GZM)^36^ and *E. coli* acyl-ACP (PDB 5USR)^45^.

Although FASII activity is dispensable for blood-stage parasites, we noted that genome-wide knockout and insertional mutagenesis studies in *P. berghei* and *P. falciparum* reported that apicoplast ACP could not be disrupted, suggesting an essential function independent of FASII.^33, 34^ ACP is a small, highly α-helical protein that contains a strictly conserved Ser residue modified by the 4-PP group derived from coenzyme A (Figure 1).^35, 36^ The 4-PP group is attached to ACP by the holo-ACP synthase enzyme (Pf3D7_0420200), which is also predicted to be essential for blood-stage *P. falciparum*.^33^ Organelle targeting and proteolytic processing of ACP have been heavily studied in *P. falciparum*,^37, 38^ but ACP roles apart from FASII have not been identified in the parasite apicoplast or in related plastids like chloroplasts.^39^ However, expanded ACP functions beyond acyl chain synthesis are well known in mitochondria. Indeed, most eukaryotes retain a complete mitochondrial FASII pathway whose ACP also plays a critical role binding and stabilizing key proteins involved in Fe-S cluster biogenesis and assembly of respiratory chain complexes and (in mammals) of mitochondrial ribosomes.^40–43^ In contrast to most eukaryotes, *P. falciparum* and other apicomplexan parasites lack mitochondrial FASII. Nevertheless, they retain an essential ACP homolog (Pf3D7_1208300) in this organelle that uses a divergent molecular interface lacking a 4-PP group to bind and stabilize the Isd11-Nfs1 complex required for mitochondrial Fe-S cluster biogenesis.^44^

We set out to test and understand the FASII-independent function of apicoplast ACP in blood-stage *P. falciparum*. ACP knockout and knockdown studies confirmed an essential role for this protein in apicoplast biogenesis and IPP synthesis. Biochemical interaction studies uncovered a critical stabilizing interaction between ACP and the apicoplast pyruvate kinase II enzyme that is essential for (d)NTP and pyruvate synthesis required for broad apicoplast metabolism. This interaction appears to require the 4-PP prosthetic group on ACP, whose modification status may differentially regulate PKII activity and thus apicoplast function in distinct host environments. Our study unveils a new molecular paradigm for essential ACP function at a key metabolic hub in the apicoplast of malaria parasites that is independent of the canonical ACP role in FASII.

## RESULTS

### ACP is essential for parasite viability and apicoplast biogenesis

Prior studies successfully deleted most FASII enzymes in blood-stage *Plasmodium*, but there have been no reported attempts to disrupt apicoplast ACP.^13^ To directly test ACP essentiality for blood-stage parasites, we targeted the ACP gene for deletion in PfMev NF54 parasites.^46^ These parasites express apicoplast-targeted GFP and have been engineered to use exogenous mevalonate to synthesize IPP in the cytoplasm, which bypasses the endogenous apicoplast IPP synthesis pathway and rescues parasites from lethal apicoplast defects. Parasites successfully returned from transfection only in the presence of mevalonate, and genomic PCR analysis confirmed successful deletion of the apicoplast ACP gene (Pf3D7_0208500). Growth of ΔACP parasites was strictly dependent on exogenous mevalonate (Figure 2A and Figure 2- figure supplement 1). Fluorescence microscopy revealed that the apicoplast-targeted GFP signal in ΔACP parasites displayed a dispersed constellation of GFP foci, suggesting organelle disruption due to a defect in apicoplast biogenesis and inheritance (Figure 2B and Figure 2- figure supplement 2). PCR analysis confirmed selective loss of the apicoplast genome in these parasites (Figure 2B), supporting the conclusion that loss of ACP results in a lethal defect in apicoplast biogenesis and IPP synthesis.

**Figure 2.**
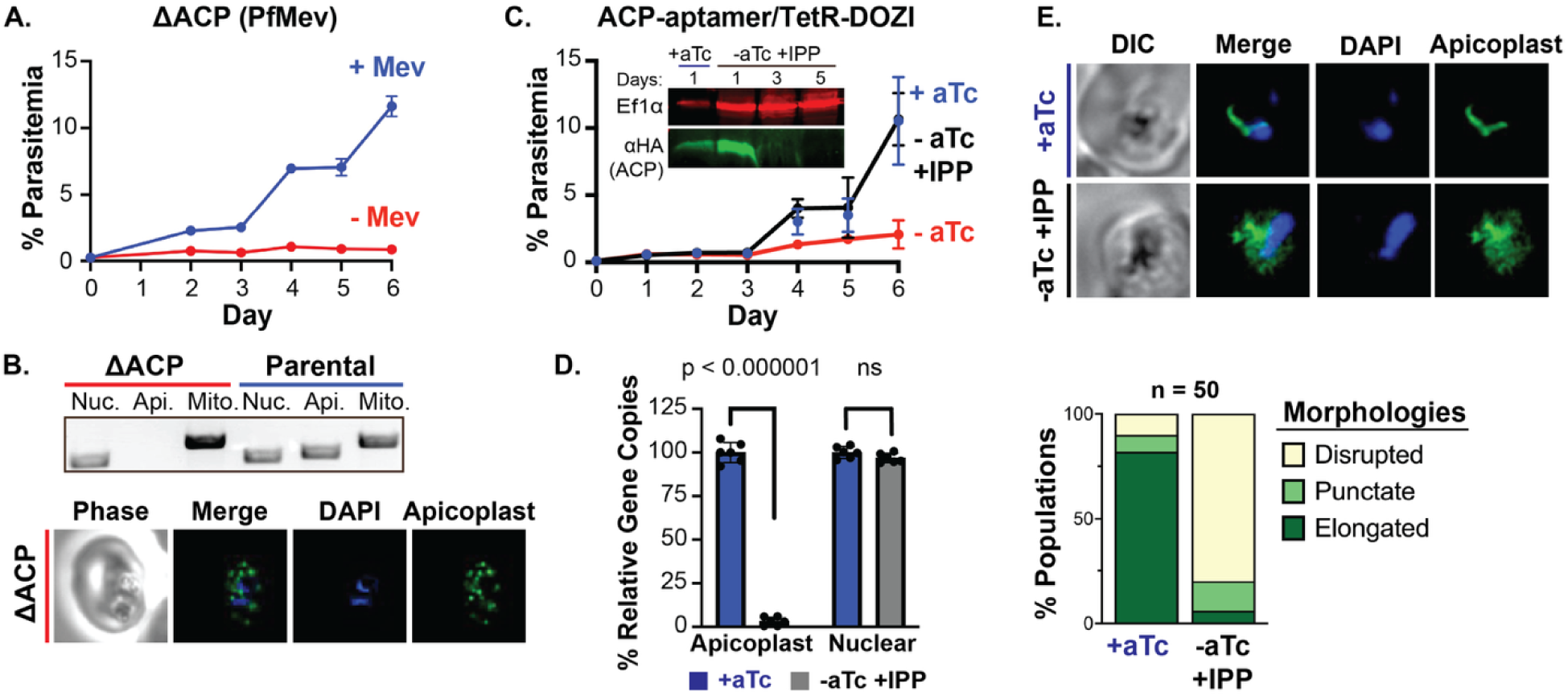
Knockout or knockdown of ACP expression blocks parasite growth and apicoplast biogenesis. (**A**) Synchronous growth assay of ΔACP PfMev parasites cultured ±50 µM mevalonate (Mev). Parasitemia values are the average ± SD from biological triplicate samples. (**B**) Genomic PCR analysis and live parasite imaging show selective loss of the apicoplast genome and disrupted apicoplast morphology based on fluorescence of the ACP_L_-GFP marker protein in PfMev parasites. Nuc. = nuclear gene (LDH, Pf3D7_1324900), Api. = apicoplast gene (SufB, Pf3D7_API04700), Mito. = mitochondrial gene (CoxI, Pf3D7_MIT02100). (**C**) Synchronous growth assay of apicoplast ACP-aptamer/TetR-DOZI Dd2 parasites cultured ±1 µM aTc and ±200 µM IPP. Parasitemia values are the average ± SD from biological triplicates. Inset: western blot analysis of parasites harvested after 1, 3, or 5 days at the indicated growth conditions and probed with anti-EF1α (cytosolic loading control) or anti-HA (ACP) antibodies. (**D**) Quantitative PCR analysis of the apicoplast:nuclear genome ratio for parasites in panel C cultured 84 hours in the indicated conditions, based on amplification of apicoplast (SufB, ClpM: Pf3D7_API03600, TufA: Pf3D7_API02900) relative to nuclear (STL: Pf3D7_0717700, I5P: Pf3D7_0802500, ADSL: Pf3D7_0206700) genes. Indicated qPCR ratios were normalized to +aTc and are the average ± SD of biological triplicates. Significance was analyzed by unpaired Student’s t-test to determine the indicated p values. (**E**) Immunofluorescence microscopy of parasites in panel C cultured for 5 days (120 hours) in the indicated conditions and stained with anti-apicoplast ACP or DAPI (nucleus). Below: population analysis of apicoplast morphology scored for disrupted (dispersed), punctate, or elongated GFP signal in 50 total parasites from biological triplicates. **Figure 2- figure supplement 1.** Integration PCRs for deletion or modification of the ACP gene. **Figure 2- figure supplement 2.** Additional microscopy images ΔACP PfMev NF54 parasites. **Figure 2- figure supplement 3.** Growth assays, fluorescence microscopy, and genomic qPCR of ACP knockdown in PfMev NF54 parasites. **Figure 2- figure supplement 4.** Additional IFA images of ACP knockdown in Dd2 parasites. **Figure 2- figure supplement 5.** Validation of custom anti-ACP antibody by western blot. **Figure 2- source data 1.** Uncropped gel images for integration and organelle genome PCR analyses of ACP knockout or knockdown parasites. **Figure 2- source data 2.** Uncropped western blot gel image of ACP levels upon knockdown. **Figure 2- source data 3.** Uncropped gel image for validation of custom anti-ACP antibody.

To further test this conclusion and to dissect ACP function in the apicoplast, we used CRISPR/Cas9 to tag the ACP gene in both Dd2 and PfMev NF54 parasites to encode a C-terminal HA-FLAG epitope tag fusion and the aptamer-TetR/DOZI system for conditional knockdown.^47^ Successful integration in clonal parasites for both lines was confirmed by genomic PCR (Figure 2- figure supplement 1). Parasites grew normally in the presence of exogenous anhydrotetracycline (aTc), which enables normal ACP translation and detection of HA-FLAG-tagged ACP at the expected size by western blot analysis (Figure 2C). However, parasite growth was strongly suppressed after 2-3 cycles following aTc washout that resulted in loss of detectable ACP expression (Figure 2C and Figure 2- figure supplement 3). In both Dd2 and PfMev parasites, culture supplementation with exogenous IPP or mevalonate, respectively, fully restored normal parasite growth in the absence of aTc, confirming an apicoplast-specific defect upon ACP knockdown. Fluorescence microscopy and genomic qPCR analysis in IPP-rescued parasites confirmed disruption of the apicoplast and selective loss of the apicoplast genome upon sustained culture without aTc (Figure 2D and 2E, Figure 2- figure supplement 2, and Figure 2-figure supplement 3). These results strongly support the conclusion that apicoplast ACP is essential for parasite viability and organelle biogenesis.

### Blood-stage parasites require ACP modification by holo-ACP synthase but not FabD

Canonical ACP function requires modification by holo-ACP synthase (ACPS), which uses coenzyme A to transfer a 4-PP group to a highly conserved Ser residue on ACP (Figure 1). Apicoplast ACP retains this conserved Ser (Figure 1), and parasites express an apicoplast-targeted ACPS enzyme (Pf3D7_0420200), ^27, 45^ as expected for 4-PP attachment to ACP to support essential FASII activity in mosquito- and liver-stage parasites.^14, 15^ Because FASII activity is dispensable in blood-stage parasites, the modification state of ACP and role of ACPS in this stage have been uncertain.^36^

To directly test if ACPS function is essential for blood-stage parasites, we targeted the ACPS gene for deletion in PfMev NF54 parasites. Like ACP, parasites only returned from transfection in the presence of exogenous mevalonate, and genomic PCR analysis confirmed disruption of the ACPS gene (Figure 3- figure supplement 1). Growth of ΔACPS parasites was strictly dependent on mevalonate (Figure 3A), and analysis by fluorescence microscopy (Figure 3B- and Figure 3- figure supplement 2) and genomic PCR (Figure 3C) revealed disruption of apicoplast morphology and selective loss of the apicoplast genome. These results indicate that ACPS function is essential for parasite viability and apicoplast biogenesis, as expected if 4-PP attachment is required for essential ACP function in blood-stage parasites. Additional biochemical results described below also support the conclusion that essential ACP function requires 4-PP attachment.

**Figure 3.**
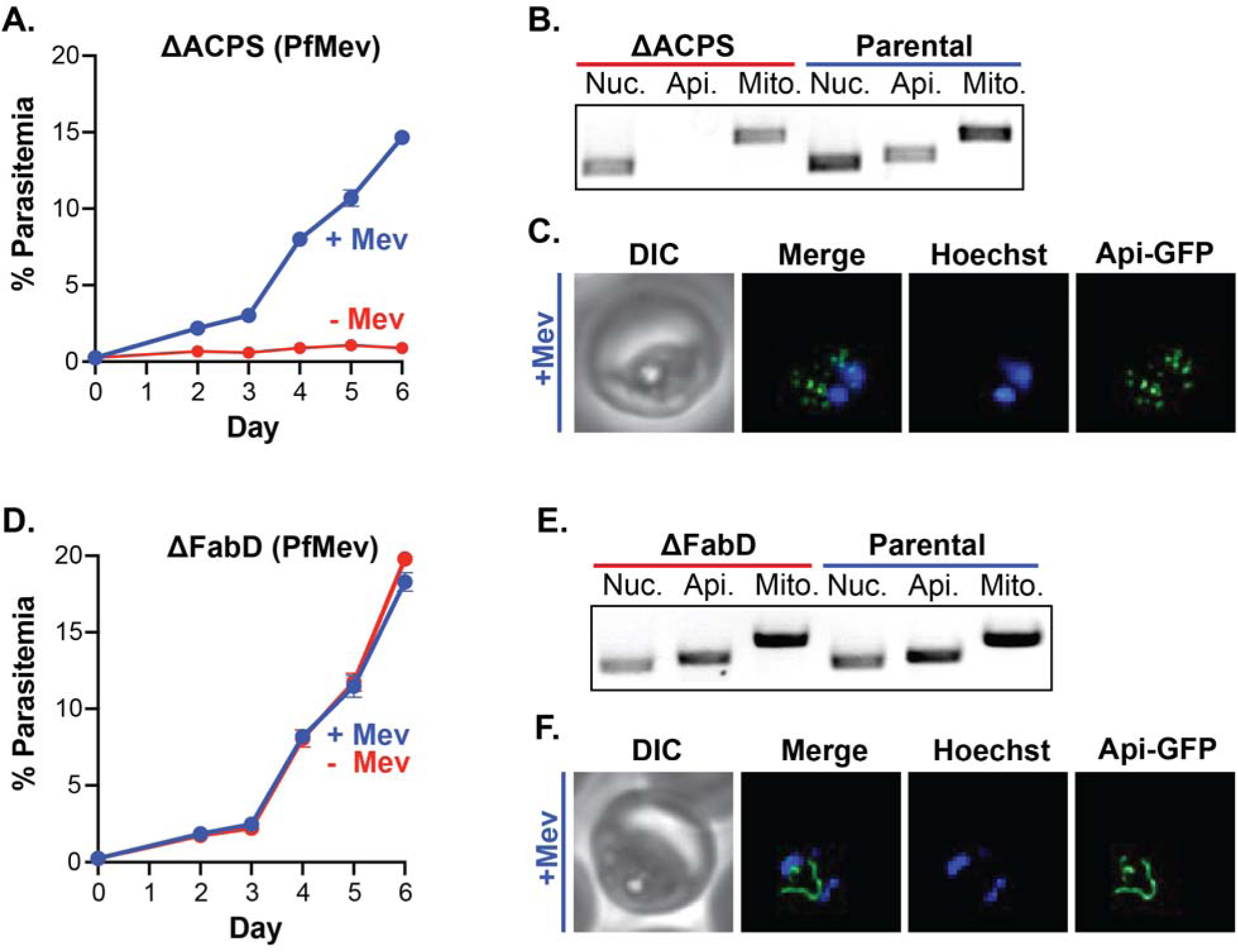
Holo-ACP synthase but not FabD is essential for blood-stage parasites and apicoplast biogenesis. (**A)** Synchronous growth assay of ΔACPS PfMev parasites cultured ±50 µM mevalonate. **(B)** Genomic PCR analysis and (**C**) live parasite imaging show selective loss of the apicoplast genome and disrupted apicoplast morphology based on fluorescence of the ACP_L_-GFP marker protein in PfMev parasites. (**D**) Synchronous growth assay of ΔFabD PfMev parasites cultured ±50 µM mevalonate. **(E)** Genomic PCR analysis and (**F**) live parasite imaging indicate retention of the apicoplast genome and normal apicoplast morphology based on ACP_L_-GFP fluorescence in ΔFabD parasites. Parasitemia values are the average ± SD from biological triplicates. Nuc. = nuclear gene (LDH), Api. = apicoplast gene (SufB), Mito. = mitochondrial gene (CoxI), DIC = differential interference contrast. **Figure 3- figure supplement 1.** Integration PCRs for deletion of ACPS and FabD in PfMev parasites. **Figure 3- figure supplement 2**. Additional microscopy images of apicoplast morphology in ΔACPS and ΔFabD parasites. **Figure 3- figure supplement 3**. Growth of ΔACP and ΔFabD PfMev parasites in low-lipid conditions. **Figure 3- source data 1**. Uncropped gel images for integration and apicoplast status PCRs for ΔACP and ΔFabD parasites.

The initial step in FASII activity involves FabD-catalyzed transfer of a malonyl group from malonyl-CoA to holo-ACP to form malonyl-ACP (Figure 1). Although downstream FASII enzymes that catalyze successive modification and elongation of malonyl-ACP into acyl chains of variable length have been successfully deleted in blood-stage *P. falciparum*,^13^ there are no reports of targeted disruption of FabD (Pf3D7_1312000). Prior genome-wide studies in blood-stage parasites have reported conflicting results, with FabD listed as essential in *P. falciparum*^33^ but dispensable in *P. berghei*.^34^ To test if ACP modification by FabD is required for essential ACP function, we targeted FabD for deletion in PfMev NF54 parasites. Parasites returned from transfection without exogenous mevalonate, and genomic PCR analysis confirmed successful deletion of the FabD gene (Figure 3- figure supplement 1). Growth of ΔFabD parasites was identical in ±mevalonate conditions (Figure 3D), and analysis by fluorescence microscopy (Figure 3E and Figure 3- figure supplement 2) and genomic PCR (Figure 3F) indicated normal apicoplast morphology and genome status. We conclude that FabD, like other FASII enzymes, is dispensable for blood-stage *P. falciparum* and not required for essential ACP function.

Prior isotope-tracing studies suggested that apicoplast FASII is largely inactive in blood-stage parasites growing in standard media but may be activated in low-lipid conditions.^32, 48^ A subsequent study thus suggested that the apicoplast FASII pathway might be essential for blood-stage *P. falciparum* growth in low-lipid conditions.^49^ We tested this suggestion by growing ΔACP and ΔFabD PfMev parasites in low-lipid conditions, in which Albumax was replaced in the culture medium with lipid-free bovine serum albumin supplemented with oleic and palmitic acids with daily media changes.^32, 50^ Both knockout lines grew steadily in low-lipid conditions, and ΔACP parasites with disrupted apicoplast retained exclusive dependence on exogenous mevalonate for viability (Figure 3- figure supplement 3). These results contrast with the prior study^49^ of ΔFabI parasites and the proposed model that blood-stage *P. falciparum* requires FASII activity for growth in low-lipid conditions and suggest that parasites can rely on scavenging host-derived fatty acids over a wide range of lipid conditions. Future studies involving tandem growth and isotope-labeling experiments of WT and ΔFASII parasites will be required to fully test and understand FASII function and the dependence of *P. falciparum* growth on this pathway in low-lipid conditions.

### ACP associates with apicoplast pyruvate kinase II using its 4-PP group

Beyond its role in FASII, mitochondrial ACP in other eukaryotes binds to LYR motif adapter proteins that interact with diverse protein partners involved in Fe-S cluster biogenesis and assembly of mitochondrial respiratory complexes and (in humans) mitoribosomes.^41, 51^ Expanded roles for ACP beyond FASII in chloroplasts and other plastids are sparsely defined, and there is no evidence for LYR motif proteins in plastids, including the *P. falciparum* apicoplast.^44, 52^ Nevertheless, based on strong biochemical precedent that ACP in mitochondria and bacteria engage in a variety of critical protein-protein interactions, we hypothesized that essential ACP function in the parasite apicoplast might involve an essential interaction with one or more key proteins required for organelle function and biogenesis.^53, 54^

To identify proximal interacting proteins of the apicoplast ACP, we first took a proximity biotinylation approach by episomally expressing C-terminal fusions of apicoplast ACP with miniTurbo or BioID2 in blood-stage Dd2 *P. falciparum*.^55, 56^ Biotinylation of proteins that spanned a range of molecular masses was exclusively detected in transgenic but not parental Dd2 parasites (Figure 4- figure supplement 1). Labeled proteins were identified by tandem mass spectrometry after pull-down with streptavidin resin, stringent washes, and tryptic digest. Pull-downs of parental Dd2 parasites served as negative control. Over fifty apicoplast-targeted proteins were detected as specifically enriched in parasites expressing ACP-miniTurbo, with apicoplast pyruvate kinase II (PKII, Pf3D7_1037100) showing the strongest enrichment (Figure 4A and Supplementary File 1- supplementary table 1). Comparatively fewer proteins were identified from ACP-BioID2-expressing parasites, but PKII was one of two apicoplast-targeted proteins detected (Figure 4- figure supplement 2). We thus considered the possibility that apicoplast ACP had a functionally relevant interaction with PKII.

**Figure 4.**
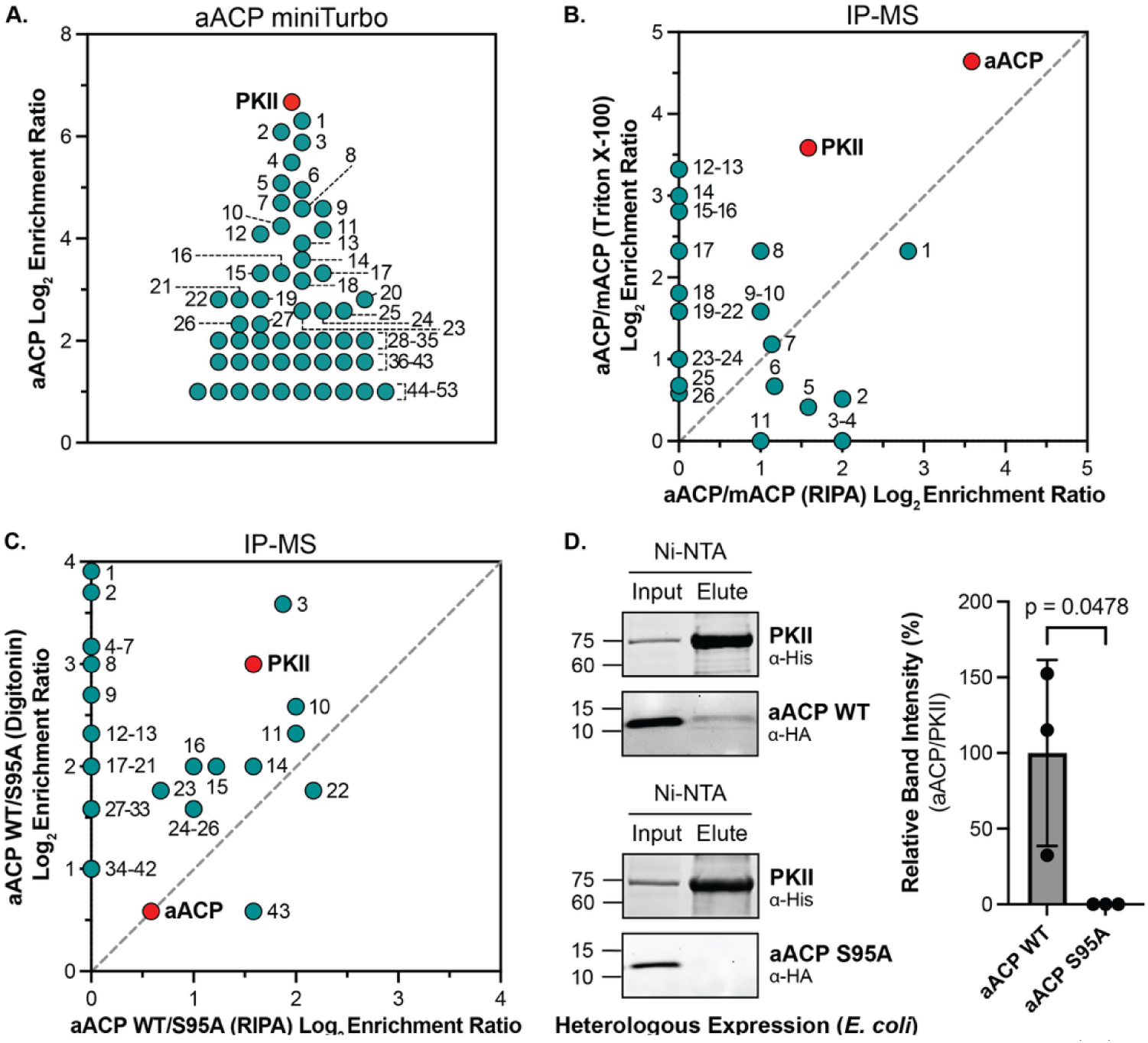
Apicoplast ACP associates with apicoplast pyruvate kinase II. (**A**) Proximity biotinylation study of Dd2 parasites episomally expressing apicoplast ACP tagged with C-miniTurbo, showing log_2_ enrichment ratio of known apicoplast-targeted proteins compared to parental Dd2 parasites, based on spectral intensity of proteins detected by tandem mass spectrometry. (**B**) Immunoprecipitation-mass spectrometry (IP-MS) study of apicoplast ACP interacting proteins, based on anti-HA-tag IP of apicoplast versus mitochondrial ACP-HA_2_ in Dd2 parasites lysed in Triton X-100 or RIPA buffer. Axes display the log_2_ enrichment ratio for detection of each protein in IP samples of aACP versus mACP. Dashed diagonal line has a slope of one. (**C**) IP-MS analysis based on anti-HA-tag IP of lysates from Dd2 parasites episomally expressing wildtype (WT) or the S95A mutant of apicoplast ACP-HA_2_ and lysed in digitonin or RIPA buffer. Axes display the log_2_ enrichment ratio for detection of each protein in IP samples of WT versus S95A ACP. Dashed diagonal line has a slope of one. The identity of numbered proteins in panels A-C is shown in Supplementary File 1. A list of all proteins detected by mass spectrometry is shown in Figure 4- source data 2. (**D**) Affinity pull-down and western blot analysis of lysates from *E. coli* bacteria heterologously expressing parasite His_6_-tagged PKII or apicoplast ACP-HA_2_ (WT or S95A mutant). PKII was affinity isolated from bacterial lysates using nickel-nitrilotriacetic acid (Ni-NTA) resin, eluted with free imidazole, and analyzed by western blot for co-purification with ACP. Membranes were probed with anti-His_6_ and anti-HA-tag antibodies. The signal intensity for WT or S95A ACP relative to PKII in pulldown samples was quantified by densitometry in biological triplicate samples and plotted as the average ± SD, with significance analyzed by Student’s t-test to determine the indicated p value. **Figure 4- figure supplement 1.** Western blot detection of biotinylated proteins in lysates from parental, aACP-miniTurbo, or aACP-BioID2 Dd2 parasites. **Figure 4- figure supplement 2.** Enriched protein interactors of apicoplast ACP detected by proximity biotinylation or immunoprecipitation. **Figure 4- figure supplement 3.** Replicate western blot images for Ni-NTA pulldown of PKII association with WT or Ser95Ala ACP in *E. col* **Supplementary File 1- supplementary table 1.** Table of apicoplast-targeted proteins identified in parasites expressing ACP-miniTurbo or ACP-BioID2. **Supplementary File 1- supplementary table 2.** Table of apicoplast-targeted proteins identified by IP/MS of apicoplast ACP-HA_2_. **Supplementary File 1- supplementary table 3.** Table of apicoplast-targeted proteins identified by IP/MS of WT or Ser95Ala ACP-HA_2_. **Supplementary File 1- supplementary table 4.** Table of apicoplast-targeted protein interactors of aACP identified by both miniTurbo and IP/MS. **Figure 4- source data 1.** Uncropped western blot for Figure 4- figure supplement 1 **Figure 4- source data 2.** Excel file of all proteins detected in proximity biotinylation and immunoprecipitation studies of apicoplast ACP. **Figure 4- source data 3.** Uncropped western blots for Figure 4D and Figure 4- figure supplement 3.

Proximity biotinylation does not require stable interaction between bait and labeled proteins. To identify apicoplast proteins that stably associate with ACP, we performed immunoprecipitation/tandem mass spectrometry (IP/MS) studies of Dd2 parasites episomally expressing ACP with a C-terminal HA_2_ tag. We lysed parasites in either RIPA or 1% Triton X-100 buffer, with parasites expressing mitochondrial ACP-HA_2_ (Pf3D7_1208300) serving as negative control.^44^ Over twenty apicoplast-targeted proteins were identified in both lysis conditions, with PKII identified as the most or nearly most-enriched protein interactor of apicoplast ACP in both samples (Figure 4B, Figure 4- figure supplement 2, and Supplementary File 1- supplementary table 2). Apicoplast PKII is an essential enzyme whose production of pyruvate and a range of NTPs and dNTPs is critical to support apicoplast biogenesis and nearly all apicoplast metabolic pathways, including IPP synthesis by the non-mevalonate pathway, Fe-S cluster biogenesis by the Suf pathway, DNA replication, RNA transcription, and protein translation.^26^ On the basis of this essentiality and consistent detection of PKII as the most or nearly most-enriched interactor of ACP across four proximity biotinylation and IP/MS experiments, we focused our attention on the association of apicoplast ACP and PKII and the physiological consequence of this interaction.

Our observation that loss of ACPS impairs apicoplast biogenesis (Figure 3) suggested most simply that essential function of apicoplast ACP requires post-translational attachment of 4-PP. To test a role for 4-PP attachment to ACP in mediating association with PKII, we performed IP/MS studies of parasites episomally expressing wildtype (WT) or the Ser95Ala mutant of ACP-HA_2_. Ser95 is the strictly conserved ACP residue required for 4-PP attachment by ACPS (Figure 1),^36^ and its mutation thus ablates the ability of ACP to covalently bind 4-PP.^44^ Parasites were lysed in either RIPA or digitonin buffer prior to identification of associated proteins by anti-HA-tag IP/MS. Over forty apicoplast-targeted proteins were identified across both lysis conditions as enriched in samples of WT over Ser95Ala ACP (Figure 4C and Figure 4- figure supplement 2). PKII was strongly detected as interacting with WT ACP but was not detected in IP/MS samples of Ser95Ala ACP (Supplementary File 1- supplementary table 3), strongly suggesting that the 4-PP group on ACP contributes critically to its association with PKII.

Our IP experiments in parasites cannot distinguish if association between apicoplast ACP and PKII is direct or indirect or requires other apicoplast proteins. To test if apicoplast ACP can bind to PKII without other *Plasmodium* proteins, we heterologously expressed ACP-HA_2_ and His_6_-PKII in *E. coli* bacteria and performed affinity pull-downs of PKII using Ni-NTA resin to bind the His_6_ tag. The expressed proteins corresponded to the mature sequences expected upon apicoplast import and N-terminal processing.^26, 38, 44^ Recombinant PKII isolated in this fashion co-purified with WT but not Ser95Ala ACP (Figure 4D and Figure 4- figure supplement 3). These observations support the conclusion that apicoplast ACP can bind to PKII without other parasite proteins, as expected for a direct interaction, and that the 4-PP group on ACP contributes critically to this association.

### Loss of ACP destabilizes PKII and impairs apicoplast DNA and RNA levels

In eukaryotes, mitochondrial ACP interaction with key protein partners contributes to their stability and function, such that loss of ACP destabilizes associating proteins and can lead to their degradation.^41, 57^ Based on this biochemical precedent, we posited that apicoplast association might enhance the stability of PKII, such that loss of ACP would destabilize and impair PKII levels and function and thus explain the lethal defect in apicoplast biogenesis observed upon ACP knockdown in *P. falciparum*. To test this model, we transfected Dd2 ACP knockdown parasites with a plasmid encoding expression of PKII with C-terminal Myc_3_ tag. We observed that ACP knockdown after 3 days of growth in −aTc/+IPP conditions resulted in selective ∼75% reduction in detectable PKII levels by western blot analysis (Figure 5A and 5B and Figure 5-figure supplement 1). This observation directly supports our model that ACP association enhances the stability and levels of PKII in the parasite apicoplast.

**Figure 5.**
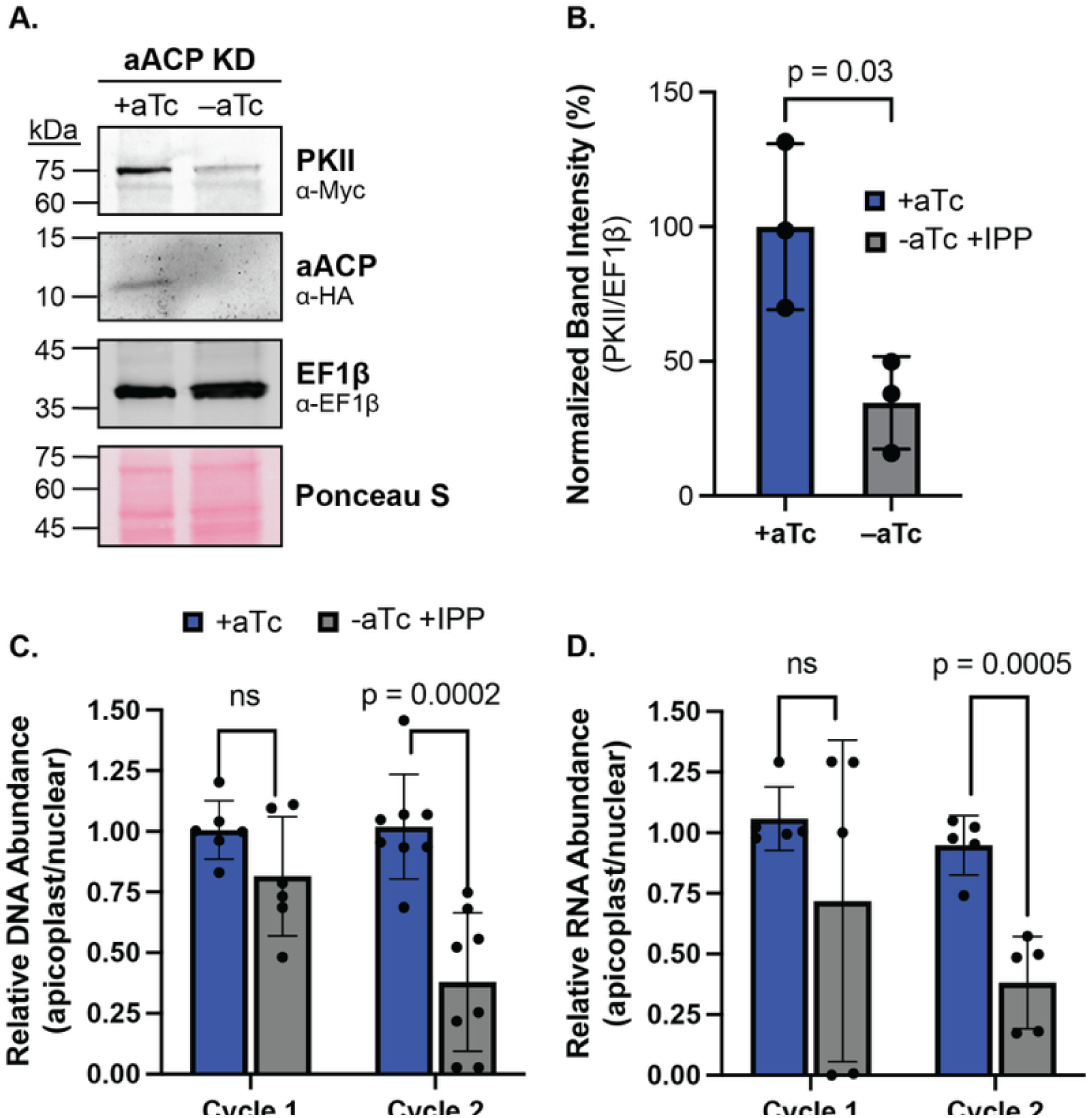
ACP knockdown destabilizes PKII and impairs apicoplast DNA and RNA levels. (A) Western blot analysis of apicoplast ACP-aptamer/TetR-DOZI Dd2 parasites episomally expressing pyruvate kinase II (PKII) C-Myc_3_ and cultured for 3 days ±1 µM aTc with 200 µM IPP. The blot was probed with α-Myc (PKII), α-HA (ACP), and α-EF1β (cytosolic loading control) antibodies and stained with Ponceau S for total protein. (B) Densitometry quantification of PKII and ACP protein levels normalized to EF1β based on western blot analysis of three biological replicates. Graph depicts the average ± SD, normalized to +aTc conditions. (C) Quantitative PCR analysis of DNA isolated from biological triplicate samples of apicoplast ACP-aptamer/TetR-DOZI parasites cultured for 36 or 84 hours ±1 µM aTc with 200 µM IPP, with normalization of Ct values averaged from three apicoplast genes (SufB, TufA, ClpM) to Ct values averaged from three nuclear genes (STL, I5P, ADSL). Graphs depict the average ± SD of each timepoint, normalized to +aTc conditions. (D) Quantitative RT-PCR analysis of RNA isolated from biological triplicate samples obtained from parasites cultured under identical conditions as for panel C to determine normalized levels of apicoplast (SufB, TufA, ClpM) relative to nuclear (STL, I5P, ADSL) transcripts. Indicated p values were determined by Student t-test analysis. **Figure 5-** figure supplement 1. Replicate western images for ACP knockdown and PKII levels **Figure 5**- source data 1. Uncropped western images for Figure 5- figure supplement 1.

Current evidence suggests that PKII is the dominant or exclusive source of NTPs and dNTPs in the apicoplast, with PKII knockdown causing defects in apicoplast genome transcription and ATP levels.^26^ To test if ACP knockdown and attendant decreases in PKII levels substantially reduced apicoplast-encoded DNA and RNA levels, we used qPCR and RT-qPCR to quantify apicoplast- versus nuclear-encoded gene and transcript levels in +aTc or −aTc/+IPP conditions in the first and second RBC growth cycles prior to apicoplast disruption. No significant differences in normalized DNA or RNA levels were observed in the first growth cycle ±aTc. However, ACP knockdown in −aTc conditions resulted in a nearly 70% reduction in apicoplast DNA and RNA levels in the second cycle, as expected for loss of PKII. This critical role for ACP in stabilizing PKII is sufficient to explain FASII-independent function and essentiality of apicoplast ACP.

## DISCUSSION

Organelles are central hubs of cellular metabolism and key drivers of eukaryotic evolution, including biochemical adaptations that specialize intracellular parasites to survive and grow within host cells. The apicoplast FASII pathway in *P. falciparum* has generated substantial interest and controversy as a possible antimalarial drug target.^13^ This pathway was initially thought to be essential for blood-stage parasites,^31^ but genetic knockouts of most FASII enzymes and isotope-tracing studies indicated that it was dispensable and largely inactive for parasites during RBC infection.^13, 14, 32^ In contrast to FASII, we have uncovered an essential blood-stage function for the apicoplast acyl carrier protein and its 4-PP prosthetic group in binding and stabilizing pyruvate kinase II, which supports nearly all aspects of apicoplast function and biogenesis. This essential interaction, which depends on the ACPS enzyme for 4-PP attachment to ACP, is independent of canonical ACP function as a soluble scaffold for short-chain fatty acid synthesis and elongation during FASII. This discovery critically expands understanding of ACP function in the *Plasmodium* apicoplast and suggests that ACP has evolved an adaptive role at a key molecular nexus controlling broad organelle metabolism.

### Molecular association of ACP and PKII

Our results reveal that PKII protein levels depend critically on ACP expression, with stringent ACP knockdown resulting in a 75% decrease in PKII abundance (Figure 5B). We detected ACP interaction with PKII in parasites by both proximity biotinylation and immunoprecipitation followed by mass spectrometry (Figure 4). This interaction persisted during heterologous expression in *E. coli* and in the absence of other parasite-specific proteins, suggesting a direct binding interaction between PKII and apicoplast ACP.

We noted that co-purification of ACP with PKII appeared to be sub-stoichiometric in pulldown experiments from bacteria, which may reflect a modest-affinity interaction that partially dissociates under non-equilibrium pulldown and wash conditions. *P. falciparum* apicoplast ACP is known to be modified by a 4-PP group in *E. coli* and to functionally replace bacterial ACP in FASII.^36, 58, 59^ The acylation status of apicoplast ACP expressed in *E. coli* may therefore differ from its native modification status in parasites and impact its heterologous association with PKII in bacteria. It is also possible that ACP and PKII associate as part of a larger metabolic complex in the apicoplast, in which additional interacting proteins tune and tighten the ACP-PKII interaction. We noted that several apicoplast-targeted proteins were identified as ACP interactors by both miniTurbo and IP/MS, suggesting additional interactors of ACP and PKII. These proteins include DXS, which catalyzes the first step in isoprenoid precursor synthesis using PKII-generated pyruvate as a substrate, as well as acyl CoA synthetase 5; falcilysin; and chaperones ClpB1, CPN60, and HSP90 (Supplementary File 1- supplementary table 4). Future biochemical and structural studies can assess the native assembly and potential complex formation of these proteins in parasites.

Two observations support a key role for the 4-PP group in mediating ACP association with PKII. Only wildtype ACP, but not the Ser95Ala mutant (which lacks 4-PP modification), was pulled down with PKII in parasites by IP/MS and in bacterial expression experiments by Ni-NTA/western blot (Figure 4). We also observed that ACPS, the enzyme that attaches 4-PP to ACP (Figure 1), is essential for parasite viability and apicoplast biogenesis (Figure 3). *P. falciparum* ACPS was previously noted to contain a fused metal hydrolase domain of uncertain function,^60^ and future studies will be required to dissect the contribution of this domain to apicoplast disruption in ΔACPS parasites. Nevertheless, our results support the model that ACPS is essential for attaching CoA-derived 4-PP to apicoplast ACP to support critical interaction with PKII. Prior work has shown that the terminal step in CoA synthesis catalyzed by dephospho-CoA kinase (DPCK) occurs in the apicoplast, despite other enzymes in this pathway localizing to the cytoplasm.^16^ The essential role for 4-PP attachment to ACP may provide the evolutionary driving force to localize DPCK for CoA production in the apicoplast.

AlphaFold modeling supports direct association of ACP with the central A domain of PKII, with Ser95 and the 4-PP group of ACP positioned at the binding interface and proximal to the expected PKII active site between the A and B domains (Figure 6 and Figure 6- figure supplement 1). PK enzymes from distinct organisms are known to be allosterically regulated by various small-molecule metabolites.^61^ It is possible that ACP binding allosterically modulates PKII activity, but our results suggest that ACP association directly stabilizes PKII such that ACP knockdown or deletion causes PKII destabilization and loss (Figure 6). This stabilizing role for apicoplast ACP is analogous to similar roles for mitochondrial ACP in binding and stabilizing key mitochondrial complexes via interaction with LYR-motif family proteins,^41, 57^ including in *P. falciparum*.^44^ Apicoplast ACP interaction with PKII is not mediated by an LYR sequence, nor do LYR-motif proteins appear to exist in the apicoplast or in plastids (e.g., chloroplasts) more broadly.^52, 62, 63^ However, bacterial ACPs are known to associate with diverse protein interactors beyond FASII enzymes without evidence for LYR-motif proteins.^53, 54^ The apicoplast ACP-PKII interaction may resemble bacterial ACP interactions and provides a new molecular paradigm for expanded ACP function in eukaryotes. Future studies can clarify if this interaction is a *Plasmodium*-specific adaptation or general to other plastid-containing cells.

**Figure 6.**
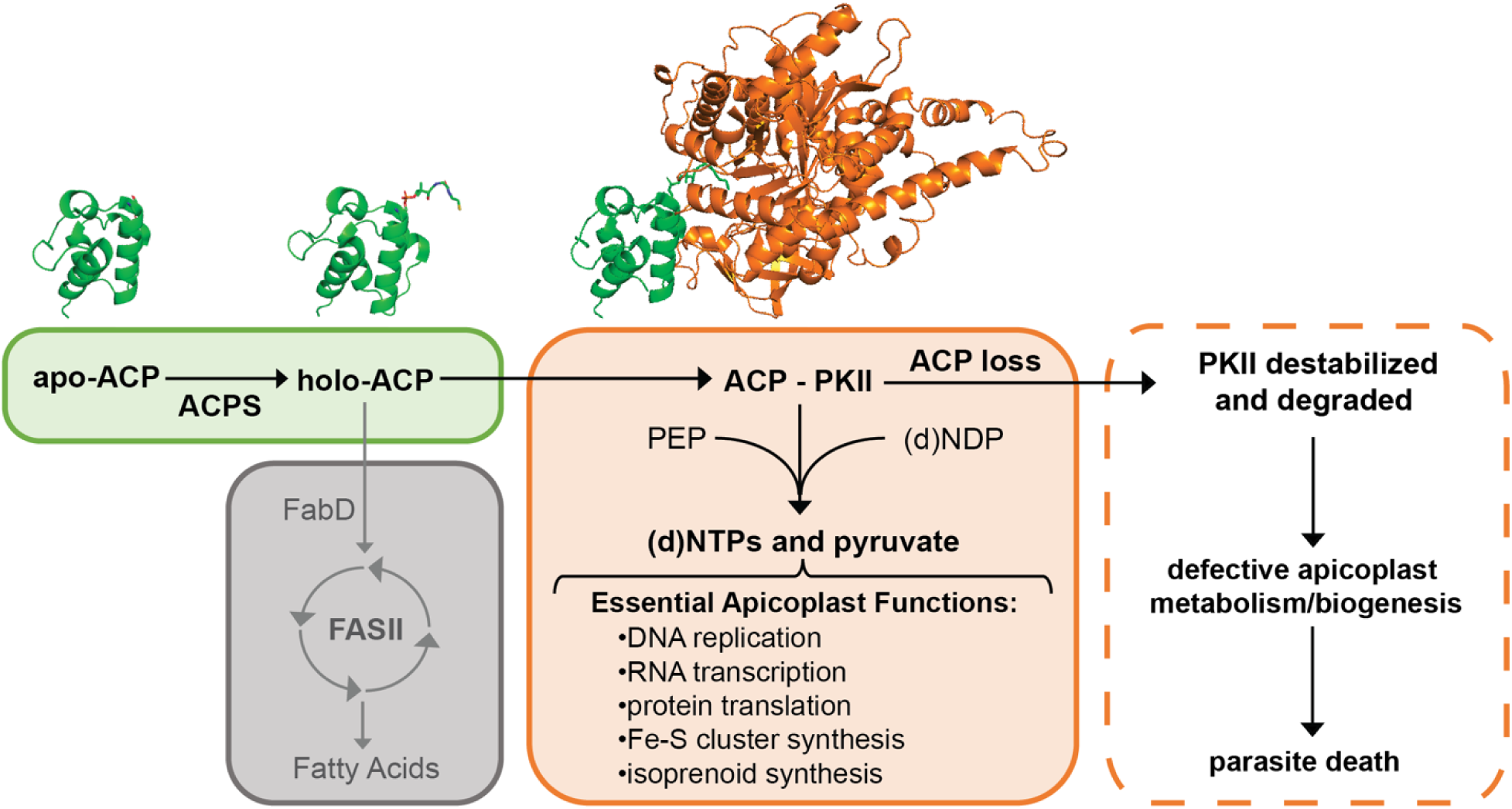
Scheme of ACP interaction with PKII and dysfunctions upon ACP knockdown. Loss of ACPS is expected to phenocopy loss of ACP. PEP = phosphoenolpyruvate, (d)NTPs = nucleoside triphosphates and deoxynucleoside triphosphates. The aACP structural model is based on PDB 3GZM and 5USR. The structural model of aACP bound to PKII was generated using AlphaFold 3. **Figure 6- figure supplement 1**. AlphaFold model of apicoplast ACP bound to pyruvate kinase II.

### Essential ACP function in the apicoplast

The critical stabilizing interaction between ACP and PKII is sufficient to explain essential ACP function in apicoplast biogenesis in *P*. *falciparum*. It is possible that ACP interactions with other apicoplast proteins, including DXS and falcilysin, also contribute to its essential function (e.g., by regulating their stability and/or activity), but we did not further investigate these interactions. PKII is proposed to be the dominant source of pyruvate, NTPs, and dNTPs required for isoprenoid precursor synthesis via the methylerythritol phosphate pathway, RNA transcription, DNA replication, Fe-S cluster synthesis, and protein translation in the apicoplast.^26^ Thus, PKII underpins nearly all aspects of apicoplast function, making it a central hub for organelle metabolism and biogenesis.

In eukaryotes with mitochondrial FASII, mACP association with respiratory complexes via LYR-motif proteins depends on the acylation state of mACP and is thought to serve a regulatory role coupling acetyl-CoA availability to assembly and function of the mitochondrial respiratory chain.^40, 41, 64^ It is intriguing to consider whether apicoplast association of ACP with PKII might similarly depend on ACP modification status and thus serve a regulatory role that modulates PKII levels and broader organelle metabolism under differing growth conditions. The acylation status of apicoplast ACP and its 4-PP group in blood-stage parasites has not been directly determined. With little to no detectable FASII activity, ACP may predominate in its holo, uncharged form with a free thiol on the 4-PP group (Figure 1), as suggested by prior experiments testing the impact of redox cycling on ACP electrophoretic mobility.^36^

It is possible that a small and variable population exists of acyl-ACP bearing host-derived fatty acids, which can occur if ACPS uses an acyl-CoA (bearing a scavenged fatty acid) to attach an acyl-PP group to ACP.^65^ *P. falciparum* ACP has also been reported to have self-acylation activity via nucleophilic reaction with an acyl-CoA, although the very low rate constant (*k*_cat_ ∼0.1 min^-1^) reported for this activity makes its biological relevance questionable.^66^ We emphasize, however, that future experiments will be required to rigorously determine the modification states of apicoplast ACP, including tandem mass spectrometry experiments that can enrich, eject, and fragment the 4-PP group to interrogate its structure.^67^ Based on a model in which ACP exists predominantly in the holo, uncharged state in blood-stage parasites, we speculate that an increasing population of acyl-ACP might tighten interaction with PKII and thus increase overall PKII protein stability and activity. Variable ACP acylation and impacts on PKII activity might play a more critical role during parasite growth within mosquitoes or human hepatocytes, when FASII activity is upregulated to increase the pool of acyl-ACP and when parasites are undergoing large-scale proliferation with enhanced reliance on apicoplast biosynthetic functions.^14, 15, 68^

### Implications for other apicomplexan parasites

*Toxoplasma gondii*, like other apicomplexan parasites, retains an apicoplast FASII pathway that is dispensable in tachyzoites but enhances their fitness and growth.^69, 70^ Knockdown of apicoplast ACP in *T. gondii*, however, produces a much more severe phenotype with strong growth impairment.^71^ This observation suggests that ACP may also plays additional roles in *Toxoplasma* beyond FASII^69^ that may also involve stabilizing interactions with diverse organelle proteins. However, apicoplast PKII is dispensable for *Toxoplasma* tachyzoites,^72^ suggesting that the specific interactions that underpin ACP essentiality in *T. gondii* may differ from *P. falciparum*. It remains unclear whether any FASII-independent functions for ACP in *Toxoplasma* depend on 4-PP attachment, but genome-wide CRISPR screens in tachyzoites suggest a critical role for ACPS.^73^ Prior work in bacteria has suggested that ACPS is a viable therapeutic target,^74, 75^ and inhibiting this essential enzyme may be a potent antimalarial strategy with efficacy against *P. falciparum* as well as other apicomplexan parasites.

## MATERIALS AND METHODS

### Key Resources Table

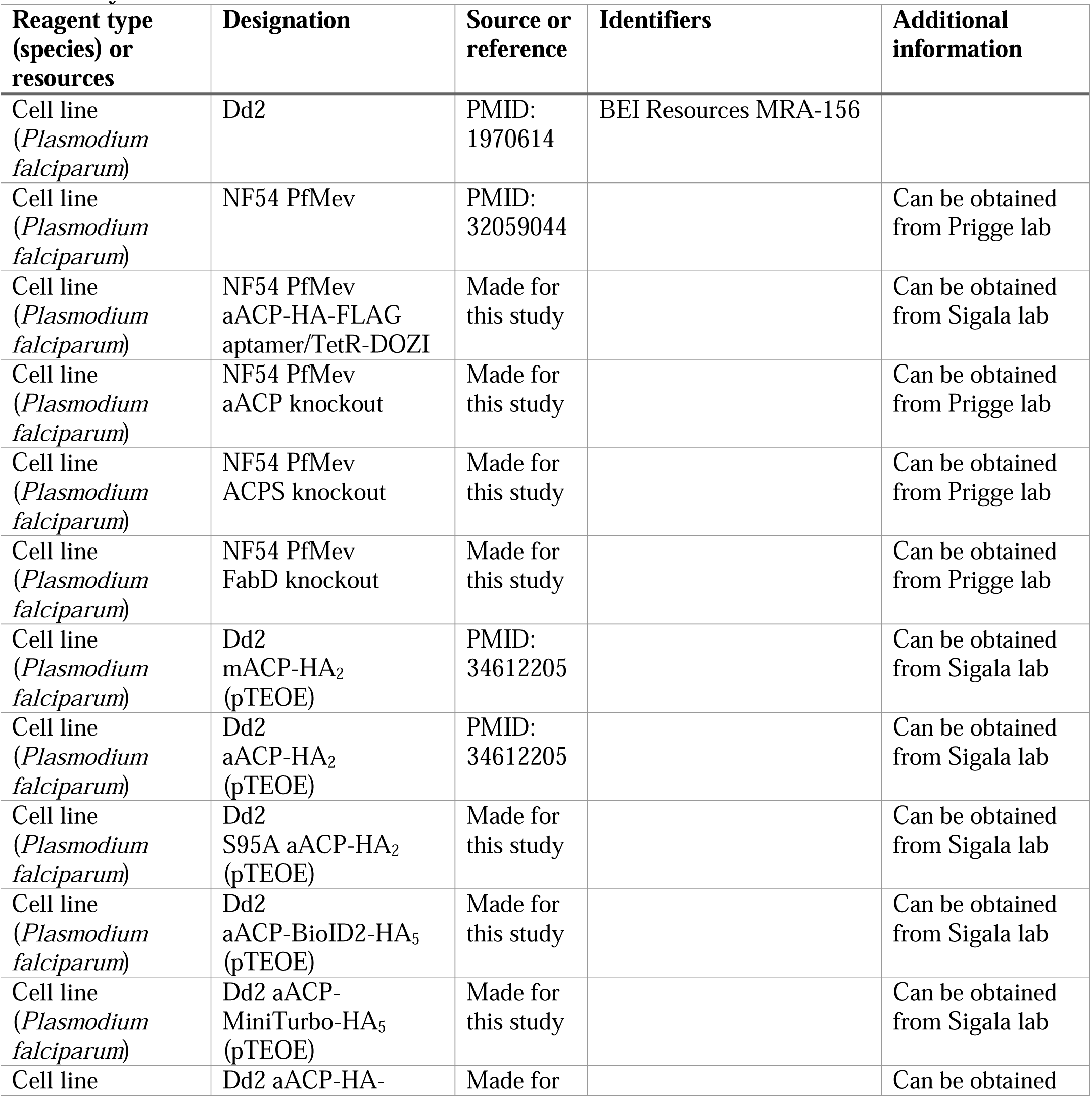
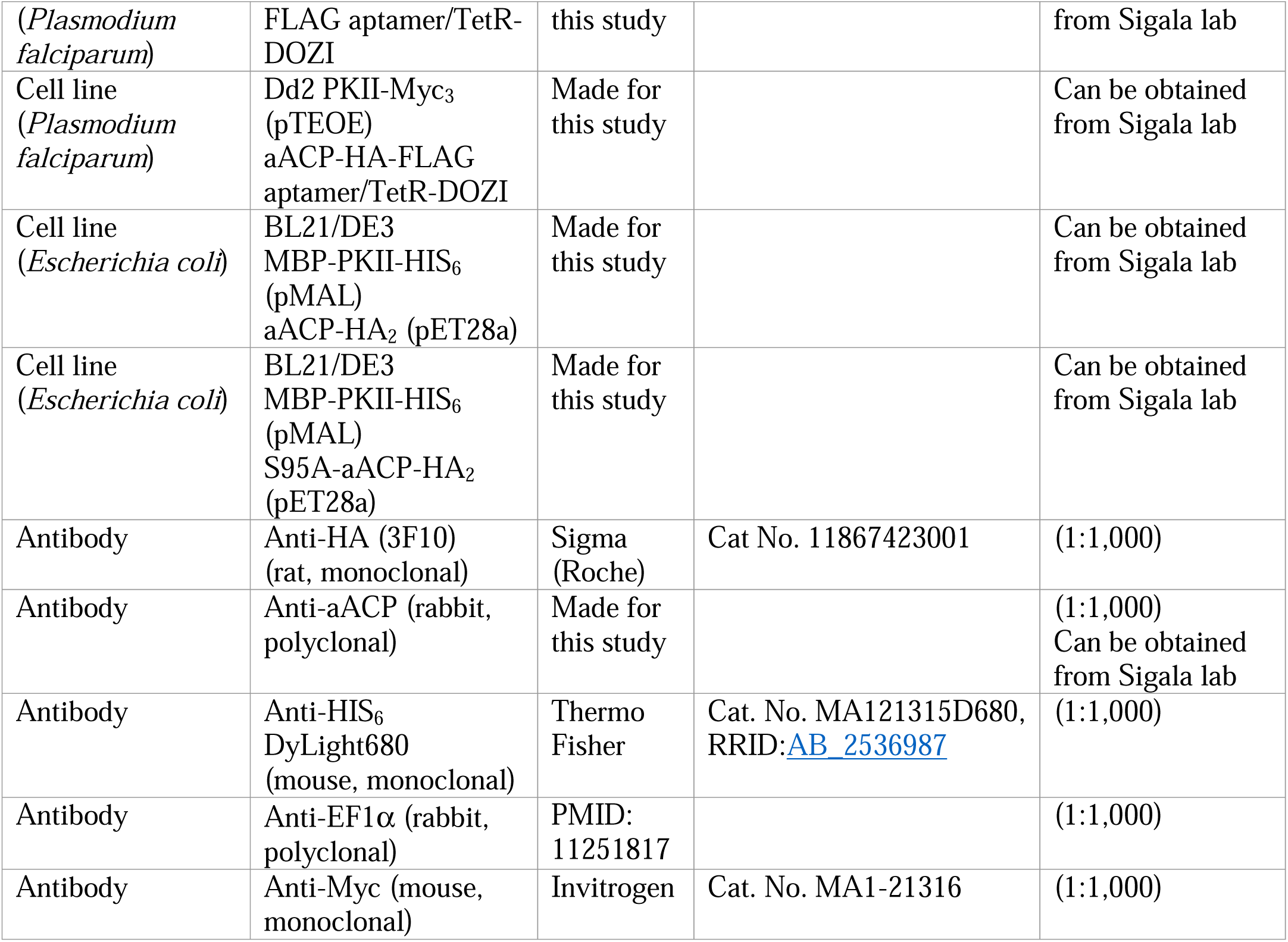

### Parasite Culturingh

All experiments were performed using *P. falciparum* Dd2^76^ or NF54 PfMev^46^ parasite strains. Parasite strains were confirmed based on expected drug resistance and were *Mycoplasma*-free by PCR. Parasite culturing was performed in Roswell Park Memorial Institute medium (RPMI-1640, Thermo Fisher 23400021) supplemented with 2.5 g/L Albumax I Lipid-Rich BSA (Thermo Fisher 11020039), 15 mg/L hypoxanthine (Sigma H9636), 110 mg/L sodium pyruvate (Sigma P5280), 1.19 g/L HEPES (Sigma H4034), 2.52 g/L sodium bicarbonate (Sigma S5761), 2 g/L glucose (Sigma G7021), and 10 mg/L gentamicin (Invitrogen Life Technologies 15750060). Cultures were generally maintained at 2% hematocrit in deidentified human erythrocytes obtained from the University of Utah Hospital blood bank, at 37°C, and at 5% O_2_, 5% CO_2_, 90% N_2_.

### Cloning and transfection for episomal expression in *P. falciparum*

The genes encoding apicoplast ACP (aACP, PF3D7_0208500) and mitochondrial ACP (mACP, PF3D7_1208300) were cloned into the pTEOE episomal protein expression vector with C-terminal HA_2_ tag as described previously.^44^ The apicoplast ACP S95A mutant in the pTEOE vector was made by site-directed mutagenesis of the prior aACP pTEOE plasmid using primers P1/P2 (primer sequences in Supplemental File 1- supplementary table 5) and confirmed by Sanger sequencing using plasmid-specific primers. For cloning of aACP-BioID2-HA_5_/pTEOE and aACP-MiniTurbo-HA_5_/pTEOE plasmids, aACP was PCR-amplified from the aACP-HA_2_/pTEOE plasmid using primers P3-P5 to yield aACP amplicons with 5’ homology to the pTEOE vector and 3’ homology to BioID2 or miniTurbo. BioID2 and miniTurbo were PCR-amplified from previously published plasmids^77^ using primers P6-P9 to produce amplicons with 5’ homology to aACP and 3’ homology to the HA_5_ tag sequence in a pTEOE vector. The aACP amplicon was combined with either the BioID2 or miniTurbo amplicon in a ligation-independent cloning reaction (NEBuilder HiFi DNA Assembly Master Mix) for insertion into pTEOE. Pyruvate kinase II (PKII, PF3D7_1037100) was PCR-amplified from Dd2 parasite gDNA via primers P10/P11 and combined with a 3xMyc amplicon created with primers P12/P13 in a ligation-independent cloning reaction for insertion into pTEOE at XhoI/NotI sites to create a PKII-Myc_3_/pTEOE plasmid. Correct clonal plasmid sequences were confirmed by Sanger and/or whole-plasmid sequencing (plasmidsaurus.com).

For episomal expression using the pTEOE vector, Dd2 parasite-infected erythrocytes were transfected in 1x cytomix containing 50-100 µg purified plasmids and 25 µg of the pHTH transposase plasmid^78^ by electroporation in 0.2 cm cuvettes using a Bio-Rad Gene Pulser Xcell system (0.31 kV, 925 µF). Transfected cultures were allowed to expand in the absence of drug for 48 hr and then selected in 5 nM WR99210 (Jacobus Pharmaceuticals). Stable drug-resistant parasites returned from transfection in 2-8 weeks and were used as polyclonal cultures.

### Cloning and transfection for CRISPR/Cas9 gene editing for aACP knockdown

CRISPR/Cas9-stimulated repair by double-crossover homologous recombination was used to tag the aACP gene (Pf3D7_0208500) to encode a C-terminal HA-FLAG epitope tag fusion and the 3’ 10X aptamer/TetR-DOZI system^47^ to enable regulated aACP expression using anhydrotetracycline (aTc, Caymen Chemicals 10009542). A guide RNA sequence of AGACCTCGTTGAATTAATTA corresponding to the sense strand in the third exon of the aACP gene was cloned by ligation independent methods using primer pairs P14/P15 into a modified version of the pAIO CRISPR/Cas9 vector that contained a HindIII site in place of the BtgZI site. For homology-directed repair to tag the aACP gene, the donor pMG75 repair plasmid was created using ligation-independent cloning to insert a gBlock gene fragment from IDT into the unique AscI/AatII cloning sites. This gBlock contained 153 bp of the 3’ untranslated region of the aACP gene (starting at position 95 downstream from the TAA stop codon), an AfeI site, and the 316 bp of the 3’ end of the aACP coding sequence (excluding the 175 bp intron 2). The gBlock sequence included two underlined shield mutations within the targeted gRNA sequence (AGA<u>T</u>CTCGTTGAATTAAT<u>C</u>A) that resulted in silent mutations. All cloned sequences were verified by Sanger sequencing. Before transfection, the pMG75 vector was linearized by AfeI digestion performed overnight at 37° C.

Dd2 and PfMev parasites were transfected by electroporation of uninfected erythrocytes with 50 µg each of the linearized pMG75 donor plasmid and pAIO Cas9 plasmid in 1x cytomix using the Bio-Rad Gene Pulser Xcell system (0.31 kV, 950 µF) and 0.2 cm cuvettes. These transfected erythrocytes were then mixed with 1-2% parasitemia synchronous schizont-stage parasites, maintained in 0.5 µM aTc, and allowed to grow for 72 hours prior to drug selection with 6 µM blasticidin-S (Invitrogen R21001). Polyclonal parasites that returned from transfection were genotyped by genomic PCR using primers P16/P17 (wildtype) and P16/P18 (integrated). Integrated PfMev parasites were fully integrated without evidence for unmodified aACP locus and used as polyclonal parasites in downstream assays. Dd2 parasites were cloned by limiting dilution. Integrated aACP-aptamer/TetR-DOZI Dd2 parasites were transfected by electroporation with 50 µg of the PKII-Myc_3_/pTEOE plasmid as above, selected with 5 nM WR99210 (Jacobus Pharmaceuticals), and used as polyclonal parasites for subsequent experiments.

### Cloning and transfection for knockout of aACP, ACPS, and FabD

The genes encoding apicoplast ACP (Pf3D7_0208500), holo-ACP synthase (ACPS, Pf3D7_0420200), and FabD (Pf3D7_1312000) were disrupted in the NF54 PfMev line using CRISPR/Cas9 and gene deletion by double-crossover homologous recombination. Homology arm regions were PCR-amplified from genomic DNA with primers P19-P22 (aACP: 421 bp 5’ arm, 362 bp 3’ arm), P23-P26 (ACPS: 416 bp 5’ arm, 426 bp 3’ arm), and P27-P30 (FabD: 346 bp 5’ arm, 402 bp 3’ arm) and cloned into vector pRS^46^ using ligation-independent cloning (Clontech In-Fusion). DNA encoding guide RNA with sequences AATGAACTCAAATTTTACCA (aACP), TAAAATATTGAACCCACGAG (ACPS), and GATAGAAAATTTGGTTTATG (FabD) were cloned into a modified pAIO vector called pCasG^79^ using primers P31/P32, P33/P34, P35/P36, respectively.

For transfection, 75 µg of each pRS plasmid was combined with 75 µg of the matching pCasG plasmid and electroporated into NF54 PfMev parasites. Transfected parasites were allowed to expand for 48 hr in 50 µM mevalonate before selection with 5 nM WR99210 and 50 µM DL-mevalonolactone (Cayman Chemicals 20348, referred to as mevalonate in this work). Parasites returning from positive selection were genotyped by PCR using primers P37-P40 (aACP), P41-P44 (ACPS), P45-P48 (FabD) in conjunction with primers P49-50 to confirm complete on-target integration in polyclonal cultures. A primer scheme and expected amplicon sizes are provided in Supplementary File 2. Apicoplast (SufB: Pf3D7_API04700), mitochondrion (CoxI, Pf3D7_MIT02100), and nuclear (LDH, Pf3D7_1324900) genome PCR was performed using primers P51-P56 to confirm apicoplast status in knockout parasites.

### Cloning for recombinant expression in *E. coli*

The pMAL expression vector encoding codon-harmonized mature (AA42-745) PKII with N-terminal maltose binding protein (MBP) domain, TEV protease site, and His_6_ tag was cloned as described previously.^26^ This plasmid was co-transformed with either a WT (Δ2-40) aACP-HA_2_ pET28a plasmid published previously^44^ or a S95A (Δ2-40) aACP-HA_2_ pET28a plasmid prepared from the WT aACP plasmid by site-directed mutagenesis using primers P1/P2 and confirmed by Sanger sequencing.

### Recombinant expression in *E. coli*

Bacterial expression experiments were performed as previously described.^44^ Plasmids for co-expression of His_6_-PKII with either WT or S95A aACP-HA_2_ were transformed into chemically competent BL21/DE3 cells and selected with kanamycin (50 µg/mL) and carbenicillin (100 µg/mL) for inducible expression with 1 mM isopropyl 1-thio-β-galactopyranoside (IPTG) (Goldbio 367931) in 20 mL LB media. Cells were grown at 37° C until reaching an optical density (600 nm) of 0.6, induced with IPTG, and grown overnight at 20° C before harvest by centrifugation. Bacterial pellets were resuspended in 1 mL cold phosphate buffered saline (pH 7.4), lysed by sonication on ice using a Branson sonicator equipped with a microtip, and clarified by centrifugation at 17,000 x g in a Fisher Scientific accuSpin 17R microcentrifuge. Clarified lysates were treated with recombinant TEV protease overnight at 4° C to cleave off the MBP tag and release His_6_-PKII. Treated lysates were then incubated with 100 µL of Ni-NTA resin (Thermo Scientific 88221), after resin equilibration with buffer A (50 mM NaH_2_PO_4_, 500 mM NaCl, 5 mM imidazole, pH 8.0) to bind His_6_-PKII and any co-associating proteins. Resin was collected by centrifugation, washed three times with buffer A, and eluted by incubation with 100 µL buffer B (buffer A + 500 mM imidazole) for 15 min. on ice. Eluates were diluted into SDS sample buffer and further analyzed by SDS-PAGE and western blot.

### Parasite Growth Assays

Parasites were synchronized with 5% (w/v) D-sorbitol for 10 min at room temperature, washed 3x with drug-free RPMI medium, and then seeded for growth assays at an initial ring parasitemia of ∼0.25% with daily media changes and parasitemia measurements. For tests of regulatable aACP expression, Dd2 aACP-HA-Flag 10xAptamer/TetR-DOZI parasites were split into biological triplicate populations either grown without aTc (-aTc), with 200 µM isopentyl pyrophosphate (IPP, Isoprenoids IPP001) added (-aTc, +IPP), or with 0.5 µM aTc (+aTc). Parasitemia was measured daily in technical duplicate 10 µL samples, diluted into 200 µL of 1 µg/mL acridine orange (Invitrogen Life Technologies A3568) in phosphate buffered saline (PBS) and analyzed by flow cytometry using a BD FACSCelesta system monitoring SSC-A, FSC-A, PE-A, FITC-A, and PerCP-Cy5-5-A channels. Averages ±SD were determined and plotted using GraphPad Prism 9.0.

Synchronous growth assays of ΔACP, ΔACPS, and ΔFabD PfMev parasites were performed similarly, with growth monitored ±50 µM mevalonate in cultures. For growth assays in low-lipid conditions, parasites and RBCs were washed into modified complete RPMI medium in which Albumax I was replaced with fatty acid-free bovine serum albumin (GoldBio A-421-10) supplemented with 45 µM oleic (Sigma O1008) and 30 µM palmitic (Sigma 57-10-3) acids with daily media changes. Parasites were cultured ≥1 week in low-lipid conditions prior to initiating synchronous growth assays, which were completed in biological triplicate samples.

### Production and validation of custom anti-apicoplast ACP antibody

The sequence encoding mature, processed (Δ1-40) apicoplast ACP was cloned into pET28 with an N-terminal His-tag, expressed in *E. coli*, purified by Ni-NTA, cleaved by thrombin, and purified by FPLC using previously reported methods.^44^ Purified protein was then injected into a rabbit for polyclonal antibody production by Cocalico Biologicals Inc (https://cocalicobiologicals.com) following their standard protocol. Rabbit serum was validated for specific detection of apicoplast ACP with recombinant protein expressed in bacteria (Figure 2- figure supplement 5).

### Fluorescence Microscopy

Live-parasite microscopy and immunofluorescence assays (IFA) with fixed parasites were performed as previously described.^80^ For live-cell experiments, parasite nuclei were visualized by incubation with 1-2 µg/mL Hoechst 33342 (Pierce 62249) for 10-20 min at room temperature. The parasite apicoplast was visualized in live PfMev parasites using the endogenous fluorescence of ACP_L_-GFP expressed in this line.^46^ For IFA experiments, parasites were fixed, stained, and mounted as described previously.^22, 81^ The nucleus was visualized in IFA samples using ProLong Gold Antifade Mountant with DAPI (Invitrogen P36931). The apicoplast was visualized using polyclonal rabbit anti-aACP antibodies either previously published^36^ or produced for this study and a goat anti-rabbit 2° antibody (Invitrogen A32731). Images of aACP knockdown parasites (Dd2 and PfMev) were taken on an EVOS M5000 imaging system, with processing done in Fiji/ImageJ. Images of ΔACPS, ΔACP, and ΔFabD parasites were acquired on a Zeiss Axioimager M2 microscope, with process done with VOLOCITY (PerkinElmer). All image processing was done on a linear scale.

### SDS-PAGE and western blots

For analysis of *P. falciparum* samples, blood-stage parasites were treated with 0.05% saponin (Sigma 84510) in PBS for 5 min at room temperature and harvested by centrifugation. Saponin pellets were then lysed by resuspension in 1% Triton X-100 in PBS after addition of protease inhibitors (Invitrogen A32955) and sonicated ∼20 pulses (50% duty cycle, 30-40% power) on a Branson microtip sonicator. Parasites lysates were clarified by centrifugation at 17,000 x g for 10 min. Bacterial samples were harvested and lysed as described above. Protein concentrations in lysates were quantified by Lowry colorimetry, and 50 µg of total protein from each sample was mixed with SDS sample buffer and heated at 95° C for 10 min. Samples were loaded and fractionated by 10-16% SDS-PAGE in Tris-HCl buffer, transferred to nitrocellulose membrane using the Bio-Rad wet-transfer system for 1 hr at 100 V, and blocked with 5% non-fat milk in TBS-T (50 mM Tris pH 7.4, 150 µM NaCl, 0.5% Tween-20). Membranes were probed with primary antibodies: rat anti-HA-tag monoclonal 3F10 (Roche), rabbit anti-EF1α or rabbit anti-EF1β^82^, mouse anti-Myc-tag monoclonal (Invitrogen MA1-21316), and/or mouse anti-His-tag monoclonal-HRP (QIAGEN 34460). Membranes were washed 3x in TBS-T buffer and probed with secondary antibodies: donkey anti-rabbit IRDye680 (LiCor 926-68023), goat anti-rat IRDye800CW (Licor 926-32219), goat anti-rat-HRP (Invitrogen 62-9520), or goat anti-mouse-HRP (Invitrogen A28177). Membranes were imaged on a Licor Odyssey CLx system or Invitrogen iBright 1500 imager. Band intensities were quantified by densitometry using ImageJ and analyzed with GraphPad Prism 9.0.

### Measuring DNA and RNA abundance in parasites by PCR

Apicoplast ACP knockdown Dd2 or PfMev parasites were synchronized with 5% D-sorbitol and cultured +aTc, −aTc/+IPP (Dd2), or −aTc/+Mev (PfMev) for 36 or 84 hours prior to harvest for DNA and RNA extraction, as previously reported.^22^ Parasite DNA was extracted using a QIAamp DNA Blood Mini kit (QIAGEN 51106). Parasite RNA was purified by Trizol (Invitrogen 15596026) and phenol-chloroform extraction. RNA was reverse transcribed to cDNA using a SuperScript IV VILO RT kit (Invitrogen 11766050) using gene-specific primers, as published previously.^22^ Invitrogen Quantstudio Real-Time PCR systems were used to quantify abundance of DNA and cDNA using SYBR green dye. The relative DNA or cDNA abundance of each apicoplast gene was normalized to the average of three nuclear-encoded genes for each samples, and −aTc conditions were compared to +aTc conditions at each time-point by the comparative Ct method.^83^

All qPCR experiments were performed in biological triplicate, with each sample analyzed in technical duplicate, and data were analyzed by unpaired Student’s t-test in GraphPad Prism 9.0. PCR analysis of apicoplast, mitochondrial, and nuclear genomes status was performed in ΔACP, ΔACPS, and ΔFabD PfMev parasites, as previously reported.^46^

### Immunoprecipitations

Immunoprecipitation (IP) experiments for mass spectrometry analysis were performed for Dd2 parasites episomally expressing WT or S95A apicoplast ACP-HA_2_ or WT mitochondrial ACP-HA_2_, as previously described.^44^ Blood-stage parasites from ∼50 mL cultures were harvested by centrifugation, treated with 0.05% saponin (Sigma 84510) in PBS for 5 min at room temperature, and clarified by centrifugation at 3000 x g for 30 min at 4°C. IP was performed using Pierce anti-HA magnetic beads (Thermo Scientific 88836). Parasite pellets were lysed in 1 mL cold 1% Triton (Sigma 9002931)/PBS, 1% digitonin (GoldBio 11024-24-1)/PBS, or RIPA buffer (150 mM NaCl, 1% Triton X-100, 0.5% deoxycholate, 0.1% SDS, 50 mM Tris, pH 8.0) plus protease inhibitor (Thermo Scientific A32955). Pellets were dispersed by brief sonication on a Branson sonicator equipped with a microtip probe and then incubated at 4°C for 1 hr on a rotator. Lysates were clarified by centrifugation at 17,000 x g for 10 min 30 μL of anti-HA magnetic beads was equilibrated in 170 μL of cold 1× Tris-buffered saline (20 mM Tris, pH 7.6, 150 mM NaCl) + 0.05% Tween-20 (TBS-T), and the beads were collected on a magnetic stand. Parasite lysates supernatants (1 mL) were added to the equilibrated beads and incubated for 1 hr at 4°C by rotation. Beads were washed three times with ice-cold 1× TBS-T and transferred to a new tube. Bound proteins were eluted with 100 μL 8 M urea (in 100 mM Tris at pH 8.8) and precipitated by adding 100% trichloroacetic acid (Sigma 76039) to a final concentration of 20%. Samples were incubated on ice for 1 hr and centrifuged at 17,000 x g for 25 min at 4° C. Supernatants were removed by vacuum aspiration, and protein pellets were washed once with 500 μL of cold acetone. The protein pellets were air-dried for 30 min and stored at –20°C.

### Proximity biotinylation experiments

For proximity biotinylation experiments with Dd2 parasites episomally expressing aACP-BioID2 or aACP-miniTurbo, 50 mL cultures were synchronized with 5% D-sorbitol and grown to ∼15% parasitemia before treatment with 50 µM biotin (Sigma 58-85-5) for 24 hr before harvesting by centrifugation and saponin treatment. Parental Dd2 parasites were used as negative control and treated identically. Parasite pellets were lysed in 1 mL 1% Triton X-100/PBS and clarified by centrifugation at 17,000 x g. Clarified lysates were added to 25 µL of M-280 streptavidin magnetic beads (Invitrogen 11205D) that had been equilibrated in PBS, incubated overnight with rotating at 4° C, and washed 5x with 1 mL 1% Triton/PBS, 4x with 1 mL 8M urea/PBS, and 2x in PBS ensuring that samples were moved to new tubes between each wash condition. The final washed beads were resuspended in 50 µL PBS and stored at 4° C briefly before shipment for on-bead tryptic digest and analysis by tandem mass spectrometry. 5 µL of the final suspended beads were diluted into SDS sample buffer, boiled at 95° C for 10 min, fractionated by 12% SDS-PAGE, transferred to nitrocellulose, and probed with streptavidin-AF680 conjugate (Invitrogen S32358).

### Mass spectrometry

Protein samples were reduced and alkylated using 5 mM Tris (2-carboxyethyl) phosphine and 10 mM iodoacetamide, respectively, and then enzymatically digested by sequential addition of trypsin and lys-C proteases, as previously described.^84^ The digested peptides were desalted using Pierce C18 tips (Thermo Fisher Scientific), dried, and resuspended in 5% formic acid. Approximately 1 μg of digested peptides was loaded onto a 25-cm-long, 75-μm inner diameter fused silica capillary packed in-house with bulk ReproSil-Pur 120 C18-AQ particles, as described previously.^85^ The 140-min water-acetonitrile gradient was delivered using a Dionex Ultimate 3,000 ultra-high performance liquid chromatography system (Thermo Fisher Scientific) at a flow rate of 200 nL/min (Buffer A: water with 3% DMSO and 0.1% formic acid, and Buffer B: acetonitrile with 3% DMSO and 0.1% formic acid). Eluted peptides were ionized by the application of distal 2.2 kV and introduced into the Orbitrap Fusion Lumos mass spectrometer (Thermo Fisher Scientific) and analyzed by tandem mass spectrometry. Data were acquired using a Data-Dependent Acquisition method consisting of a full MS1 scan (resolution = 120,000) followed by sequential MS2 scans (resolution = 15,000) for the remainder of the 3-s cycle time. Data were analyzed using the Integrated Proteomics Pipeline 2 (Integrated Proteomics Applications, San Diego, CA). Data were searched against the protein database from *P. falciparum* 3D7 downloaded from UniprotKB (10,826 entries) on October 2013. Tandem mass spectrometry spectra were searched using the ProLuCID algorithm followed by filtering of peptide-to-spectrum matches by DTASelect using a decoy database-estimated false discovery rate of <1%. The proteomics data are deposited in the MassIVE data repository (https://massive.ucsd.edu) under the identifier MSV000100931. Protein enrichment values were calculated as the log_2_ ratio of spectral counts or intensity values for the indicated protein in the experimental versus negative-control datasets (mini-Turbo:Dd2, BioID2:Dd2, aACP:mACP, WT aACP:S95A ACP). Protein spectral count values of zero were arbitrarily converted to one for purposes of calculating a log_2_ enrichment ratio.

### Structural modeling

A structural model of mature *P. falciparum* apicoplast ACP bound to mature PKII was generated using AlphaFold 3^86^ and visualized using PyMOL 3.1^87^.

## Materials availability

All materials created during this study can be obtained by contacting the Sigala or Prigge labs.

## Data availability

The proteomics data are deposited in the MassIVE data repository (https://massive.ucsd.edu) under the identifier MSV000100931. All data generated or analyzed during this study are included in the manuscript and supporting files. Source data files have been provided for Figures 2-5. Figure 4- source data 2 contains the full list of proteins identified by mass spectrometry experiments. Supplementary file 1 contains supplementary tables 1-4 (apicoplast-targeted proteins identified in mass spectrometry experiments) and supplementary table 5 (PCR primer sequences). Supplementary file 2 shows the primer scheme and amplicon sizes for PCR analyses of gene deletions.

## ACKNOWLEDGEMENTS

We thank Jared Rutter, Mike Burkart, Dennis Winge and members of the Sigala lab for helpful discussions and thank Amanda Mixon Blackwell for assistance with qPCR experiments. This work was funded by NIH grants R35GM161383 (to PAS), R35GM153408 (to JAW), and R01AI125534 (to STP) and a pilot award (to PAS) from the Utah Center for Iron and Heme Disorders (funded by U54DK110858). STP was supported by Bloomberg Philanthropies and the Johns Hopkins Malaria Research Institute. MO was supported in part by NIH grant T32AI055434. JPA was supported in part by a predoctoral fellowship from the American Heart Association (24PRE1193935). Microscopy, DNA synthesis and sequencing, and flow cytometry were performed using core facilities at the University of Utah.

## Figure Supplements

**Figure 2- figure supplement 1.**
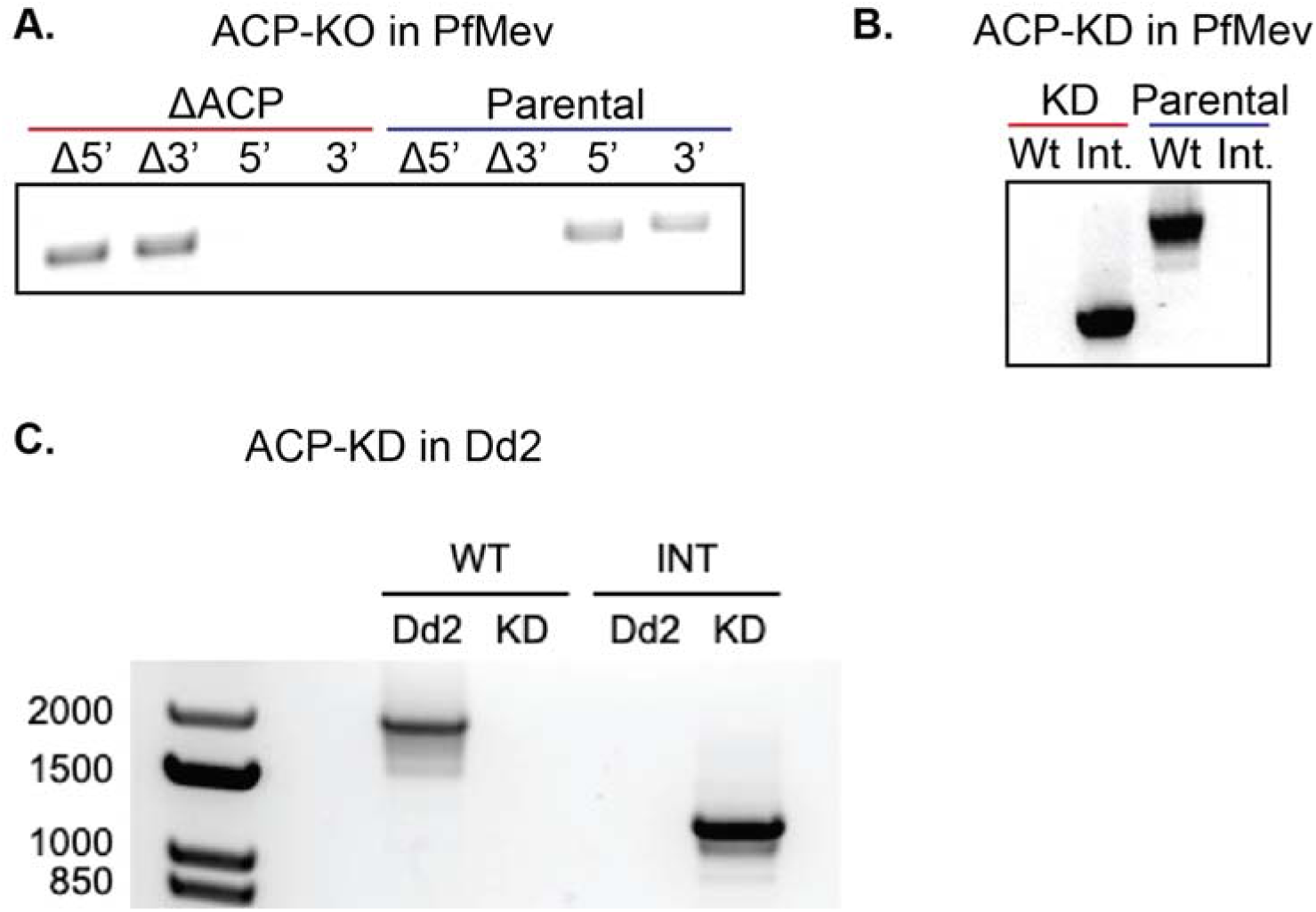
Integration PCRs for deletion or modification of the apicoplast ACP gene. (**A**) Genotyping PCR for successful deletion of the apicoplast ACP gene (Pf3D7_0208500) in polyclonal PfMev parasites showing selective amplification of sequence at the 5’ and 3’ ends of the drug-resistance marker (Δ5’ and Δ3’) in knockout parasites but not in parental PfMev parasites and exclusive amplification of sequence at the 5’ and 3’ ends of the ACP CDS in WT but not ΔACP parasites. Genotype PCR analysis confirming on-target integration of the 3’ aptamer/TetR-DOZI cassette at the apicoplast ACP locus in polyclonal PfMev (**B**) or clonal Dd2 parasites (**C**) and the absence of the unmodified wildtype (WT) locus, with parental parasites serving as negative control for integration.

**Figure 2- figure supplement 2.**
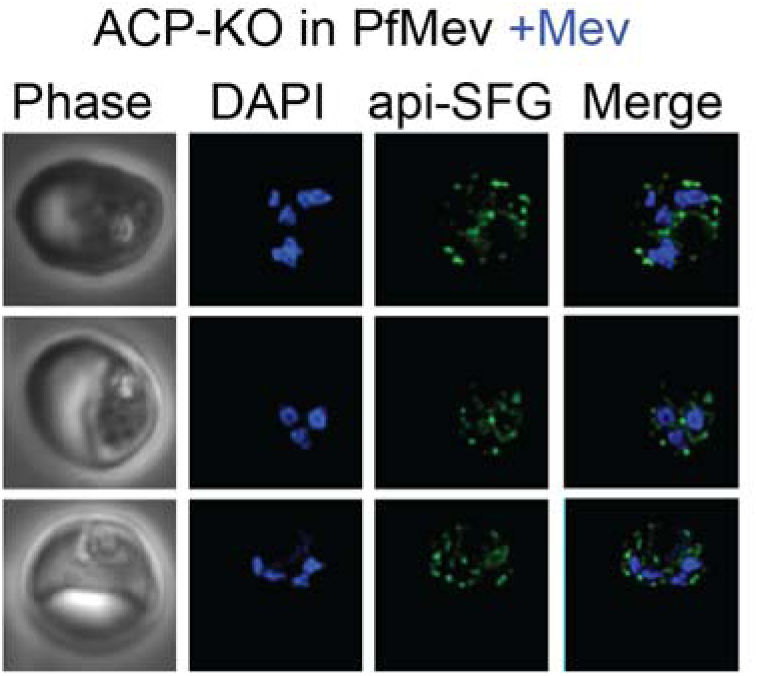
Additional microscopy images ΔACP PfMev NF54 parasites. Phase = phase contrast images, api-SFG = ACP_L_-superfolder GFP, DAPI = nuclear DNA stain.

**Figure 2- figure supplement 3.**
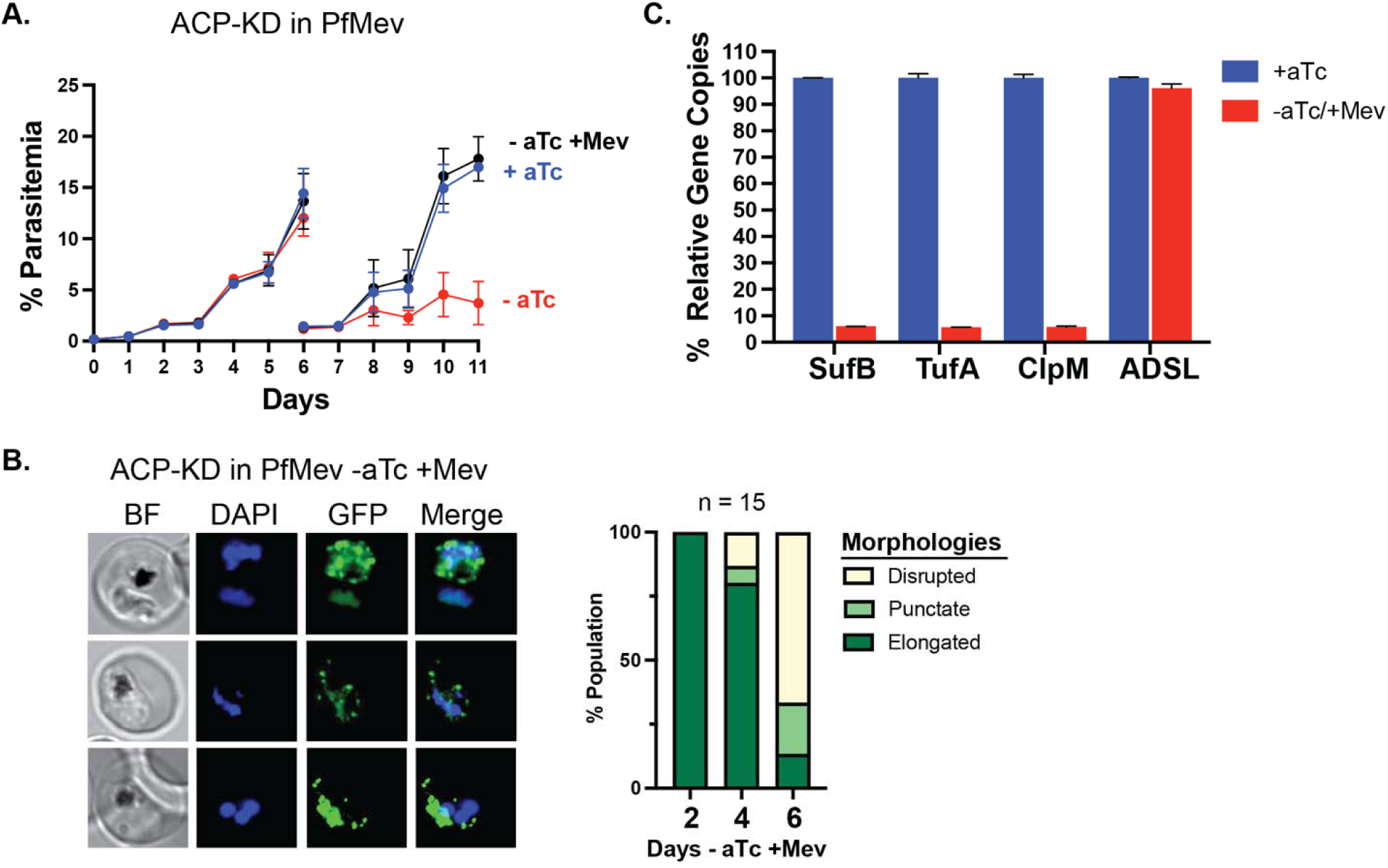
Synchronous growth assays (**A**), fluorescence microscopy (**B**), and genomic qPCR (**C**) of ACP knockdown in PfMev NF54 parasites. For the growth assay in panel A, cultures were split on day 6. Samples for genomic qPCR were cultured as indicated for ≥8 days. Gene copy numbers were based on amplification of apicoplast (SufB: Pf3D7_API04700, ClpM: Pf3D7_API03600, TufA: Pf3D7_API02900) or nuclear (ADSL: Pf3D7_0206700) relative to nuclear (STL: Pf3D7_0717700, I5P: Pf3D7_0802500) genes. Indicated qPCR ratios were normalized to +aTc and are the average ± SD of biological triplicates.

**Figure 2- figure supplement 4.**
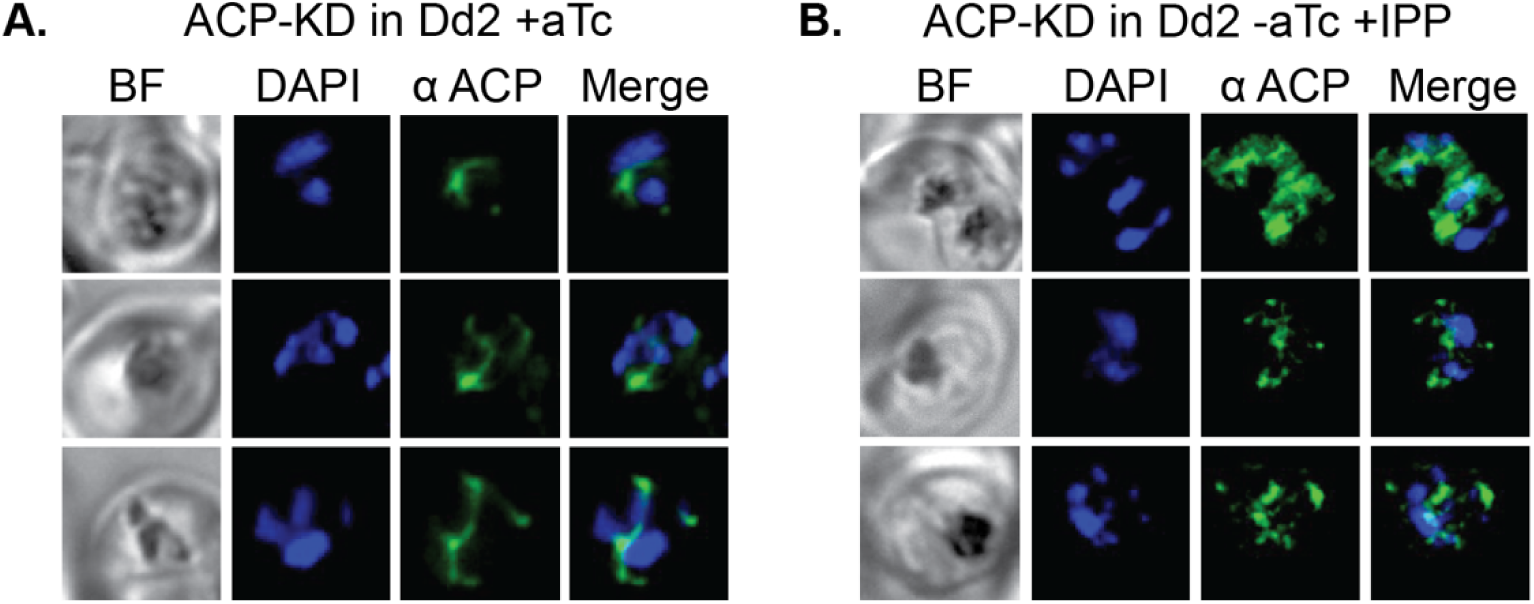
Additional IFA images for apicoplast morphology upon ACP knockdown in Dd2 parasites in +aTc (**A**) or −aTc/+IPP (**B**) conditions. BF = bright field.

**Figure 2- figure supplement 5.**
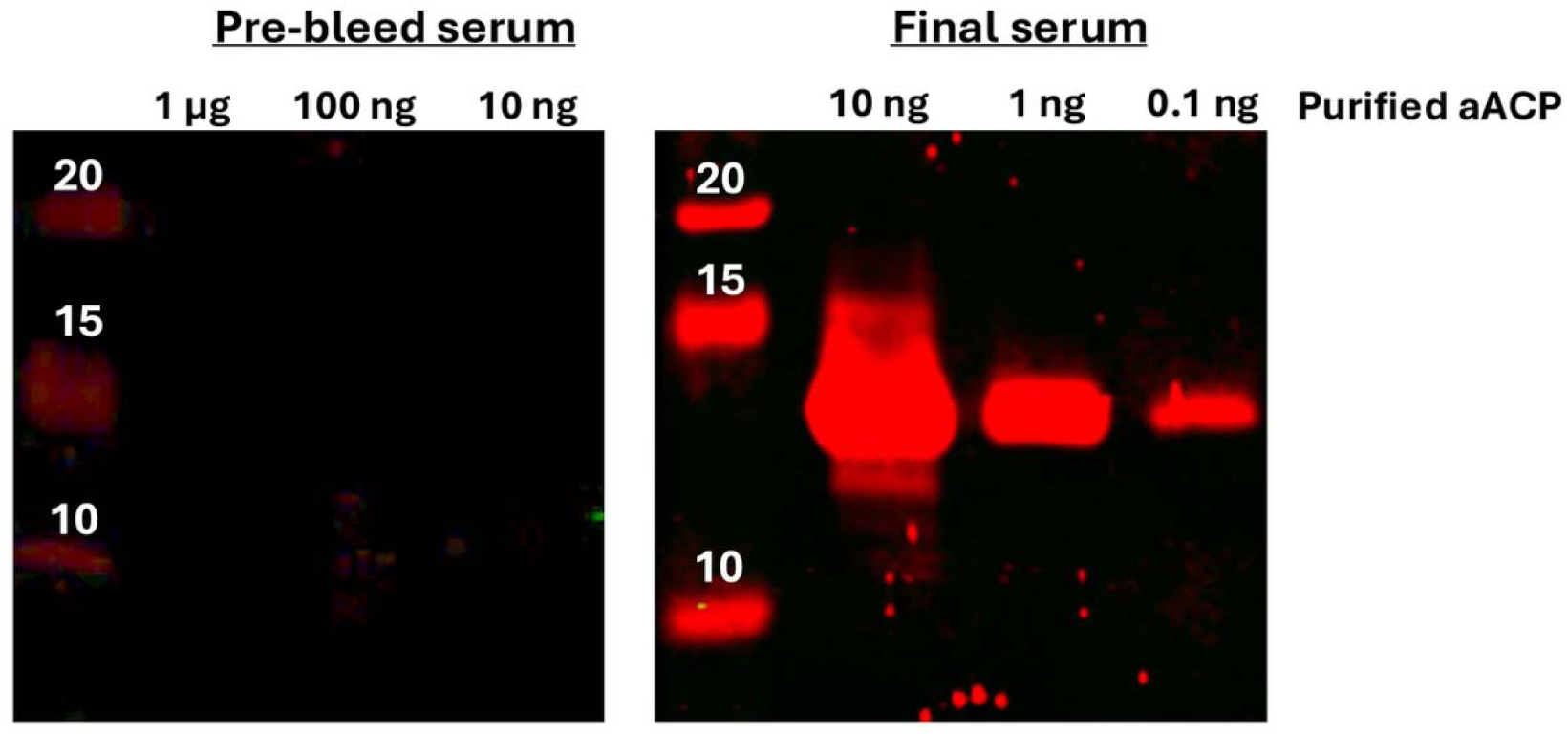
Validation of custom anti-apicoplast ACP antibody. Indicated amount of purified aACP antigen was loaded and fractionated on SDS-PAGE gel, transferred to membrane, blocked, and probed with 1:1000 dilution of rabbit pre-bleed serum or final serum, washed, and probed with donkey anti-rabbit IRDye680 secondary antibody.

**Figure 3- figure supplement 1.**
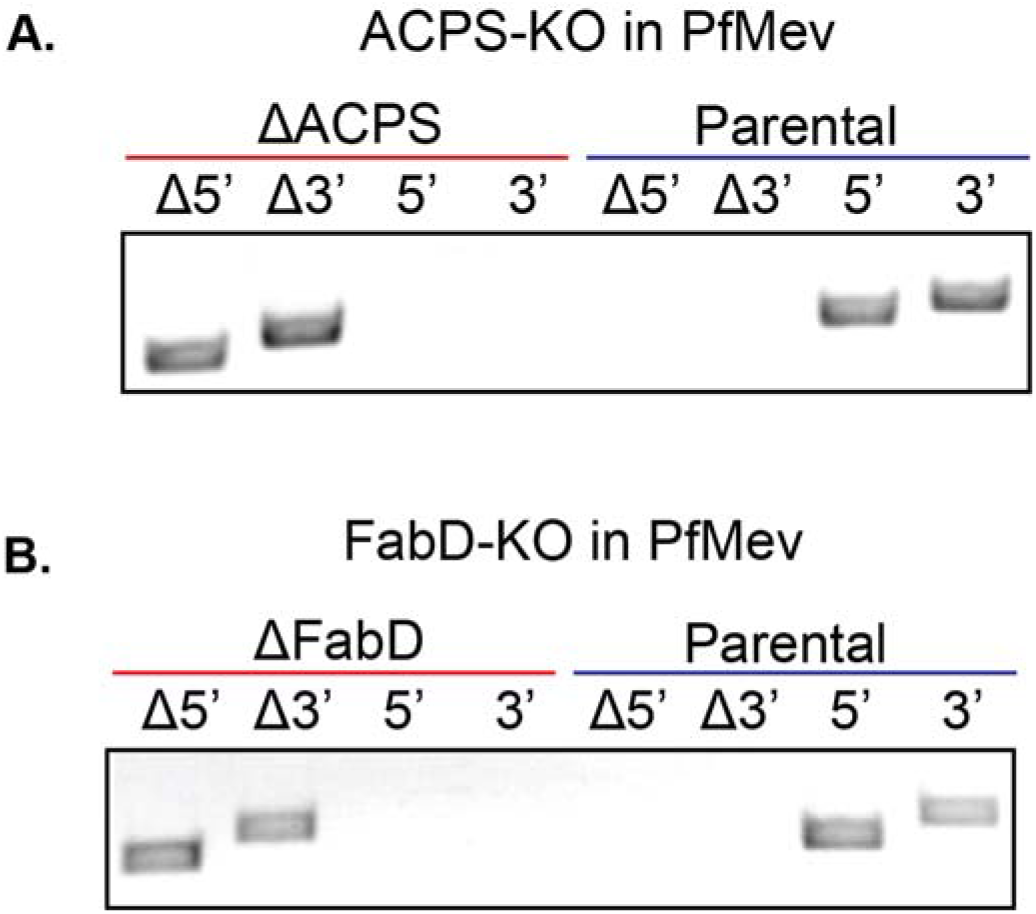
Integration PCRs for deletion of ACPS (**A**) or FabD (**B**) in PfMev parasites.

**Figure 3- figure supplement 2.**
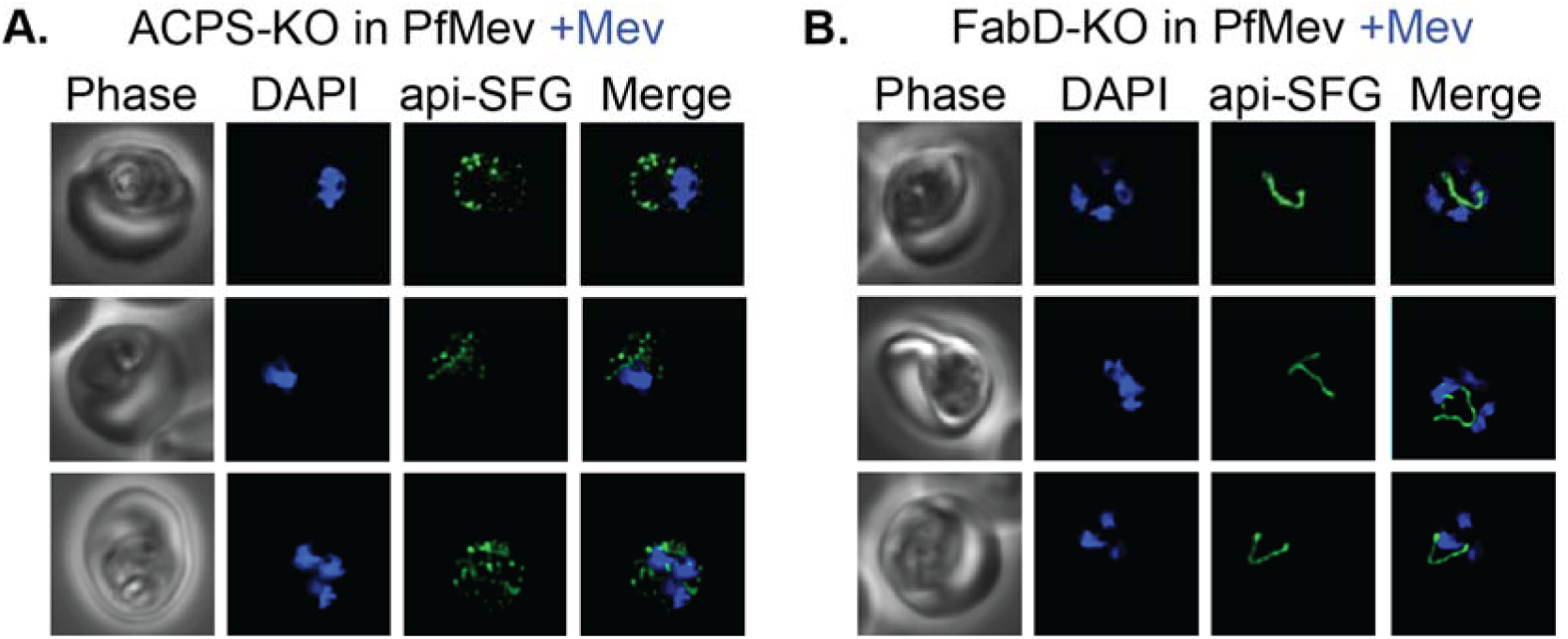
Additional microscopy images of apicoplast morphology in ΔACPS (**A**) and ΔFabD (**B**) PfMev parasites. Phase = phase contrast, api-SFG = apicoplast-targeted superfolder GFP.

**Figure 3- figure supplement 3.**
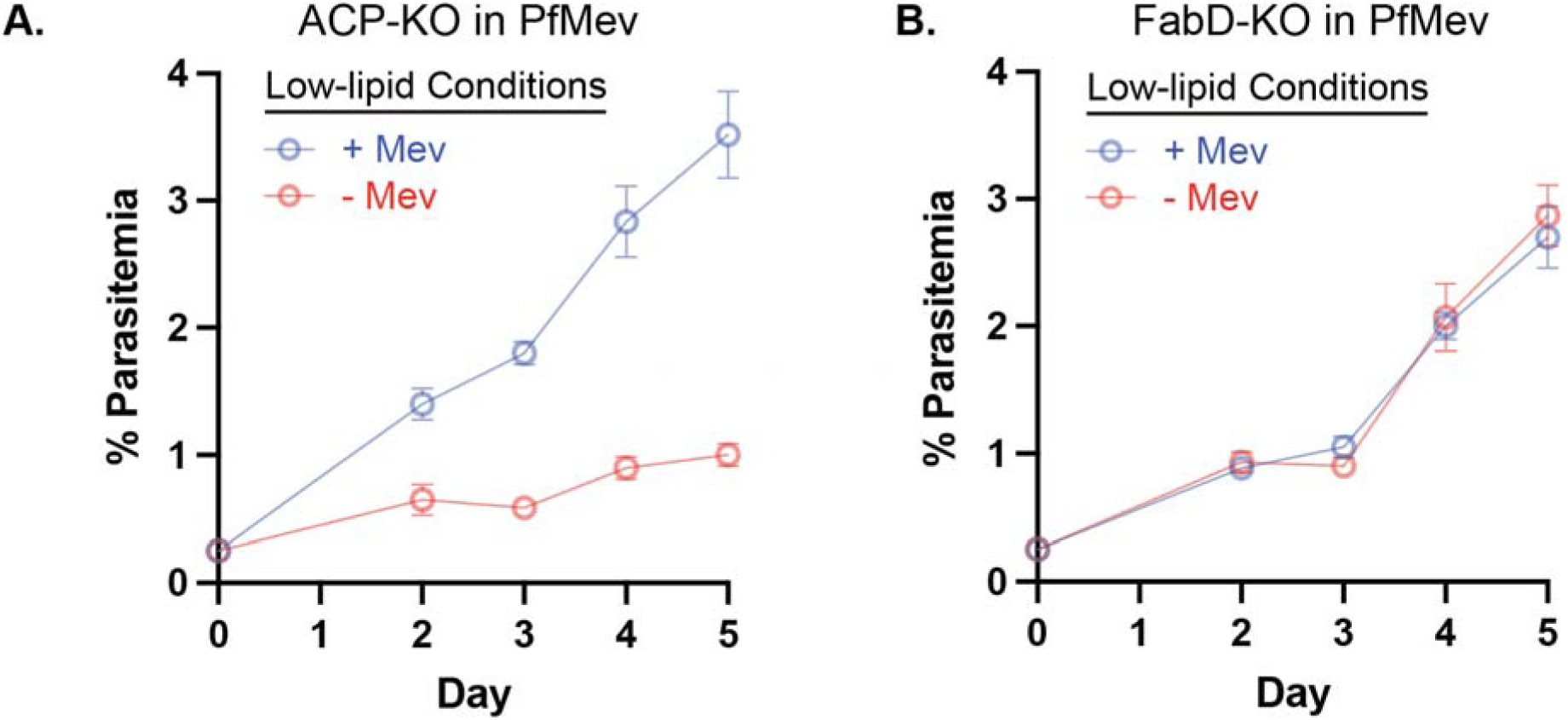
Growth of ΔACP (**A**) and ΔFabD (**B**) PfMev parasites in low-lipid conditions with daily media changes.

**Figure 4- figure supplement 1.**
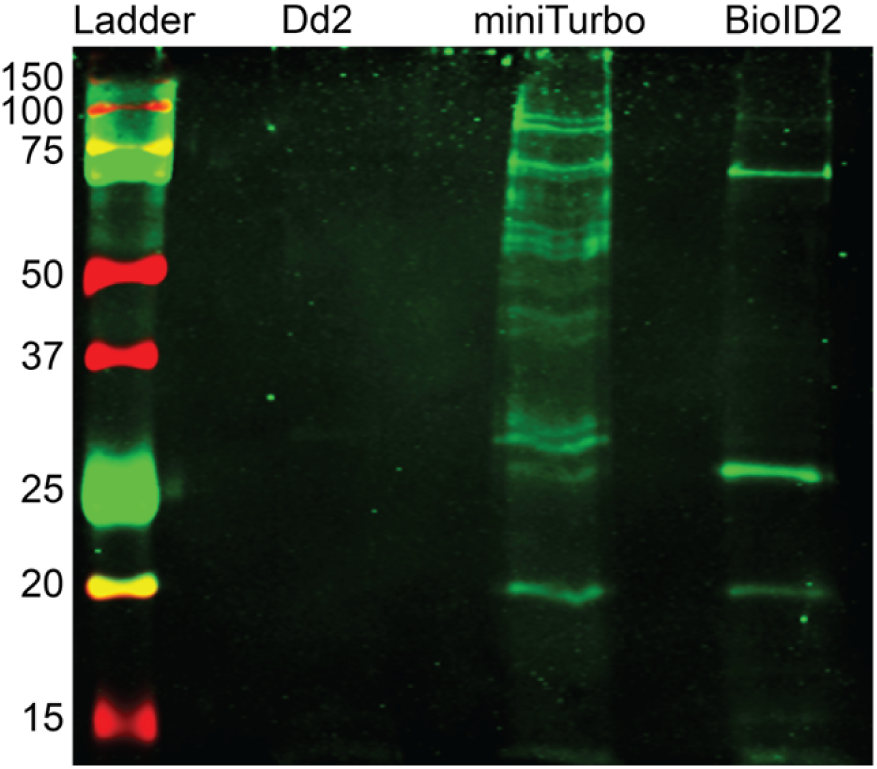
Western blot detection of biotinylated proteins in lysates from parental, aACP-miniTurbo, or aACP-BioID2 Dd2 parasites.

**Figure 4- figure supplement 2.**
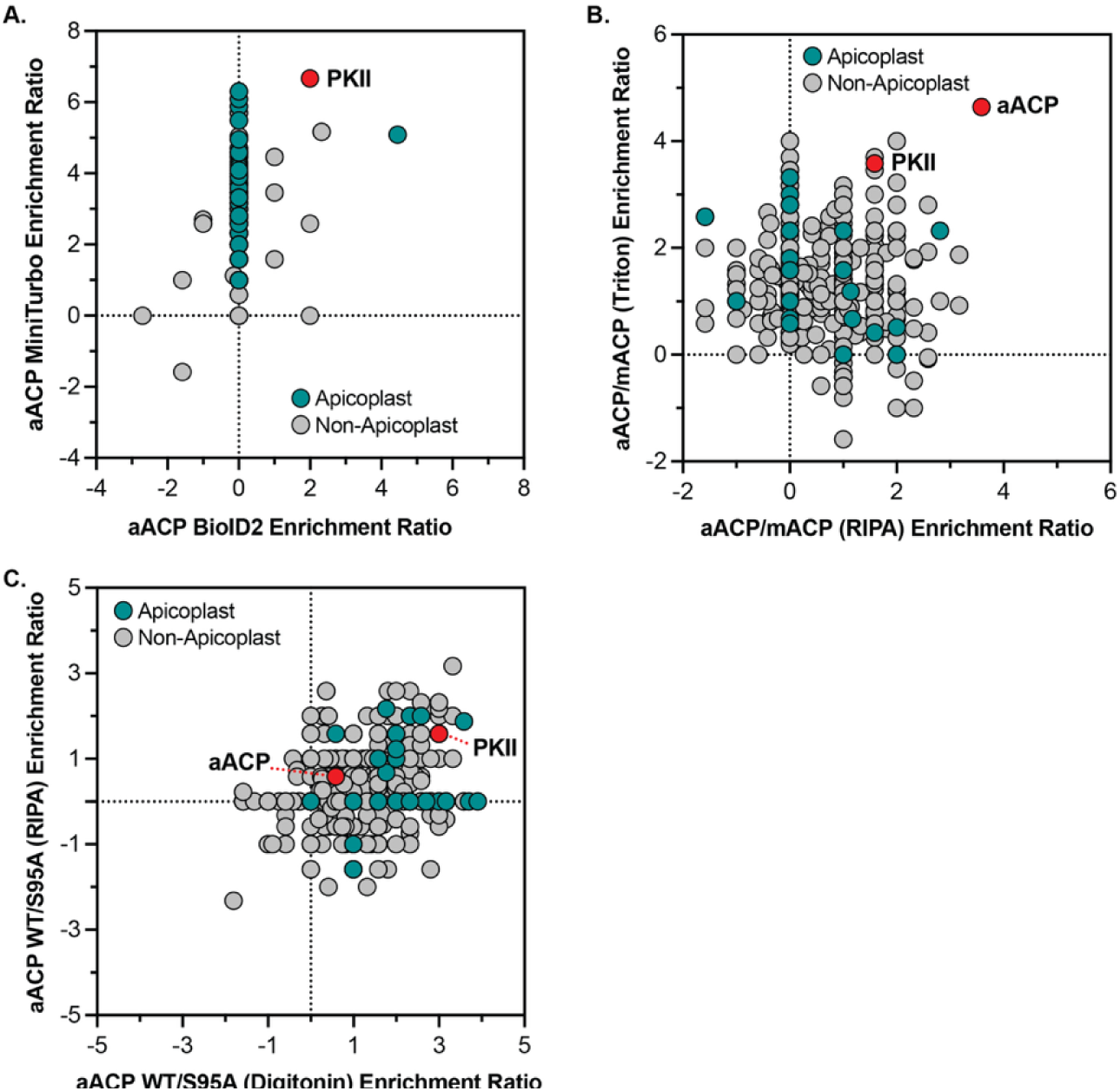
Enriched protein interactors of apicoplast ACP detected by proximity biotinylation or immunoprecipitation. (A) Apicoplast ACP interactors identified by miniTurbo versus BioID2. (B) Interacting proteins of ACP enriched in IP samples of apicoplast compared to mitochondrial ACP in Triton X-100 or RIPA buffer. (C) Interacting proteins of ACP enriched in IP samples of WT compared to S95A ACP in RIPA or digitonin buffer. All axes display the log_2_ enrichment ratio.

**Figure 4- figure supplement 3.**
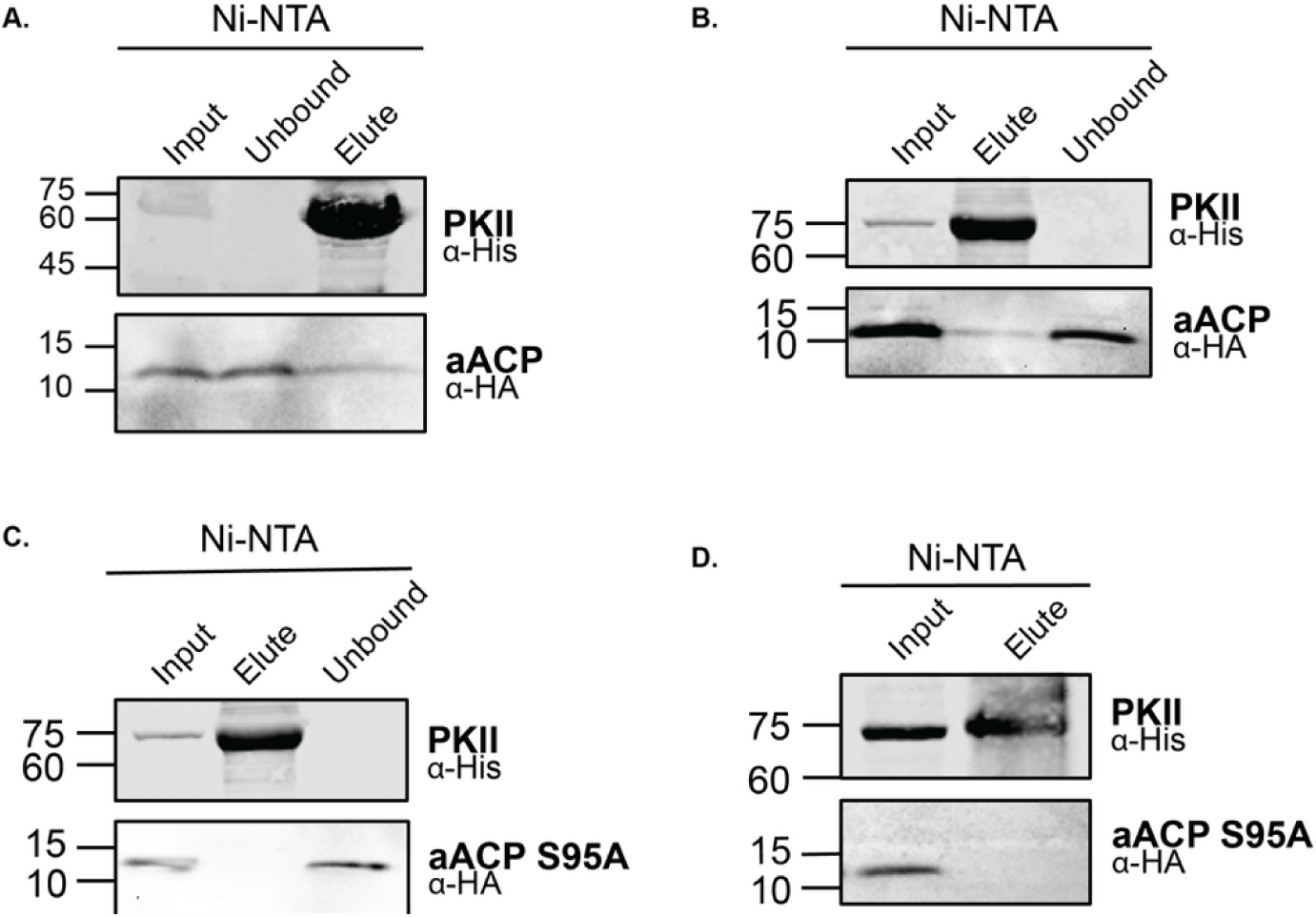
Replicate western blot images for Ni-NTA pulldown of PKII association with WT (A and B) or Ser95Ala (C and D) ACP in *E. coli*.

**Figure 5- figure supplement 1.**
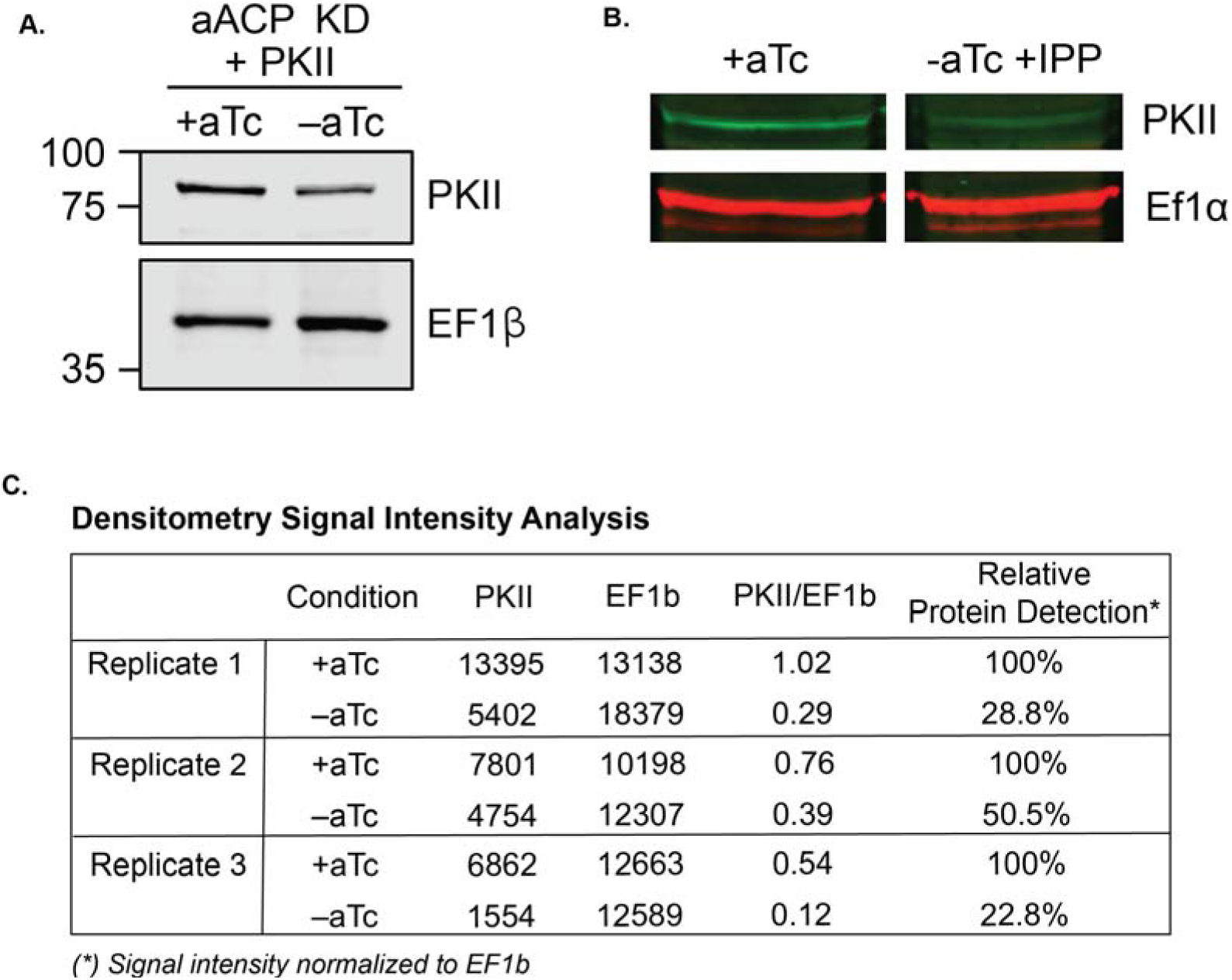
Replicate western images for ACP knockdown and PKII levels

**Figure 6- figure supplement 1.**
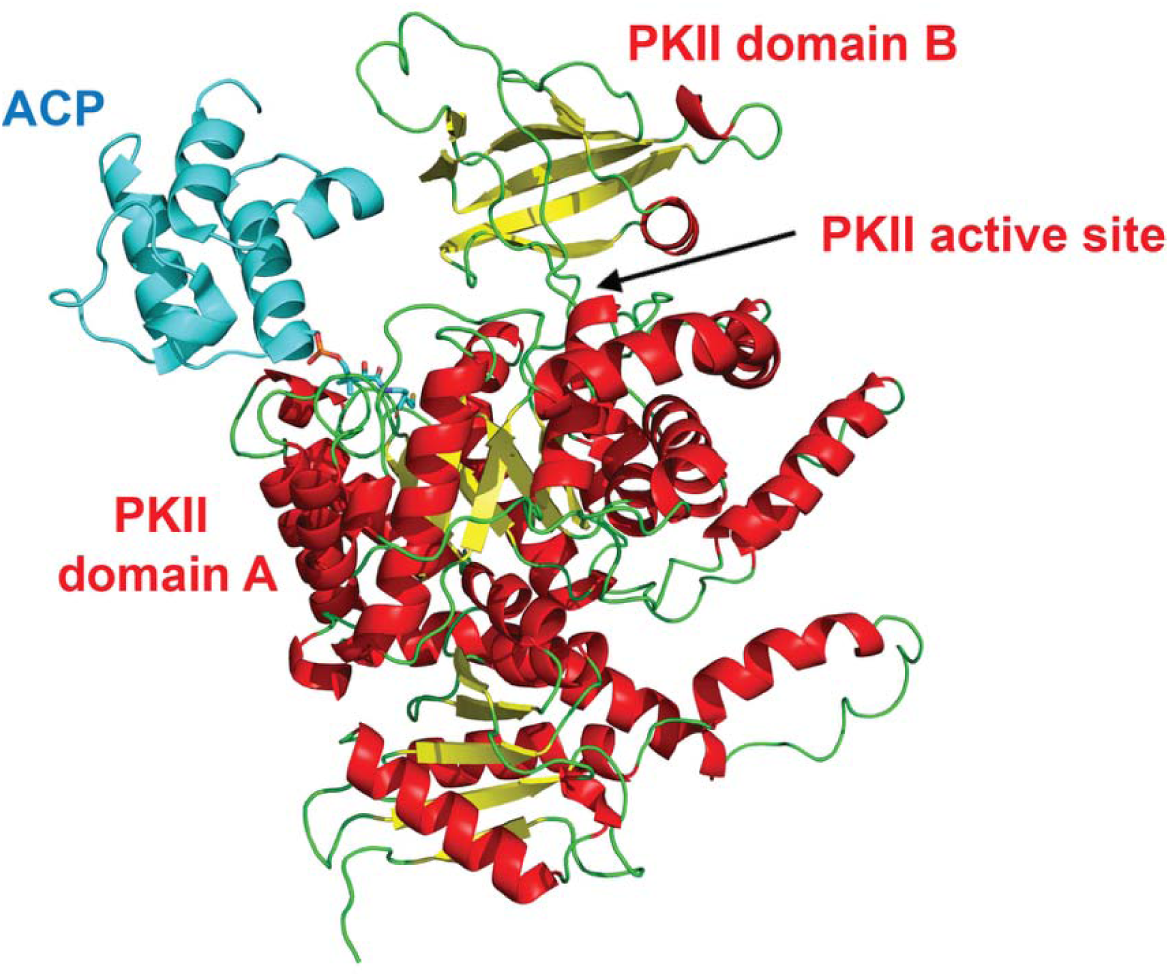
AlphaFold3 model of apicoplast ACP bound to pyruvate kinase II, rendered in PyMOL. The unstructured N-terminus of parasite PKII, which is expected to function as the apicoplast-targeting sequence and be cleaved upon organelle import, was not included in modeling. The presence of the 4-PP group was modeled by superposition with ACP in the PDB file 5USR.

