## Supplementary File 1 for "Acyl Carrier Protein is Essential for Apicoplast Biogenesis in Malaria Parasites Independent of Fatty Acid Synthesis"

**Supplementary Table 1**

Apicoplast-targeted proteins identified in parasites expressing ACP-miniTurbo or ACP-BioID2

| | | | | Enrichment Ratio | |
|---|---|---|---|---|---|
| **Plot ID** | **Gene ID** | **Annotation** | **Gene Name** | **MiniTurbo** | **BioID2** |
| PKII | PF3D7_1037100 | pyruvate kinase 2 | PKII | 6.672 | 2.000 |
| 1 | PF3D7_0816600 | chaperone protein ClpB1 | ClpB1 | 6.304 | 0.000 |
| 2 | PF3D7_0520800 | conserved Plasmodium protein, unknown function | N/A | 6.087 | 0.000 |
| 3 | PF3D7_1443900 | heat shock protein 90, putative | HSP90 | 5.883 | 0.000 |
| 4 | PF3D7_1025300 | conserved Plasmodium protein, unknown function | N/A | 5.492 | 0.000 |
| 5 | PF3D7_1232100 | 60 kDa chaperonin | CPN60 | 5.087 | 4.459 |
| 6 | PF3D7_1406600 | ATP-dependent Clp protease regulatory subunit ClpC, putative | ClpC | 4.954 | 0.000 |
| 7 | PF3D7_1440200 | stromal-processing peptidase, putative | SPP | 4.700 | 0.000 |
| 8 | PF3D7_0526700 | conserved protein, unknown function | N/A | 4.585 | 0.000 |
| 9 | PF3D7_1469600 | biotin carboxylase subunit of acetyl CoA carboxylase, putative | ACC | 4.585 | 0.000 |
| 10 | PF3D7_1103400 | FeS cluster assembly protein SufD, putative | SufD | 4.248 | 0.000 |
| 11 | PF3D7_0521400 | conserved Plasmodium protein, unknown function | N/A | 4.170 | 0.000 |
| 12 | PF3D7_1123500 | conserved Plasmodium protein, unknown function | N/A | 4.087 | 0.000 |
| 13 | PF3D7_1106500 | conserved Plasmodium protein, unknown function | N/A | 3.907 | 0.000 |
| 14 | PF3D7_0508800 | single-stranded DNA-binding protein | SSB | 3.585 | 0.000 |
| 15 | PF3D7_1411400 | plastid replication-repair enzyme | PREX | 3.322 | 0.000 |
| 16 | PF3D7_1358000 | patatin-like phospholipase, putative | PNPLA2 | 3.322 | 0.000 |
| 17 | PF3D7_0316400 | conserved Plasmodium protein, unknown function | N/A | 3.322 | 0.000 |
| 18 | PF3D7_1360800 | falcilysin | Falcilysin | 3.170 | 0.000 |
| 19 | PF3D7_1338600 | conserved Plasmodium protein, unknown function | N/A | 2.807 | 0.000 |
| 20 | PF3D7_1306200 | conserved Plasmodium protein, unknown function | N/A | 2.807 | 0.000 |
| 21 | PF3D7_1229600 | conserved Plasmodium protein, unknown function | N/A | 2.807 | 0.000 |
| 22 | PF3D7_1333000 | 20 kDa chaperonin | CPN20 | 2.807 | 0.000 |
| 23 | PF3D7_1406400 | pentatricopeptide repeat domain-containing protein, putative | PPR1 | 2.585 | 0.000 |
| 24 | PF3D7_1352000 | conserved Plasmodium protein, unknown function | N/A | 2.585 | 0.000 |
| 25 | PF3D7_0731600 | acyl-CoA synthetase | ACS5 | 2.585 | 0.000 |
| 26 | PF3D7_0916200 | conserved Plasmodium protein, unknown function | N/A | 2.322 | 0.000 |
| 27 | PF3D7_0607900 | conserved Plasmodium protein, unknown function | N/A | 2.322 | 0.000 |
| 28 | PF3D7_1413500 | FeS assembly ATPase SufC | SufC | 2.000 | 0.000 |
| 29 | PF3D7_1472700 | DNA-directed RNA polymerase, alpha subunit, putative | N/A | 2.000 | 0.000 |
| 30 | PF3D7_1223300 | DNA gyrase subunit A | GyrA | 2.000 | 0.000 |
| 31 | PF3D7_1427000 | conserved Plasmodium protein, unknown function | N/A | 2.000 | 0.000 |
| 32 | PF3D7_1323600 | conserved Plasmodium protein, unknown function | N/A | 2.000 | 0.000 |
| 33 | PF3D7_0313800 | conserved Plasmodium protein, unknown function | N/A | 2.000 | 0.000 |

| | | | | | |
|---|---|---|---|---|---|
| 34 | PF3D7_0504400 | ATP-dependent helicase, putative | N/A | 2.000 | 0.000 |
| 35 | PF3D7_0711000 | AAA family ATPase, CDC48 subfamily | CDC48 | 2.000 | 0.000 |
| 36 | PF3D7_0518100 | RAP protein, putative | AMR2 | 1.585 | 0.000 |
| 37 | PF3D7_0727500 | mTERF domain-containing protein, putative | N/A | 1.585 | 0.000 |
| 38 | PF3D7_1225100 | isoleucine--tRNA ligase, putative | api-IRS | 1.585 | 0.000 |
| 39 | PF3D7_0706100 | conserved Plasmodium protein, unknown function | N/A | 1.585 | 0.000 |
| 40 | PF3D7_0505700 | conserved Plasmodium membrane protein, unknown function | N/A | 1.585 | 0.000 |
| 41 | PF3D7_0904700 | bacterial histone-like protein | HU | 1.585 | 0.000 |
| 42 | PF3D7_1429100 | apicoplast ribosomal protein L15 precursor, putative | N/A | 1.585 | 0.000 |
| 43 | PF3D7_1337200 | 1-deoxy-D-xylulose 5-phosphate synthase | DXS | 1.585 | 0.000 |
| 44 | PF3D7_0804400 | methionine aminopeptidase 1c, putative | METAP1c | 1.000 | 0.000 |
| 45 | PF3D7_1307600 | DNA-directed RNA polymerase alpha chain, putative | rpoA | 1.000 | 0.000 |
| 46 | PF3D7_0920200 | CS domain protein, putative | N/A | 1.000 | 0.000 |
| 47 | PF3D7_1470800 | conserved Plasmodium protein, unknown function | N/A | 1.000 | 0.000 |
| 48 | PF3D7_1369600 | conserved Plasmodium protein, unknown function | N/A | 1.000 | 0.000 |
| 49 | PF3D7_0827600 | conserved Plasmodium protein, unknown function | N/A | 1.000 | 0.000 |
| 50 | PF3D7_0505400 | conserved Plasmodium protein, unknown function | N/A | 1.000 | 0.000 |
| 51 | PF3D7_0509600 | asparagine--tRNA ligase | AsnRS | 1.000 | 0.000 |
| 52 | PF3D7_0827500 | apicoplast ribosomal protein L21 precursor, putative | N/A | 1.000 | 0.000 |
| 53 | PF3D7_1022800 | 4-hydroxy-3-methylbut-2-en-1-yl diphosphate synthase | ISPG | 1.000 | 0.000 |

**Supplementary Table 2**

Apicoplast-targeted proteins identified by IP/MS of apicoplast versus mitochondrial ACP-$HA_2$

| | | | | aACP/mACP Enrichment Ratio | |
|---|---|---|---|---|---|
| **Plot ID** | **Gene ID** | **Annotation** | **Gene Name** | **RIPA** | **Urea** |
| aACP | PF3D7_0208500 | acyl carrier protein | aACP | 3.585 | 4.644 |
| PKII | PF3D7_1037100 | pyruvate kinase 2 | PKII | 1.585 | 3.585 |
| 1 | PF3D7_1126000 | threonine--tRNA ligase | ThrRS | 2.807 | 2.322 |
| 2 | PF3D7_0731600 | acyl-CoA synthetase | ACS5 | 2.000 | 0.515 |
| 3 | PF3D7_0912400 | alkaline phosphatase, putative | Alkaline Phosphatase | 2.000 | 0.000 |
| 4 | PF3D7_1123500 | conserved Plasmodium protein, unknown function | N/A | 2.000 | 0.000 |
| 5 | PF3D7_1410700 | conserved Plasmodium protein, unknown function | N/A | 1.585 | 0.415 |
| 6 | PF3D7_1232100 | 60 kDa chaperonin | CPN60 | 1.170 | 0.670 |
| 7 | PF3D7_1360800 | falcilysin | FLN | 1.138 | 1.184 |
| 8 | PF3D7_1420400 | glycine--tRNA ligase | GlyRS | 1.000 | 2.322 |
| 9 | PF3D7_0816600 | chaperone protein ClpB1 | ClpB1 | 1.000 | 1.585 |
| 10 | PF3D7_0627700 | transportin | Transportin | 1.000 | 1.585 |
| 11 | PF3D7_1212000 | glutathione peroxidase-like thioredoxin peroxidase | TPx(Gl) | 1.000 | 0.000 |
| 12 | PF3D7_1367700 | alanine--tRNA ligase | AlaRS | 0.000 | 3.322 |
| 13 | PF3D7_1025300 | conserved Plasmodium protein, unknown function | N/A | 0.000 | 3.322 |
| 14 | PF3D7_0111500 | UMP-CMP kinase, putative | UMP-CMP Kinase | 0.000 | 3.000 |
| 15 | PF3D7_1437200 | ribonucleoside-diphosphate reductase large subunit, putative | RNR-LSU | 0.000 | 2.807 |
| 16 | PF3D7_0529000 | conserved Plasmodium protein, unknown function | N/A | 0.000 | 2.807 |
| 17 | PF3D7_0623500 | superoxide dismutase [Fe] | SOD2 | 0.000 | 2.322 |
| 18 | PF3D7_1457300 | conserved Plasmodium protein, unknown function | N/A | 0.000 | 1.807 |
| 19 | PF3D7_1443900 | heat shock protein 90, putative | HSP90 | 0.000 | 1.585 |
| 20 | PF3D7_1337200 | 1-deoxy-D-xylulose 5-phosphate synthase | DXS | 0.000 | 1.585 |
| 21 | PF3D7_1205700 | targeted glyoxalase II | tGLO2 | 0.000 | 1.585 |
| 22 | PF3D7_0520800 | conserved Plasmodium protein, unknown function | N/A | 0.000 | 1.585 |
| 23 | PF3D7_1119000 | acyl-CoA-binding protein, putative | Acyl-CoA-binding protein | 0.000 | 1.000 |
| 24 | PF3D7_0505700 | conserved Plasmodium membrane protein, unknown function | N/A | 0.000 | 1.000 |
| 25 | PF3D7_0721100 | conserved Plasmodium protein, unknown function | N/A | 0.000 | 0.678 |
| 26 | PF3D7_0209300 | 2C-methyl-D-erythritol 2,4-cyclodiphosphate synthase | IspF | 0.000 | 0.585 |

**Supplementary Table 3**

Apicoplast-targeted proteins identified by IP/MS of WT or Ser95Ala ACP-$HA_2$

| Plot ID | Gene ID | Annotation | Gene Name | aACP/S95A Enrichment Ratio | |
|---|---|---|---|---|---|
| | | | | Digitonin | RIPA |
| aACP | PF3D7_0208500 | acyl carrier protein | aACP | 0.585 | 0.585 |
| PKII | PF3D7_1037100 | pyruvate kinase 2 | PKII | 3.000 | 1.585 |
| 1 | PF3D7_1307600 | DNA-directed RNA polymerase alpha chain, putative | rpoA | 3.907 | 0.000 |
| 2 | PF3D7_1338600 | conserved Plasmodium protein, unknown function | N/A | 3.700 | 0.000 |
| 3 | PF3D7_1360800 | falcilysin | FLN | 3.585 | 1.874 |
| 4 | PF3D7_1364600 | aldehyde reductase, putative | Aldehyde reductase | 3.170 | 0.000 |
| 5 | PF3D7_0820800 | conserved Plasmodium protein, unknown function | N/A | 3.170 | 0.000 |
| 6 | PF3D7_0920600 | conserved Plasmodium protein, unknown function | N/A | 3.170 | 0.000 |
| 7 | PF3D7_1025300 | conserved Plasmodium protein, unknown function | N/A | 3.170 | 0.000 |
| 8 | PF3D7_1005900 | conserved Plasmodium protein, unknown function | N/A | 3.000 | 0.000 |
| 9 | PF3D7_1228700 | conserved Plasmodium protein, unknown function | N/A | 2.700 | 0.000 |
| 10 | PF3D7_1212000 | glutathione peroxidase-like thioredoxin peroxidase | TPx(Gl) | 2.585 | 2.000 |
| 11 | PF3D7_0912400 | alkaline phosphatase, putative | Alkaline Phosphatase | 2.322 | 2.000 |
| 12 | PF3D7_0313700 | conserved Plasmodium protein, unknown function, unspecified product | N/A | 2.322 | 0.000 |
| 13 | PF3D7_1021000 | conserved Plasmodium protein, unknown function | N/A | 2.322 | 0.000 |
| 14 | PF3D7_1410700 | conserved Plasmodium protein, unknown function | N/A | 2.000 | 1.585 |
| 15 | PF3D7_1126000 | threonine--tRNA ligase | ThrRS | 2.000 | 1.222 |
| 16 | PF3D7_1443900 | heat shock protein 90, putative | HSP90 | 2.000 | 1.000 |
| 17 | PF3D7_0731600 | acyl-CoA synthetase | ACS5 | 2.000 | 0.000 |
| 18 | PF3D7_1221800 | conserved Plasmodium protein, unknown function | N/A | 2.000 | 0.000 |
| 19 | PF3D7_1306200 | conserved Plasmodium protein, unknown function | N/A | 2.000 | 0.000 |
| 20 | PF3D7_1323600 | conserved Plasmodium protein, unknown function | N/A | 2.000 | 0.000 |
| 21 | PF3D7_1457300 | conserved Plasmodium protein, unknown function | N/A | 2.000 | 0.000 |
| 22 | PF3D7_1232100 | 60 kDa chaperonin | CPN60 | 1.766 | 2.170 |
| 23 | PF3D7_0816600 | chaperone protein ClpB1 | ClpB1 | 1.766 | 0.678 |
| 24 | PF3D7_1420400 | glycine--tRNA ligase | GlyRS | 1.585 | 1.000 |
| 25 | PF3D7_0627700 | transportin | Transportin | 1.585 | 1.000 |
| 26 | PF3D7_1123500 | conserved Plasmodium protein, unknown function | N/A | 1.585 | 1.000 |
| 27 | PF3D7_1472700 | DNA-directed RNA polymerase, alpha subunit, putative | N/A | 1.585 | 0.000 |
| 28 | PF3D7_1440200 | stromal-processing peptidase, putative | SPP | 1.585 | 0.000 |
| 29 | PF3D7_1419200 | thioredoxin-like protein, putative | ATrx1 | 1.585 | 0.000 |
| 30 | PF3D7_1367700 | alanine--tRNA ligase | AlaRS | 1.585 | 0.000 |
| 31 | PF3D7_0405400 | pre-mRNA-processing-splicing factor 8, putative | PRPF8 | 1.585 | 0.000 |
| 32 | PF3D7_0721100 | conserved Plasmodium protein, unknown function | N/A | 1.585 | 0.000 |

| | | | | | |
|---|---|---|---|---|---|
| 33 | PF3D7_1226900 | conserved Plasmodium protein, unknown function | N/A | 1.585 | 0.000 |
| 34 | PF3D7_1409100 | aldo-keto reductase, putative | Aldo-Keto Reductase | 1.000 | 0.000 |
| 35 | PF3D7_1205700 | targeted glyoxalase II | tGLO2 | 1.000 | 0.000 |
| 36 | PF3D7_1015200 | cysteine--tRNA ligase | CysRS | 1.000 | 0.000 |
| 37 | PF3D7_0711000 | AAA family ATPase, CDC48 subfamily | CDC48 | 1.000 | 0.000 |
| 38 | PF3D7_0530200 | phosphoenolpyruvate/phosphate translocator | PPT | 1.000 | 0.000 |
| 39 | PF3D7_0504400 | ATP-dependent helicase, putative | ATP-dependent Helicase | 1.000 | 0.000 |
| 40 | PF3D7_0111500 | UMP-CMP kinase, putative | UMP-CMP kinase | 1.000 | 0.000 |
| 41 | PF3D7_0624400 | conserved Plasmodium protein, unknown function | N/A | 1.000 | 0.000 |
| 42 | PF3D7_1125000 | conserved Plasmodium protein, unknown function | N/A | 1.000 | 0.000 |
| 43 | PF3D7_0209300 | 2C-methyl-D-erythritol 2,4-cyclodiphosphate synthase | IspF | 0.585 | 1.585 |

**Supplementary Table 4**

Apicoplast-targeted protein interactors of aACP identified by both miniTurbo and IP/MS

(common to Supplementary Tables 1 and 2)

| No. | Gene ID | Annotation | Gene Name |
|---|---|---|---|
| 1 | PF3D7_1232100 | 60 kDa chaperonin | CPN60 |
| 2 | PF3D7_1025300 | conserved protein, unknown function | N/A |
| 3 | PF3D7_1443900 | heat shock protein 90, putative | HSP90 |
| 4 | PF3D7_0505700 | conserved Plasmodium membrane protein, unknown function | N/A |
| 5 | PF3D7_0816600 | chaperone protein ClpB1 | ClpB1 |
| 6 | PF3D7_0731600 | acyl-CoA synthetase | ACS5 |
| 7 | PF3D7_1337200 | 1-deoxy-D-xylulose 5-phosphate synthase | DXS |
| 8 | PF3D7_0520800 | conserved protein, unknown function | N/A |
| 9 | PF3D7_1360800 | falcilysin | FLN |
| 10 | PF3D7_1037100 | pyruvate kinase 2 | PKII |
| 11 | PF3D7_1123500 | golgi protein 2 | N/A |

**Supplementary Table 5**

Primers sequences used for cloning and PCR analyses

| Prime | Name | Sequence |
|---|---|---|
| P1 | aACP_S95A_F | GGGGGCTGATGCTTTAGACCTCGTTG |
| P2 | aACP_S95A_R | CAACGAGGTCTAAAGCATCAGCCCCC |
| P3 | aACP_TEOE_F | aaacacgattttttctcgagATGAAGATCTTATTACTTTG |
| P4 | aACP_BioID2_R | AAtcctccacttcccctaggTTGCTTATTATTTTTTTCTATATAA |
| P5 | aACP_miniTurbo_ | ATtcctccacttcccctaggTTGCTTATTATTTTTTTCTATATAA |
| P6 | BioID2_aACP_F | ACTAAACATTTCGAAAAAGTAAAATCAcctaggggaagtggaggaTTCAAGAACCTGATC |
| P7 | BioID2_HA5_R | GATAAGCGTAATCTGGAACATCGTATGGGTAGGCATAATCTGGAACATCGTAAGGATACG |
| P8 | miniTurbo_aACP_ | GATATTAACTAAACATTTCGAAAAAGTAAAATCAcctaggggaagtggaggaatcccgct |
| P9 | miniTurbo_aACP_ | GATAAGCGTAATCTGGAACATCGTATGGGTAGGCATAATCTGGAACATCGTAAGGATACG |
| P10 | PKII_TEOE_F | aaacacgattttttctcgagATGAATTTAATTCATATTTGTTTATTTATC |
| P11 | PKII_Myc3_R | TCTGAGATGAGTTTTTGTTCCCTAGGATTTGTTAGACATGGTTGACATAA |
| P12 | Myc3_PKII_F | TTATGTCAACCATGTCTAACAAATCCTAGGGAACAAAAACTCATCTCAGAAGAGGATCTG |
| P13 | Myc3_PKII_R | gtacataaatatattatataactcgacgCGGCCGC |
| P14 | aACP_KD_gRNA_ | TAAGTATATAATATTAGACCTCGTTGAATTAATTAGTTTTAGAGCTAGAA |
| P15 | aACP_KD_gRNA_ | TTCTAGCTCTAAAACTAATTAATTCAACGAGGTCTAATATTATATACTTA |
| P16 | aACP_5'UTR_F | CCATAACGATAGGGGAGAAATATTATATATAGC |
| P17 | aACP_3'UTR_R | ATAAAAATACCGTTAACTTAAGATATATGTTAACCCAAATTAGCTAGCCAATTCATTT |
| P18 | pMG75_apt_R | CCCAGGCCTCTAGTTTAC |
| P19 | aACPKO_5'HF_F | gccacgaGCGGCCtgaagtatatatccacgtccataacag |
| P20 | aACPKO_5'HF_R | aagcgcaGCGGCCccatctttttgtgtatttttaaaagct |
| P21 | aACPKO_3'HF_F | cgacagacGCCGGggtacaaaaaaataaaatgatacacaa |
| P22 | aACPKO_3'HF_R | ggccaccaGCCGGgggatacattttaacaaagaagaaata |
| P23 | ACPSKO_5'HF_F | gccacgaGCGGCCccaggagttcacacacataatagattg |
| P24 | ACPSKO_5'HF_R | aagcgcaGCGGCCattcggcatttgtttttgtattcaatg |
| P25 | ACPSKO_3'HF_F | cgacagacGCCGGgaggggtcaataaattgttttccaata |
| P26 | ACPSKO_3'HF_R | ggccaccaGCCGGaccctctaccaatagatttgacgatag |
| P27 | FabDKO_5'HF_F | gccacgaGCGGCCacactcacactcacactcacattatcc |
| P28 | FabDKO_5'HF_R | aagcgcaGCGGCCacgaccctatcttatttgtattctcct |
| P29 | FabDKO_3'HF_F | cgacagacGCCGGgtggaaaaccagaatctatggattatc |
| P30 | FabDKO_3'HF_R | ggccaccaGCCGGggtggtatcatatatatggaattgtcc |
| P31 | aACPKO_gRNA_F | TAAGTATATAATATTaatgaactcaaattttaccaGTTTTAGAGCTAGAA |
| P32 | aACPKO_gRNA_R | TTCTAGCTCTAAAACtggtaaaatttgagttcattAATATTATATACTTA |
| P33 | ACPSKO_gRNA_F | TAAGTATATAATATTtaaaatattgaacccacgagGTTTTAGAGCTAGAA |
| P34 | ACPSKO_gRNA_R | TTCTAGCTCTAAAACctcgtgggttcaatattttaAATATTATATACTTA |
| P35 | FabDKO_gRNA_F | TAAGTATATAATATTgatagaaaatttggtttatgGTTTTAGAGCTAGAA |
| P36 | FabDKO_gRNA_R | TTCTAGCTCTAAAACcataaaccaaattttctatcAATATTATATACTTA |
| P37 | ACP.5F | cctatgtttgtgtactttttttttcccc |
| P38 | ACP.5WTR | ggttgcttcttttctttttaatatttgtaag |
| P39 | ACP.3WTF | cccatctagctctttaaaaagtacttttg |
| P40 | ACP.3R | catacataaatatatgaacatttttataaagaggtaac |
| P41 | ACPS.5F | gtaatatttgtggatatatatttctttacataccttgc |
| P42 | ACPS.5WTR | gtaatatcatttctcttacaaaaagaagataaaatattc |
| P43 | ACPS.3WTF | cagaacaagttgtaatgtttatgatattagtgttg |

| | | |
|---|---|---|
| P44 | ACPS.3R | cctaattgtctagctattttttttgctttg |
| P45 | FabD.5F | gagaagatatattaaaagagaaatttttttttttttttcg |
| P46 | FabD.5WTR | ttccttttatatctttatttacactaacattatgtctaag |
| P47 | FabD.3WTF | acaaaatgaatgatgatatttttattgttagttatatgac |
| P48 | FabD.3R | ctttaaaatataaaatatacttttttttgtcaatttgtaag |
| P49 | pRS.R | TACAAAATGCTTAAGCGCAGCGGCC |
| P50 | pRS.F | CATATTTATTAAATCTAGAATTCGACAGACGCCG |
| P51 | LDH.F | GGAGATGTAGTTTTGTTCGATATTG |
| P52 | LDH.R | CTTGTAAAGGGATACCACCTACAG |
| P53 | SufB.F | CATGTAGCTATAGTAGAAATAATAGTAAAAGATTATGG |
| P54 | SufB.R | GACTCTGAAATACTTAAACCACGTTGC |
| P55 | Cox1.F | CTTCATCTTTAAGAATAATTGCACAAGAAAATGTAAATC |
| P56 | Cox1.R | GGAAGCTTAGTATGGGTACATCATATGTAC |
