## Supplementary File 2 for "Acyl Carrier Protein is Essential for Apicoplast Biogenesis in Malaria Parasites Independent of Fatty Acid Synthesis"

A

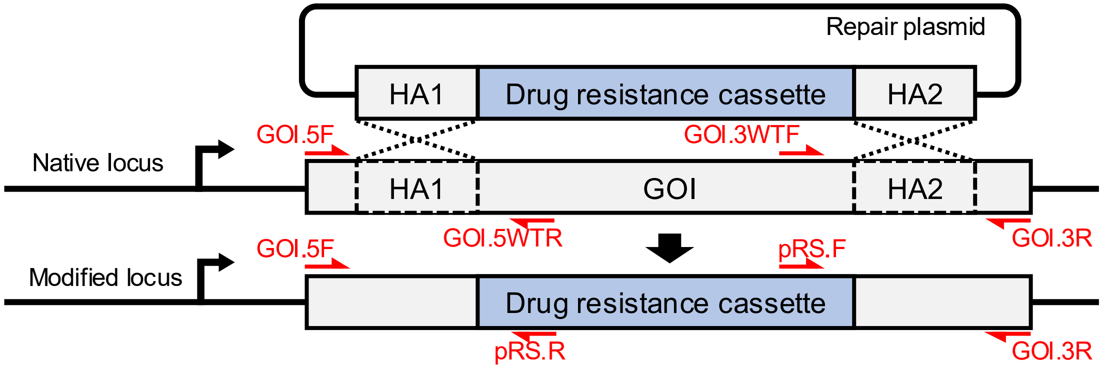


B

| PCR reaction | Primer combination | | Parasite line |
|---|---|---|---|
| | Forward | Reverse | |
| Δ5’ | GOI.5F | pRS.R | Gene knockout lines |
| Δ3’ | pRS.F | GOI.3R | |
| 5’ | GOI.5F | GOI.5WTR | |
| 3’ | GOI.3WTF | GOI.3R | |
| Δ5’ | GOI.5F | pRS.R | PfMev (parental) |
| Δ3’ | pRS.F | GOI.3R | |
| 5’ | GOI.5F | GOI.5WTR | |
| 3’ | GOI.3WTF | GOI.3R | |

C

| | Anticipated amplicon sizes (bp) | | | |
|---|---|---|---|---|
| Gene of interest | Δ5’ | Δ3’ | 5’ | 3’ |
| *acp* | 526 | 526 | 539 | 596 |
| *acps* | 547 | 584 | 560 | 654 |
| *fabD* | 488 | 547 | 501 | 617 |

Genomic PCR analysis to confirm targeted gene disruption. (A) Schematic depiction of targeted gene and primer-annealing sites. (B) Primer combinations to test 5’ or 3’ integration or confirm parental locus. (C) Expected PCR amplicon sizes for indicated primer pairs.
