## Supplementary material for "Acyl Carrier Protein is Essential for Apicoplast Biogenesis in Malaria Parasites Independent of Fatty Acid Synthesis": Source Data

Figure 2 – source data 1

A. Uncropped PCR gels for deletion of ACP in Mev Bypass NF54

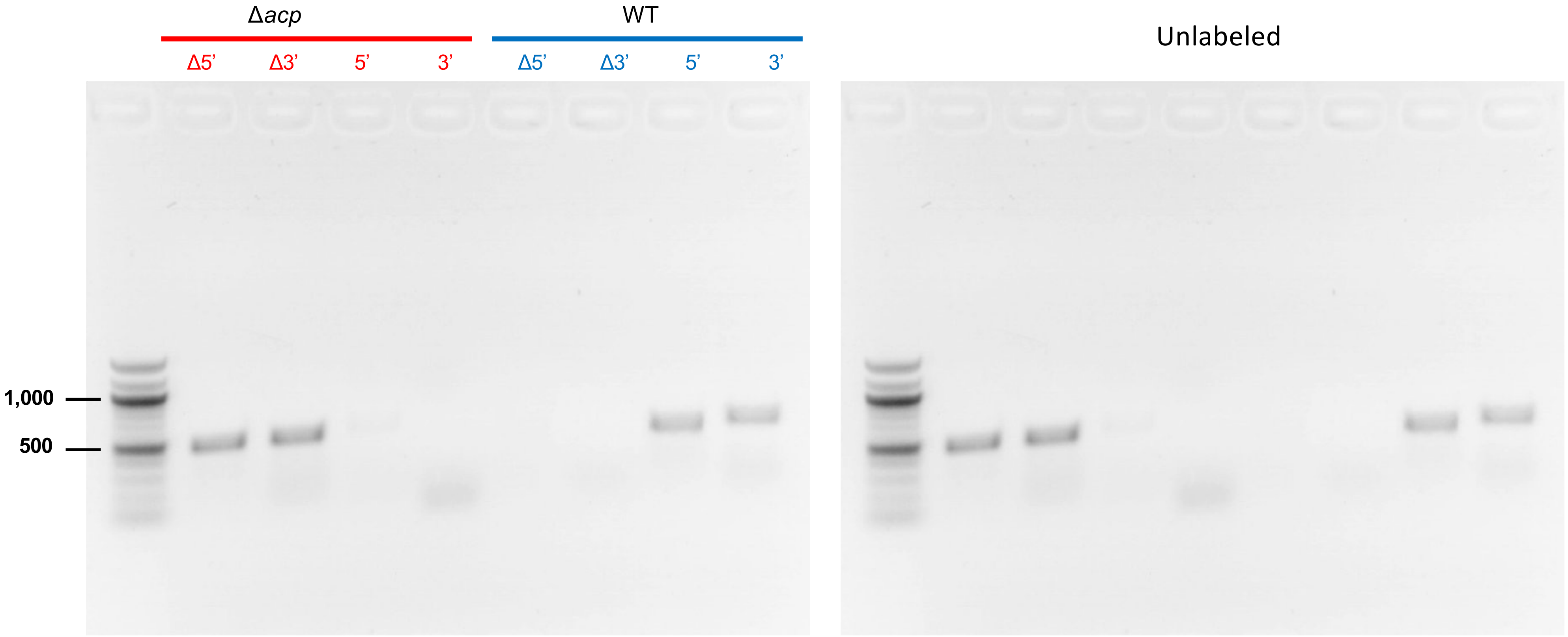

Figure 2 – source data 1

B. Uncropped PCR gels for organellar genomes in ACP-KO and Wt Mev Bypass NF54

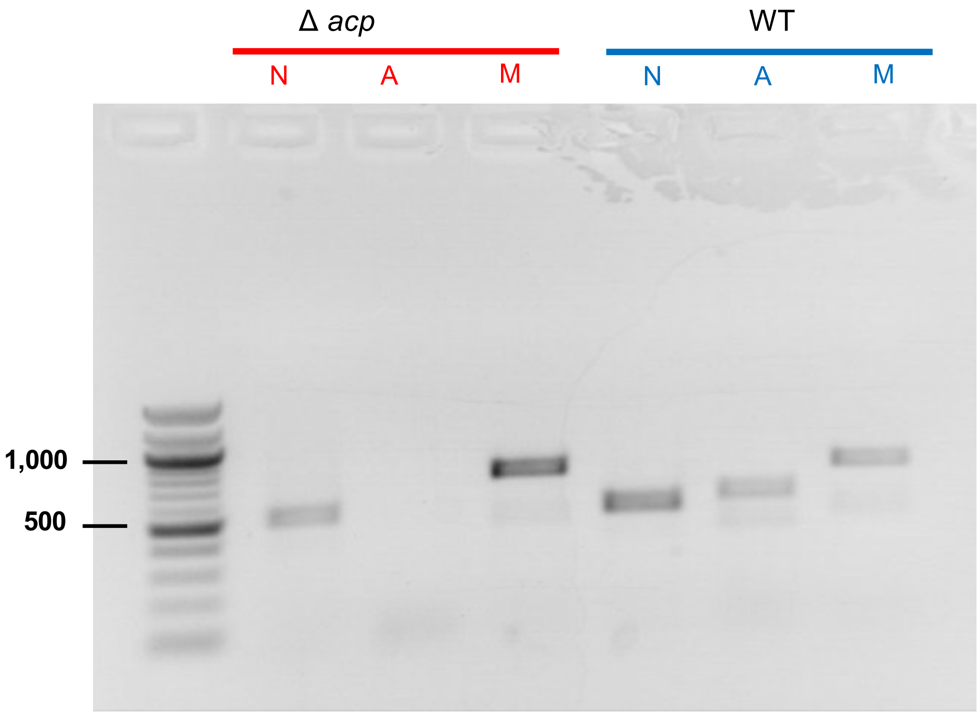


Unlabeled

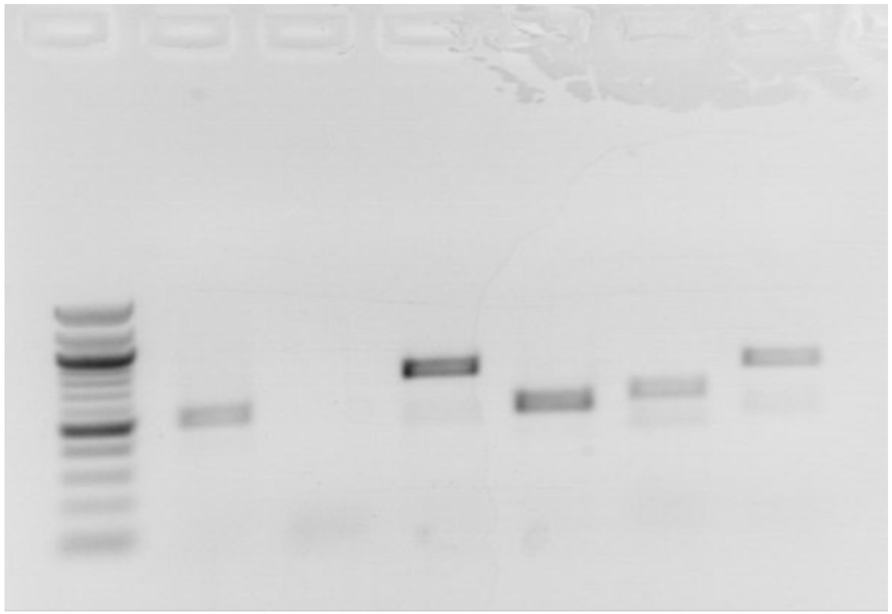

N = nuclear gene
A = apicoplast gene
M = mitochondrial gene

Figure 2 – source data 1

C. Uncropped PCR gels for integration of ACP-KD system into Dd2

Labeled

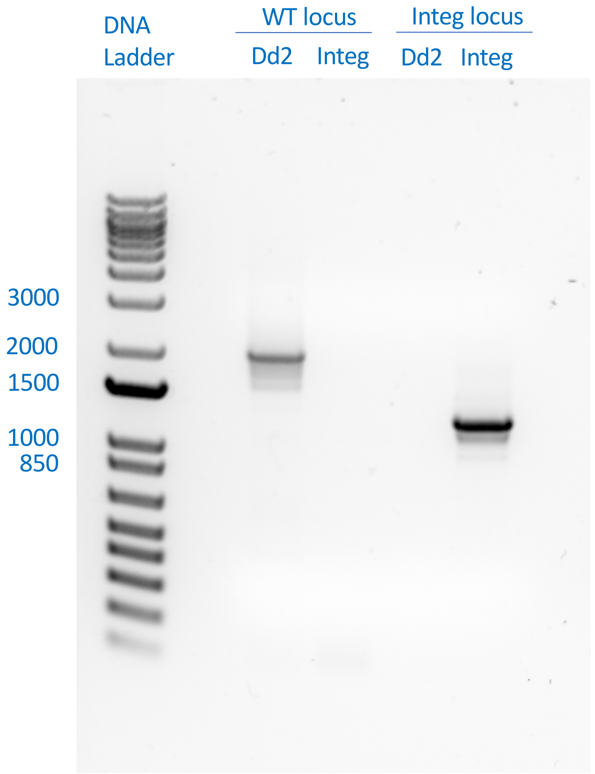


Unlabeled

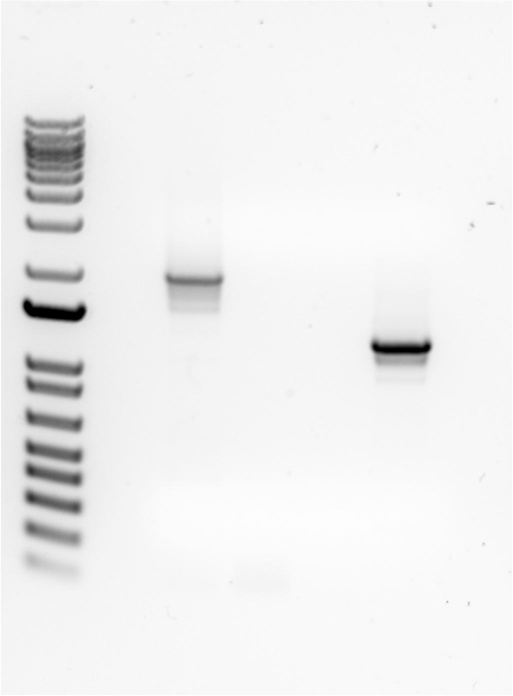

Figure 2 – source data 1

D. Uncropped PCR gels for integration of ACP-KD system into Mev Bypass NF54

Labeled

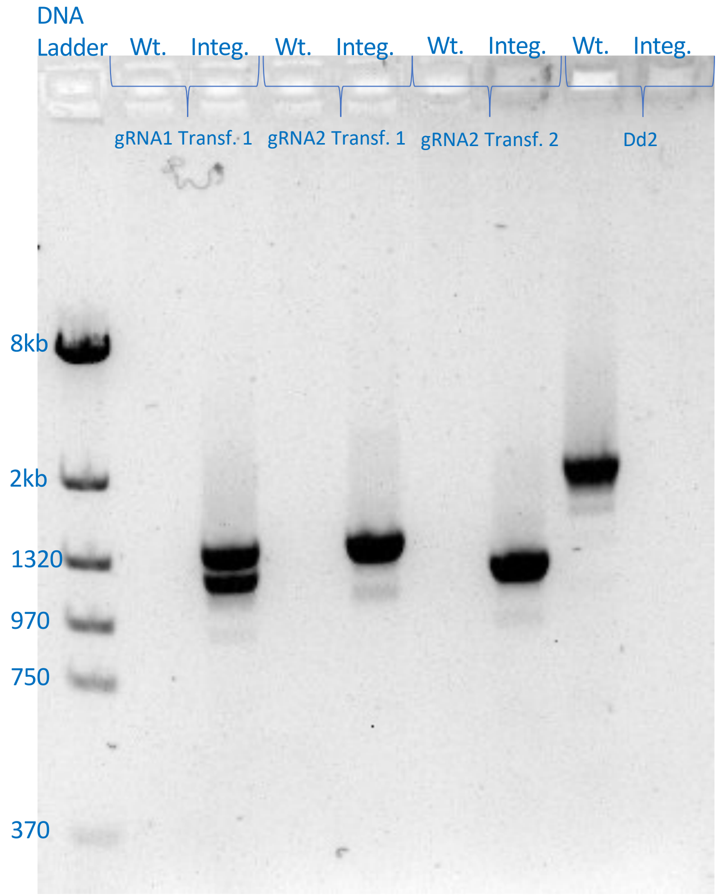


Unlabeled

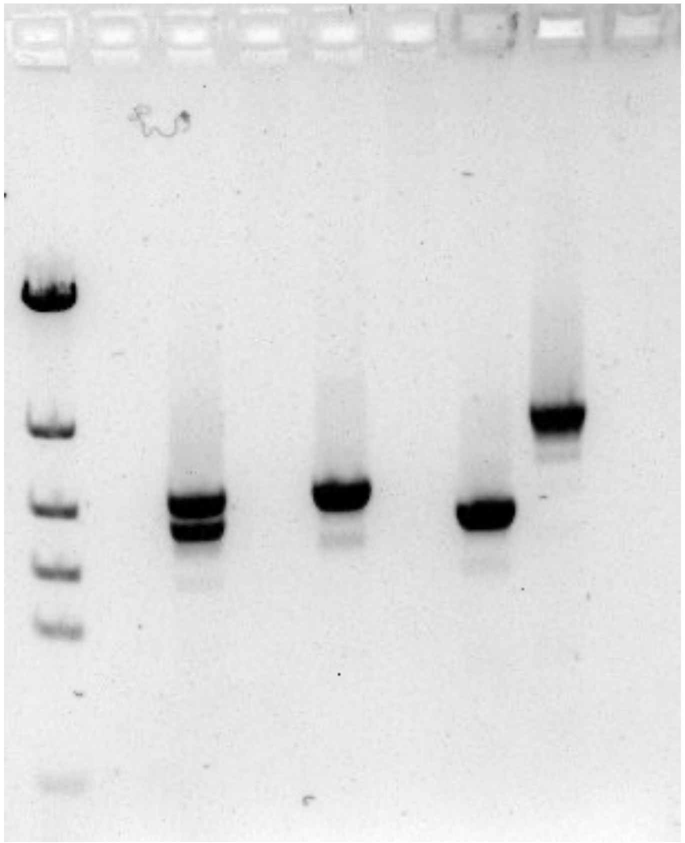

Figure 2 – source data 2
Uncropped WB for detection of ACP-KD in Dd2 upon aTc washout

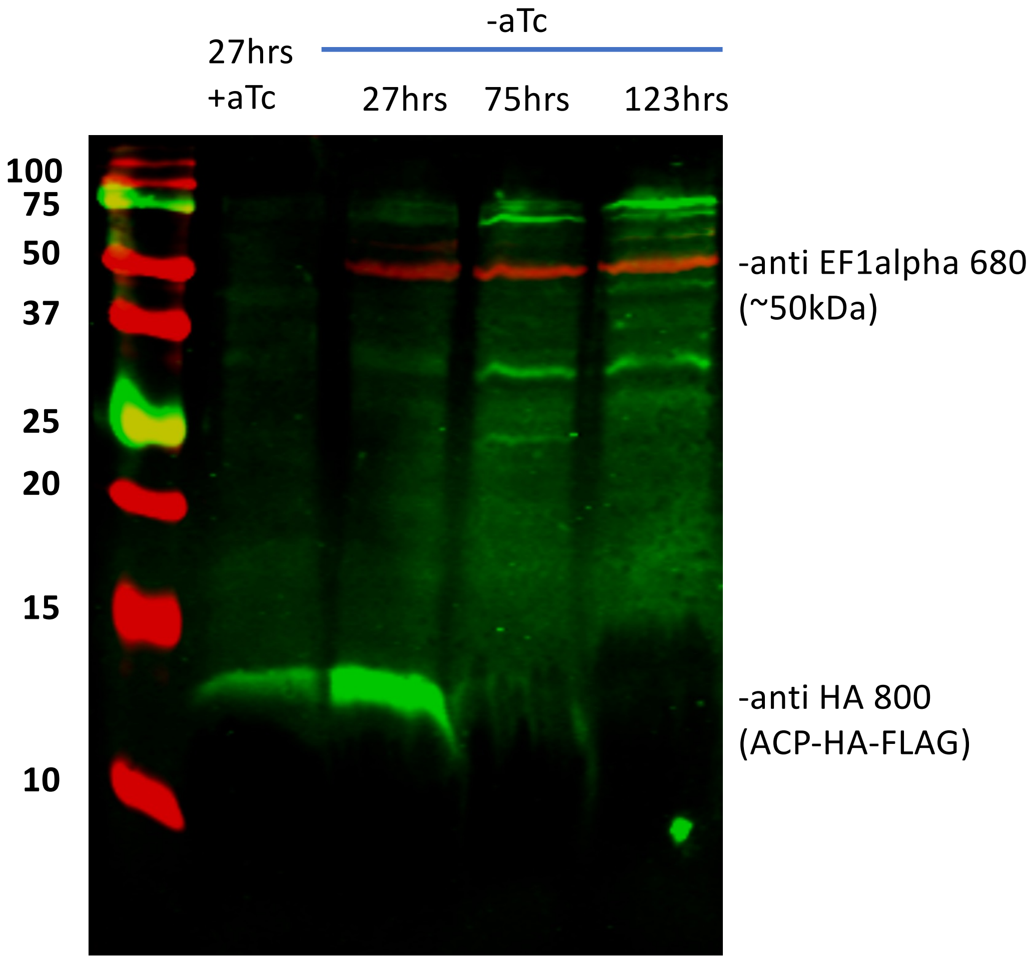


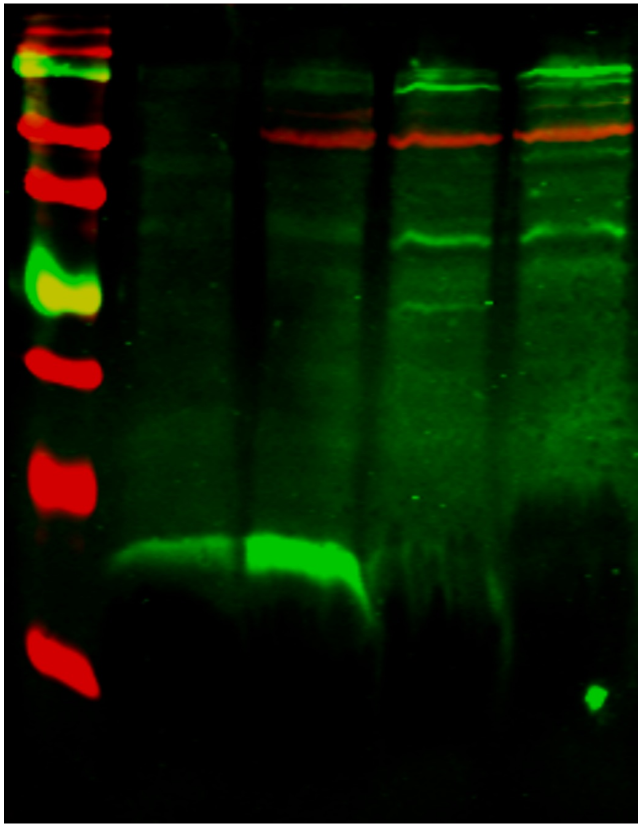

Figure 2 – source data 3
Uncropped gel image for western blot validation of custom anti-ACP antibody.

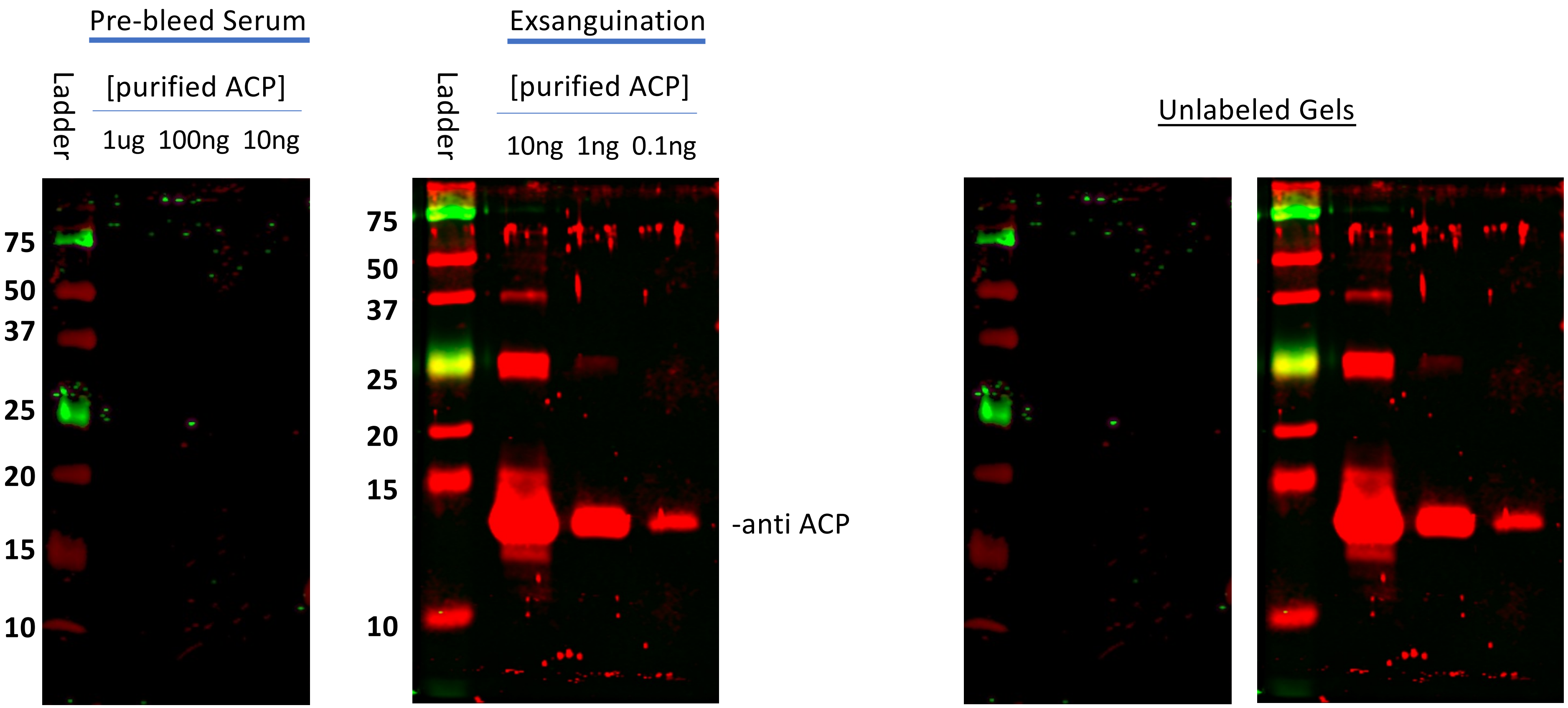

Figure 3 – source data 1

A. Uncropped PCR gels for deletion of ACPS in PfMev NF54

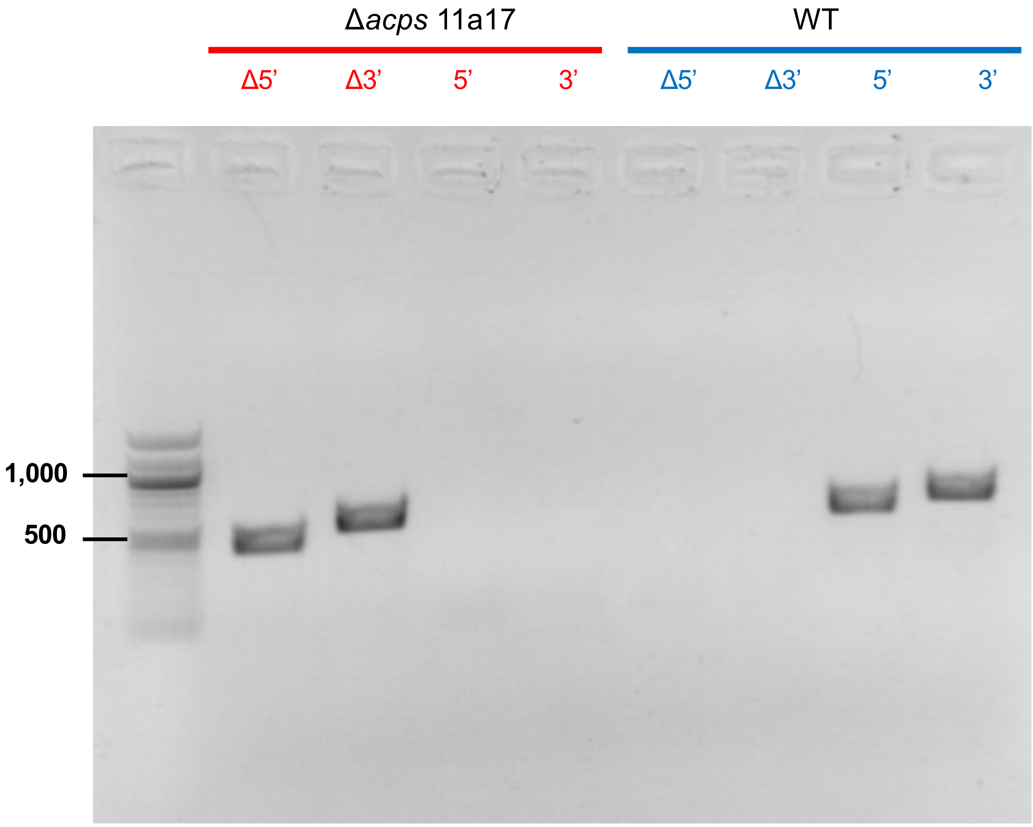


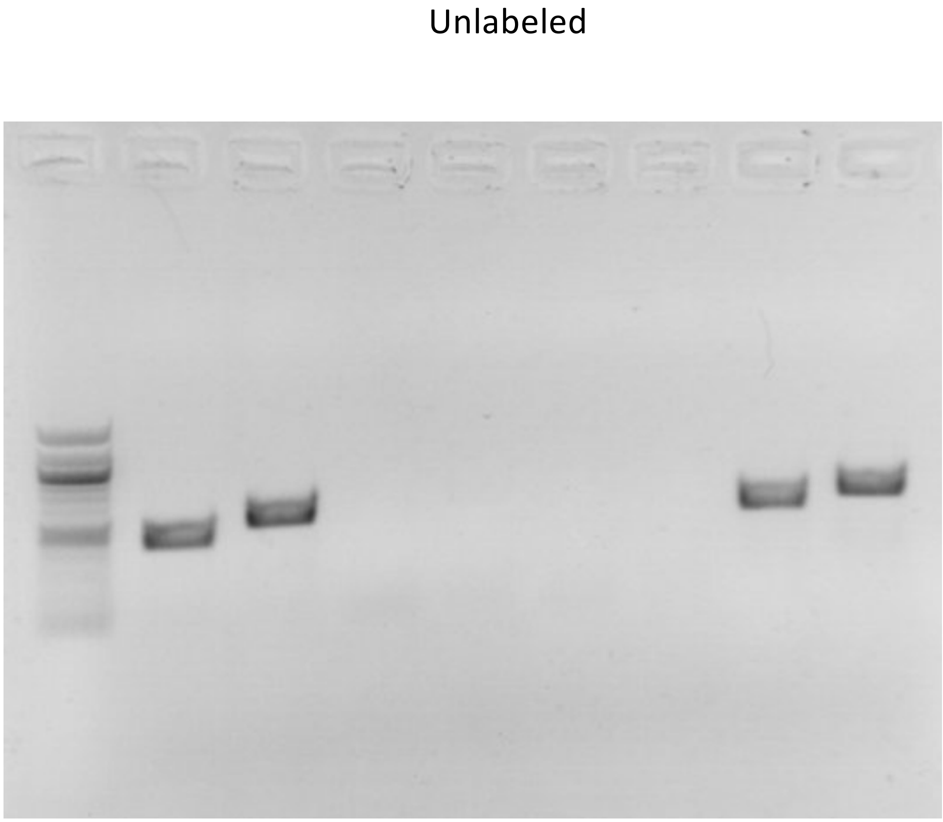

Figure 3 – Source Data 1

B. Uncropped PCR gels for organellar genomes in ACPS-KO and Wt PfMev NF54

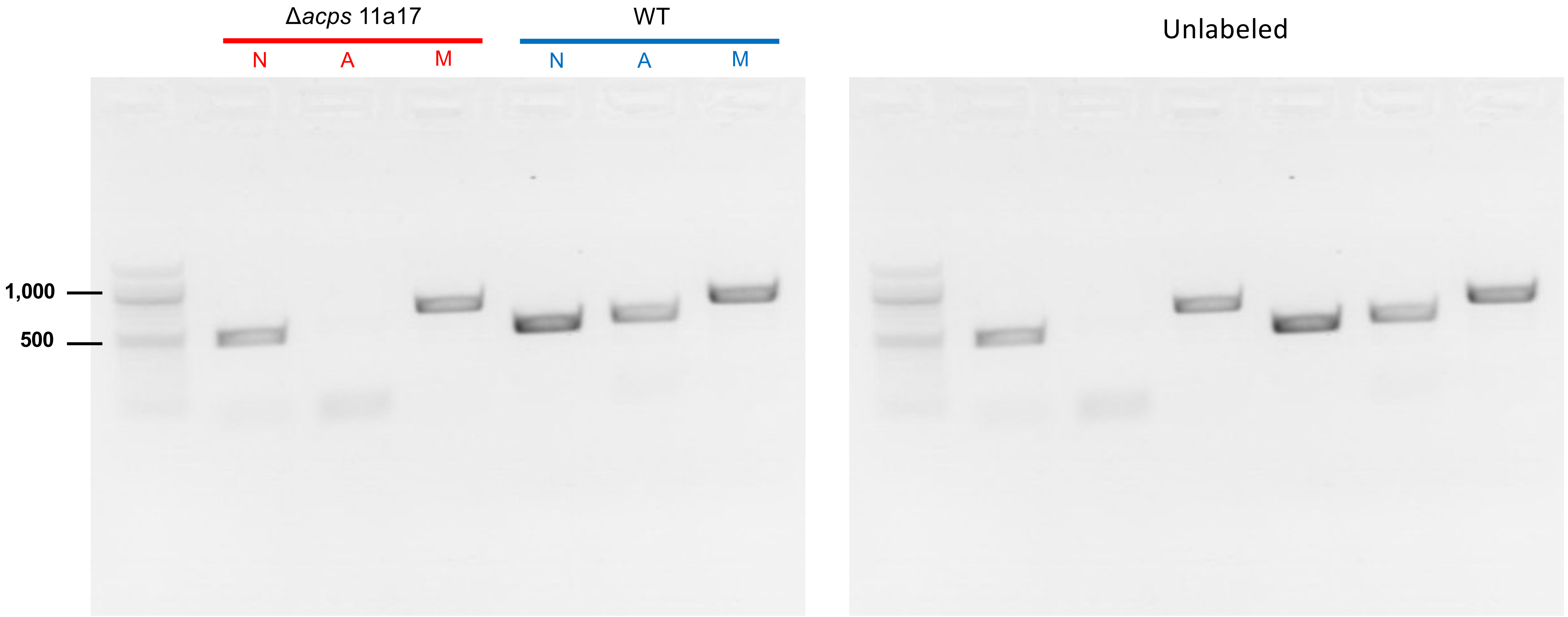


N = nuclear gene

A = apicoplast gene

M = mitochondrial gene

Figure 3 – source data 1

C. Uncropped PCR gels for deletion of FabD in PfMev NF54

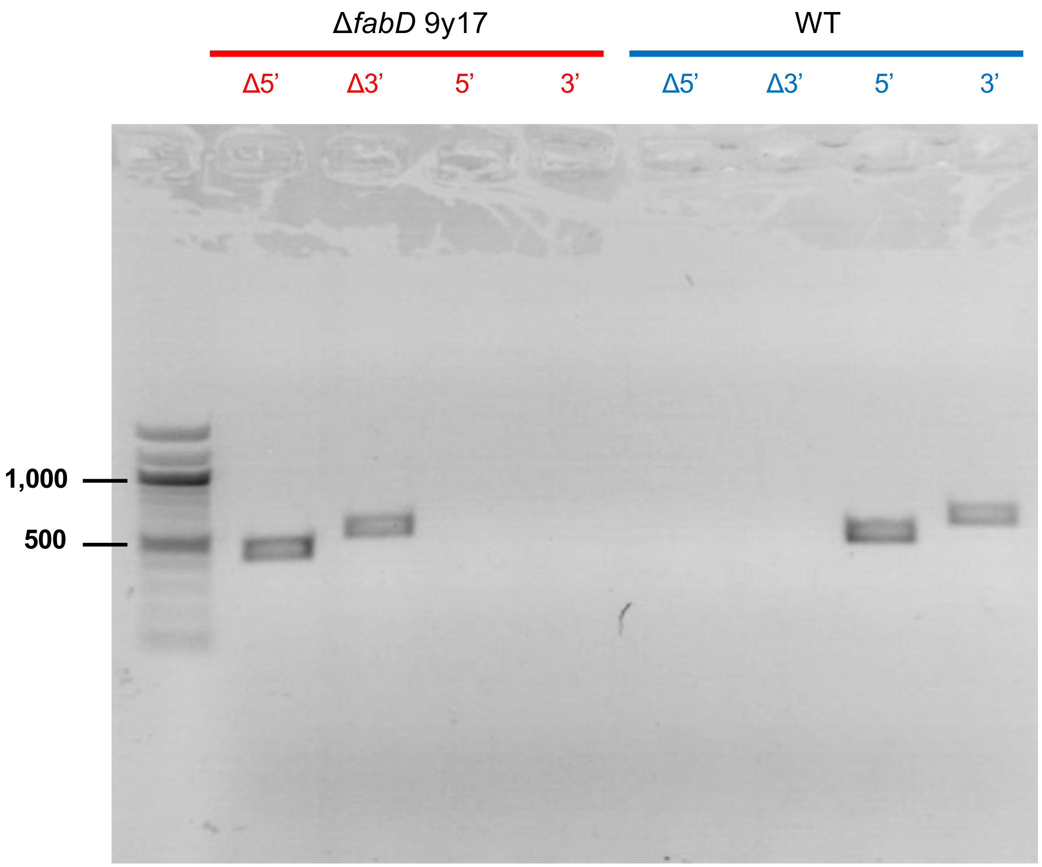


Unlabeled

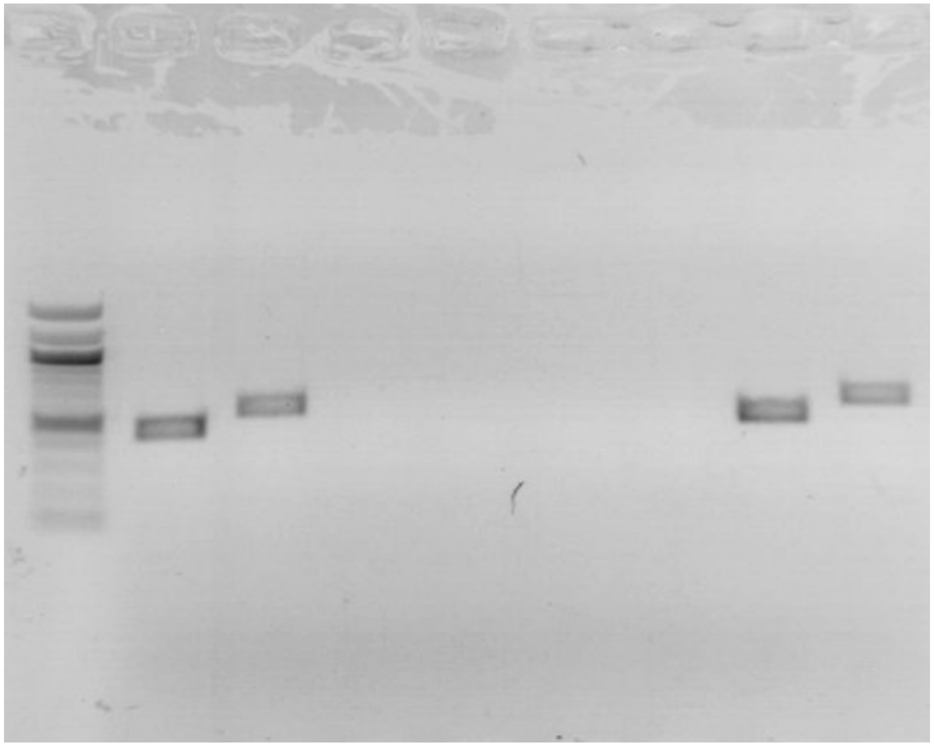

Figure 3 – source data 1

D. Uncropped PCR gels for organellar genomes in FabD-KO and PfMev NF54

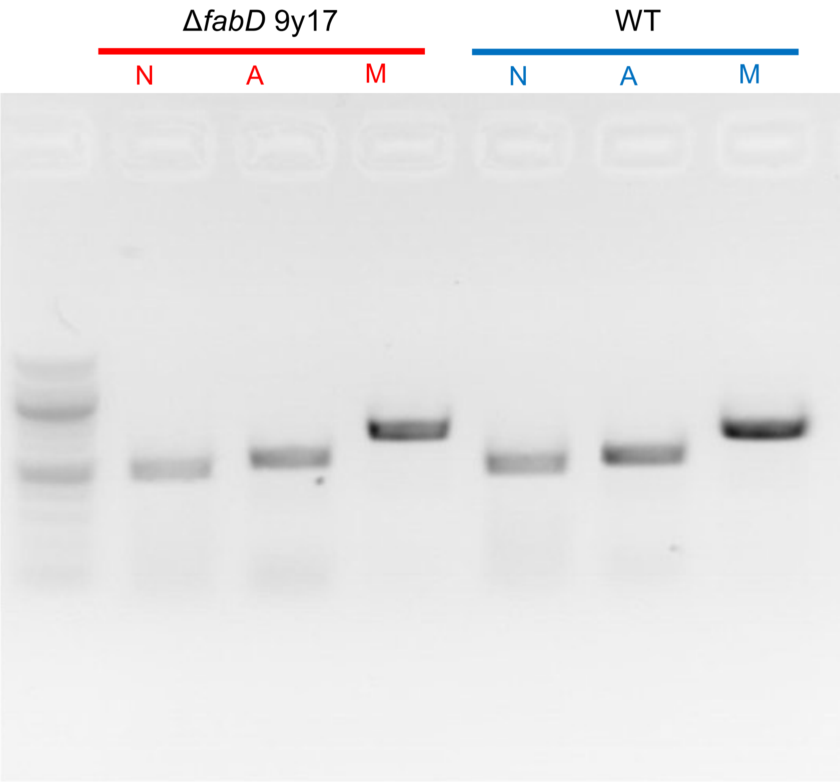


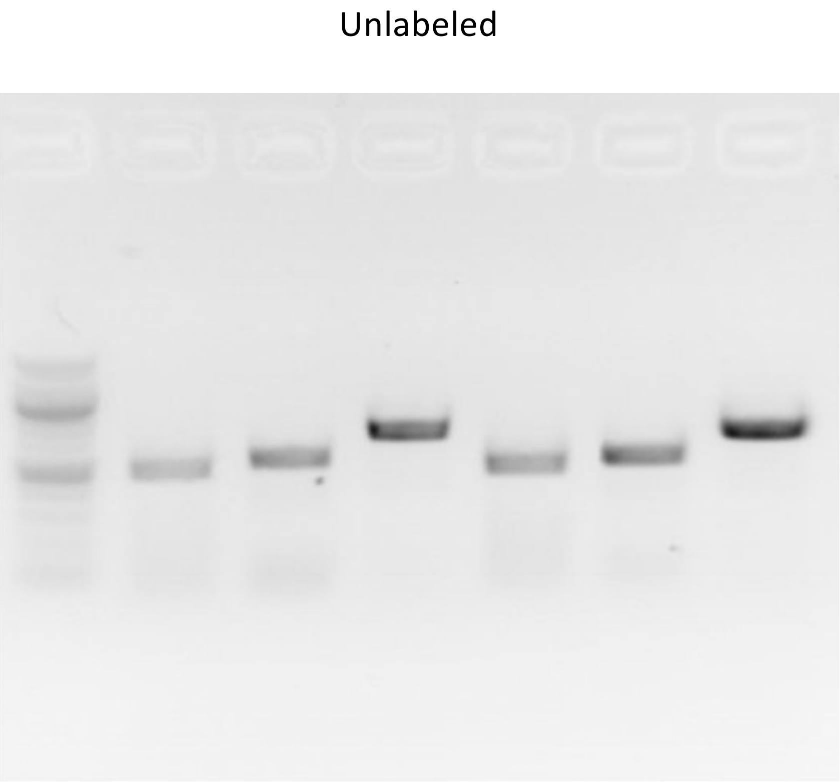


N = nuclear gene

A = apicoplast gene

M = mitochondrial gene

Figure 4 – source data 1
Uncropped blot for Figure 4- figure supplement 1

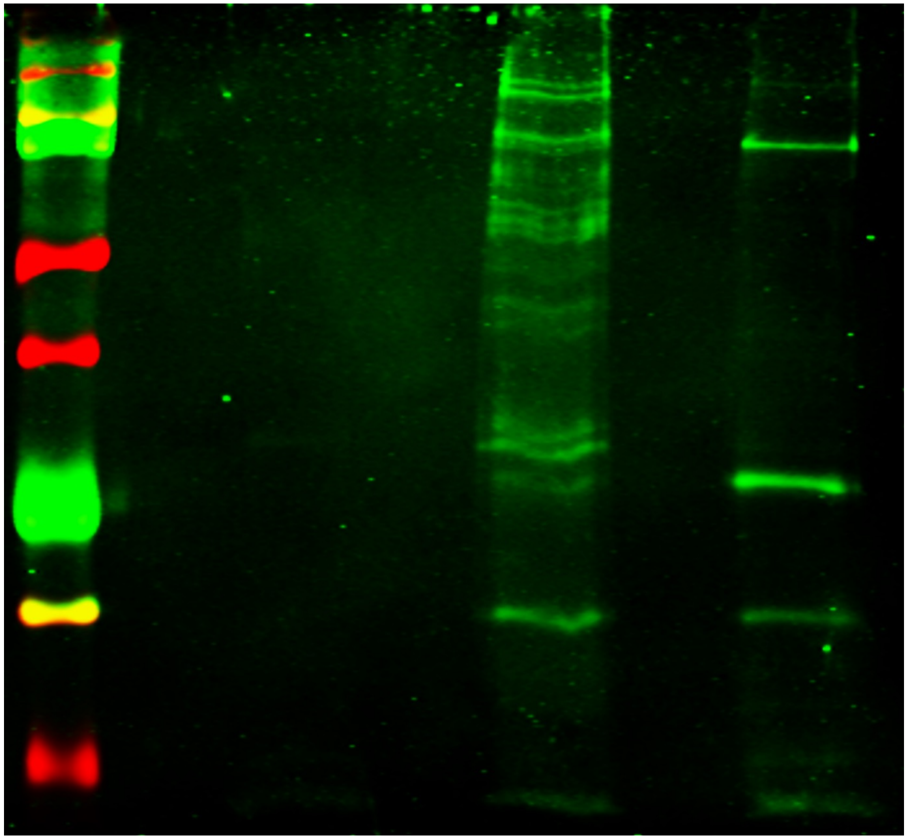

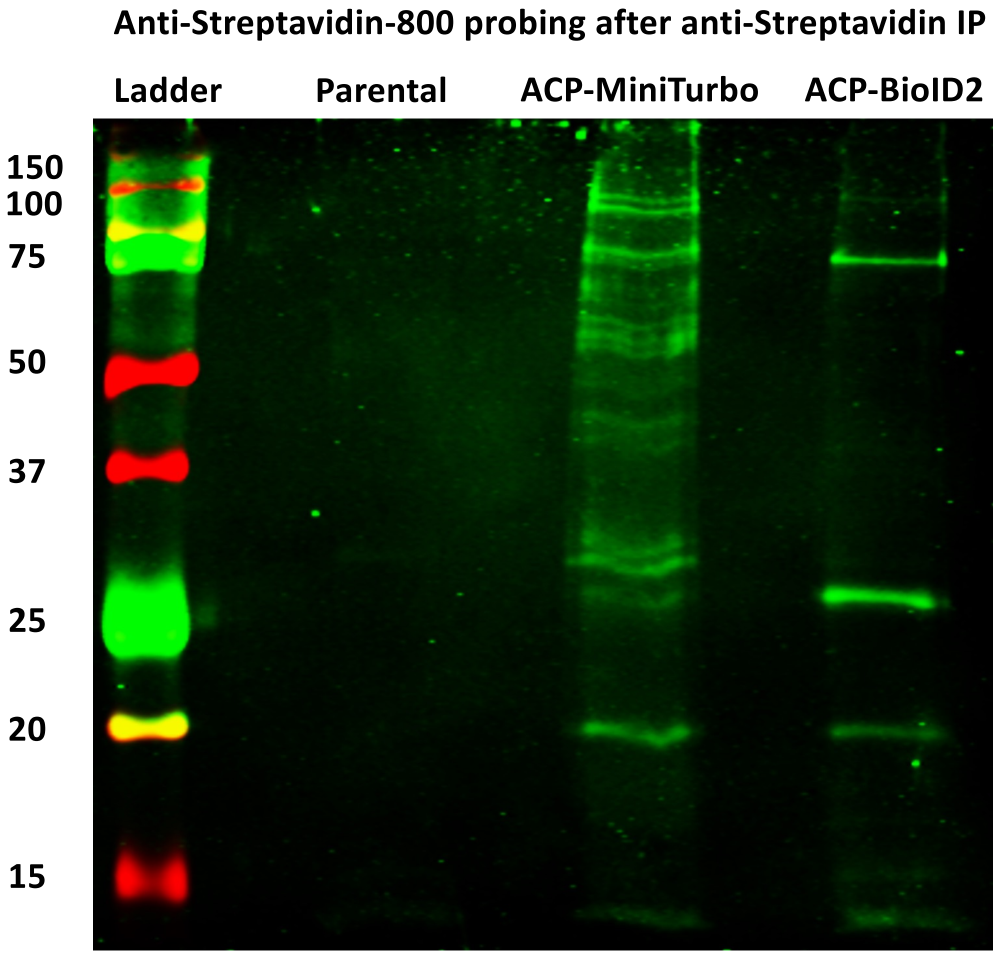

Figure 4 – source data 3
Uncropped Western Blots of immunoprecipitation of aACP and PKII in *E. coli*
*Replicate 1 – Probed for PKII*

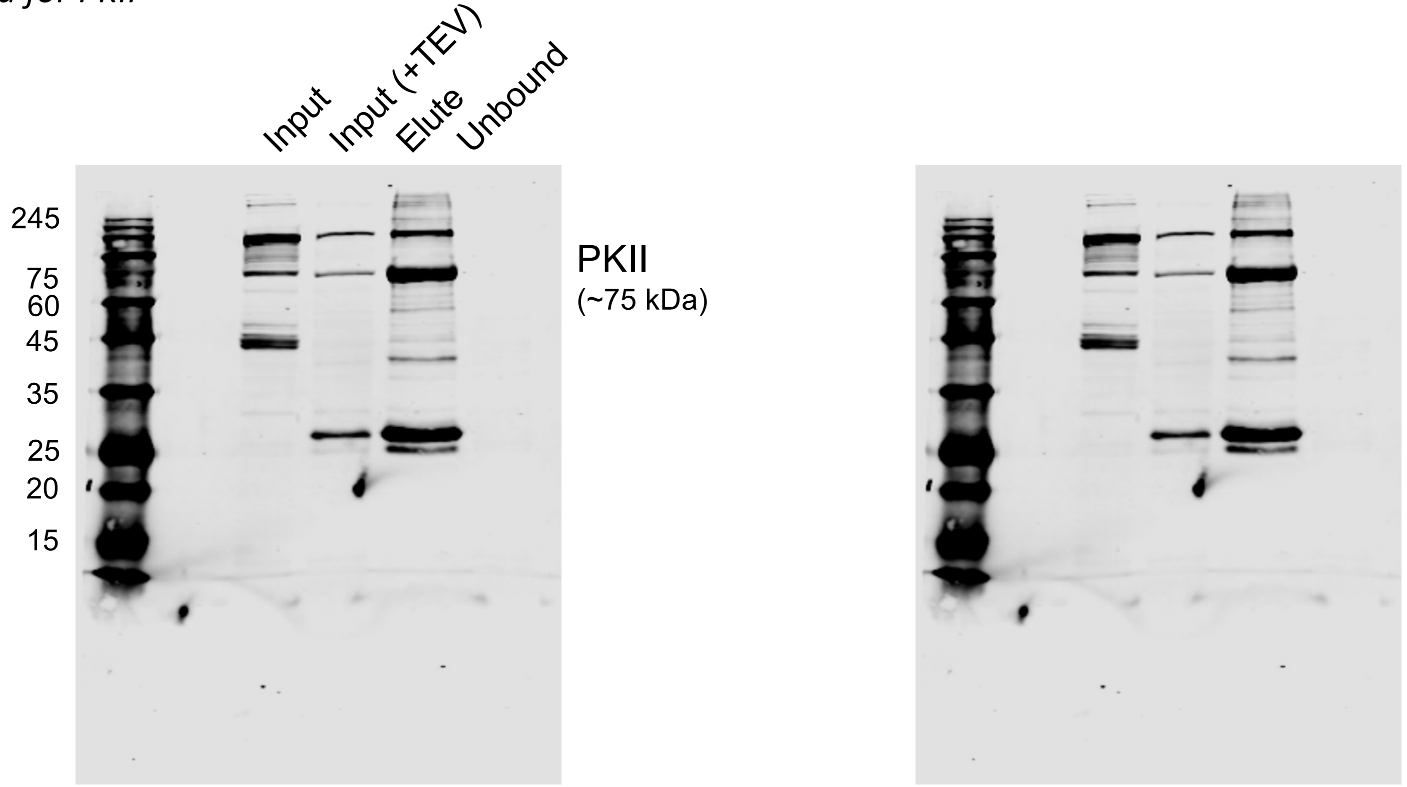


Labeled

Unlabeled

Uncleaved PKII 100 kDa
Cleaved PKII 75 kDa

Ni-NTA pulldown (PKII 6xHis)
TEV protease-treated

Figure 4 – source data 3
Uncropped Western Blots of immunoprecipitation of aACP and PKII in *E. coli*
*Replicate 1 – Probed for aACP*

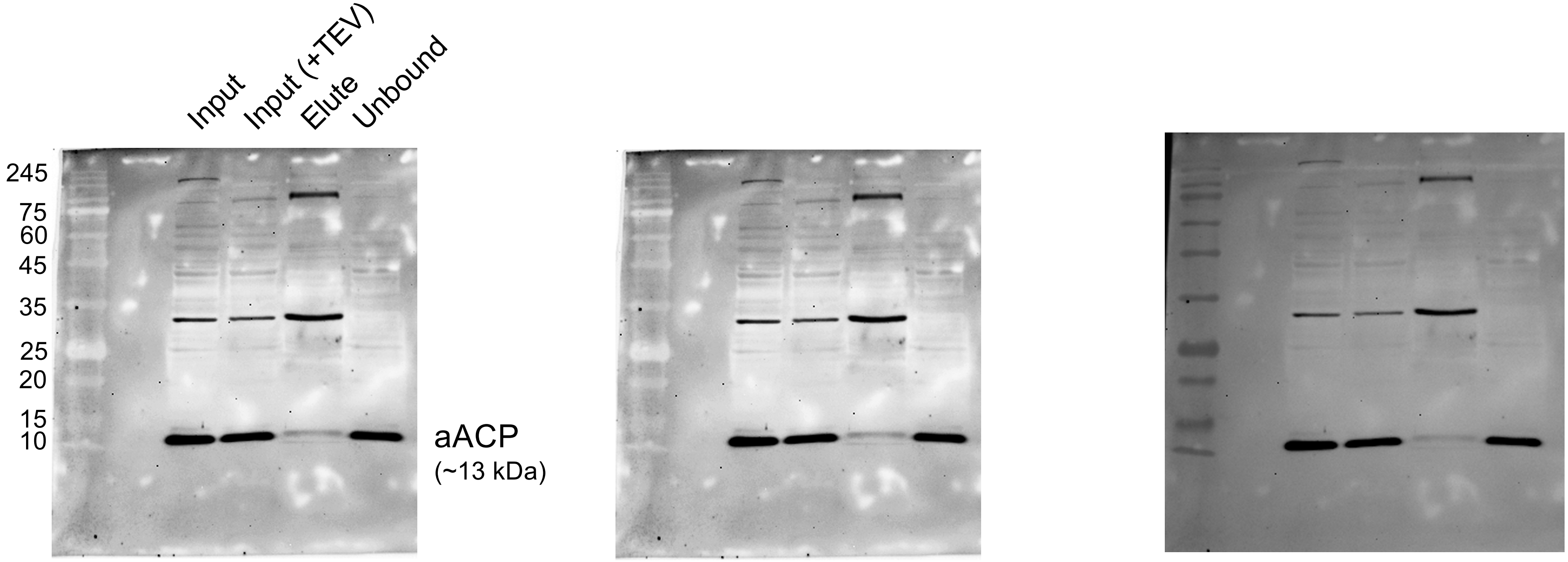


Chemi
Labeled

Chemi
Unlabeled

Membrane Overlay
Unlabeled

Ni-NTA pulldown (PKII 6xHis)
TEV protease-treated

Figure 4 – source data 3

Uncropped Western Blots of immunoprecipitation of aACP and PKII in *E. coli*

*Replicate 2 – Probed for PKII*

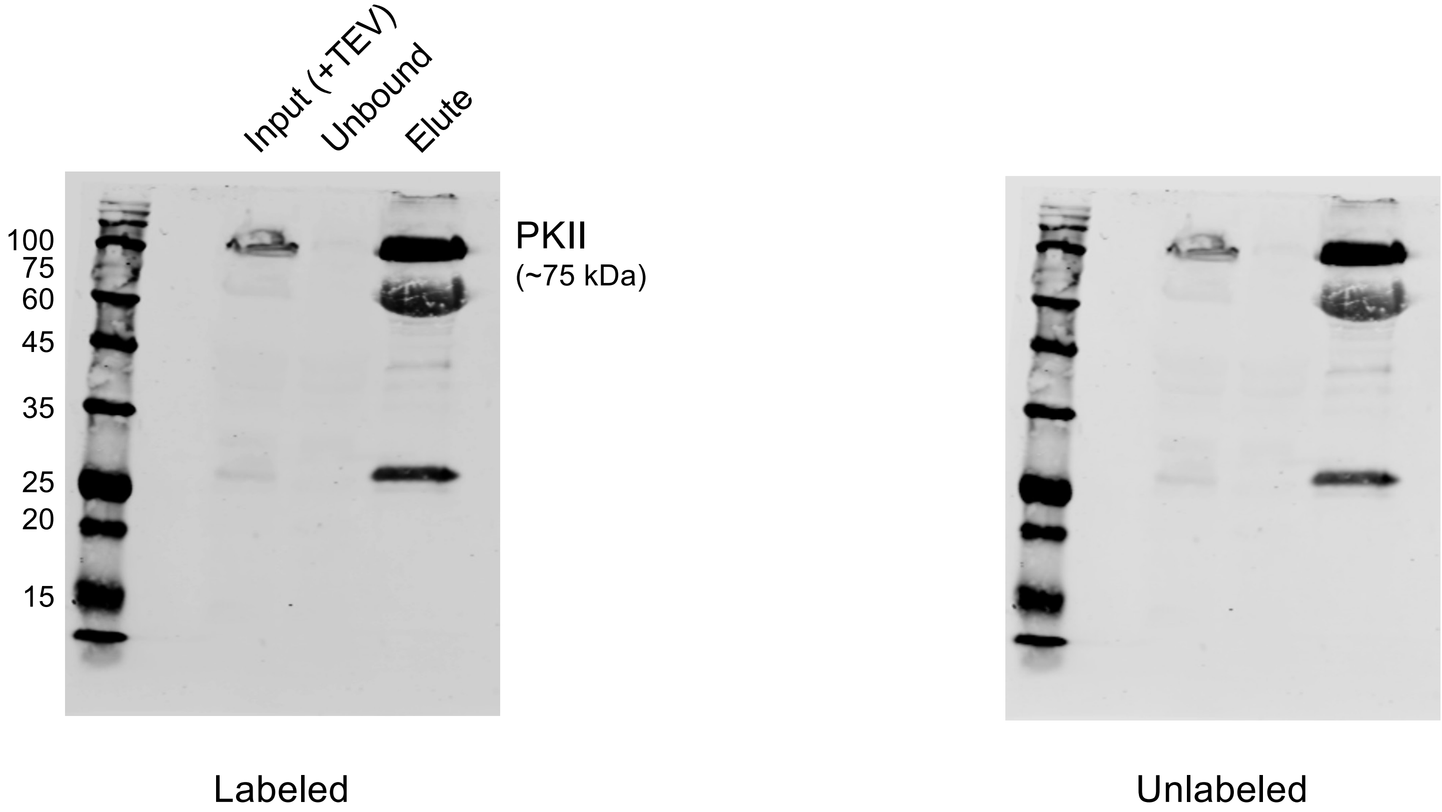


Ni-NTA pulldown (PKII 6xHis)
TEV protease-treated

Uncleaved PKII 100 kDa
Cleaved PKII 75 kDa

Figure 4 – source data 3
Uncropped Western Blots of immunoprecipitation of aACP and PKII in *E. coli*
*Replicate 2 – Probed for aACP*

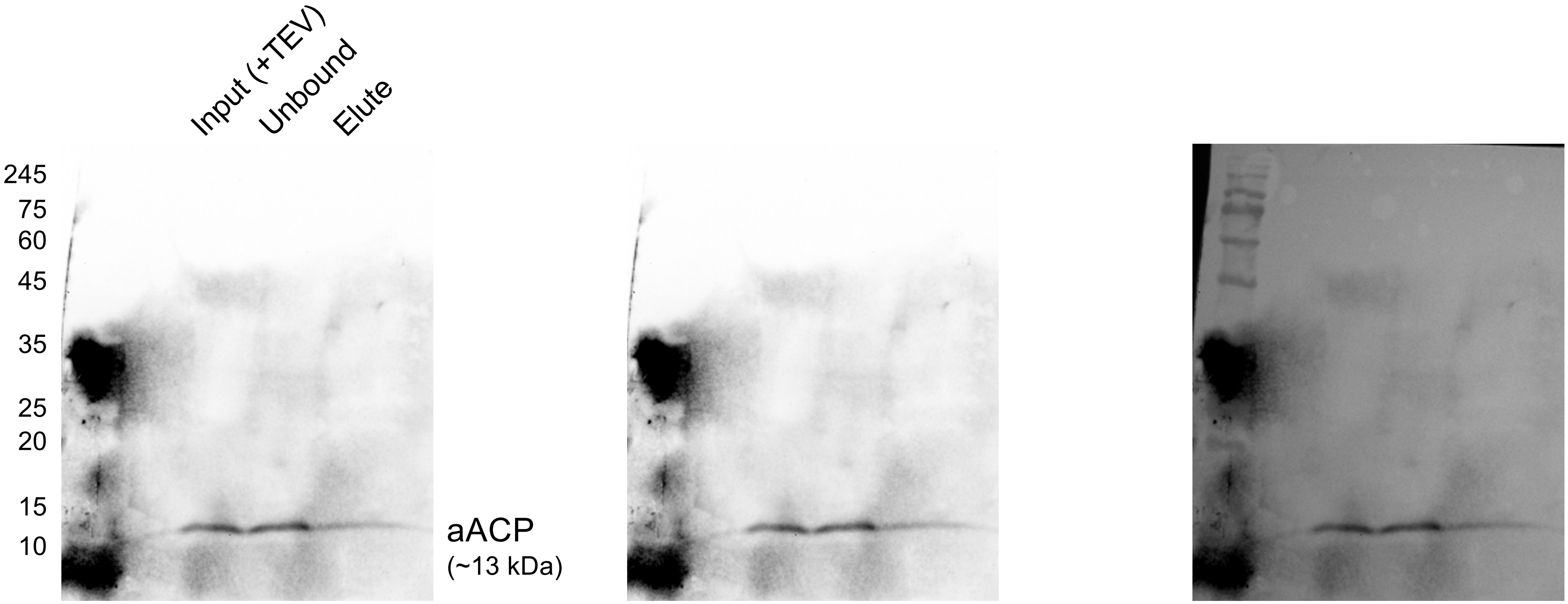


Chemi
Labeled

Chemi
Unlabeled

Membrane Overlay
Unlabeled

Ni-NTA pulldown (PKII 6xHis)
TEV protease-treated

Figure 4 – source data 3

Uncropped Western Blots of immunoprecipitation of aACP and PKII in *E. coli*

*Replicate 3 – Probed for PKII*

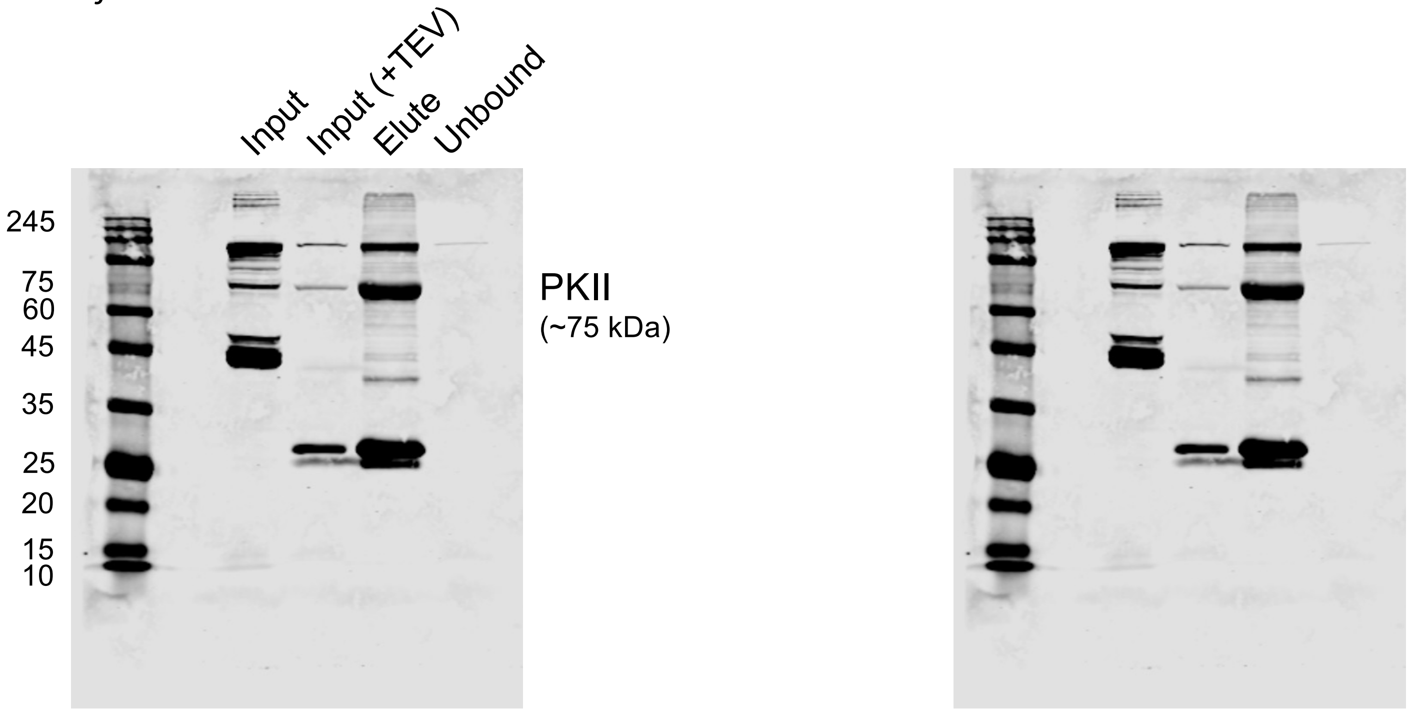


Labeled

Unlabeled

Ni-NTA pulldown (PKII 6xHis)
TEV protease-treated

Uncleaved PKII 100 kDa
Cleaved PKII 75 kDa

Figure 4 – source data 3

Uncropped Western Blots of immunoprecipitation of aACP and PKII in *E. coli*

*Replicate 3 – Probed for aACP*

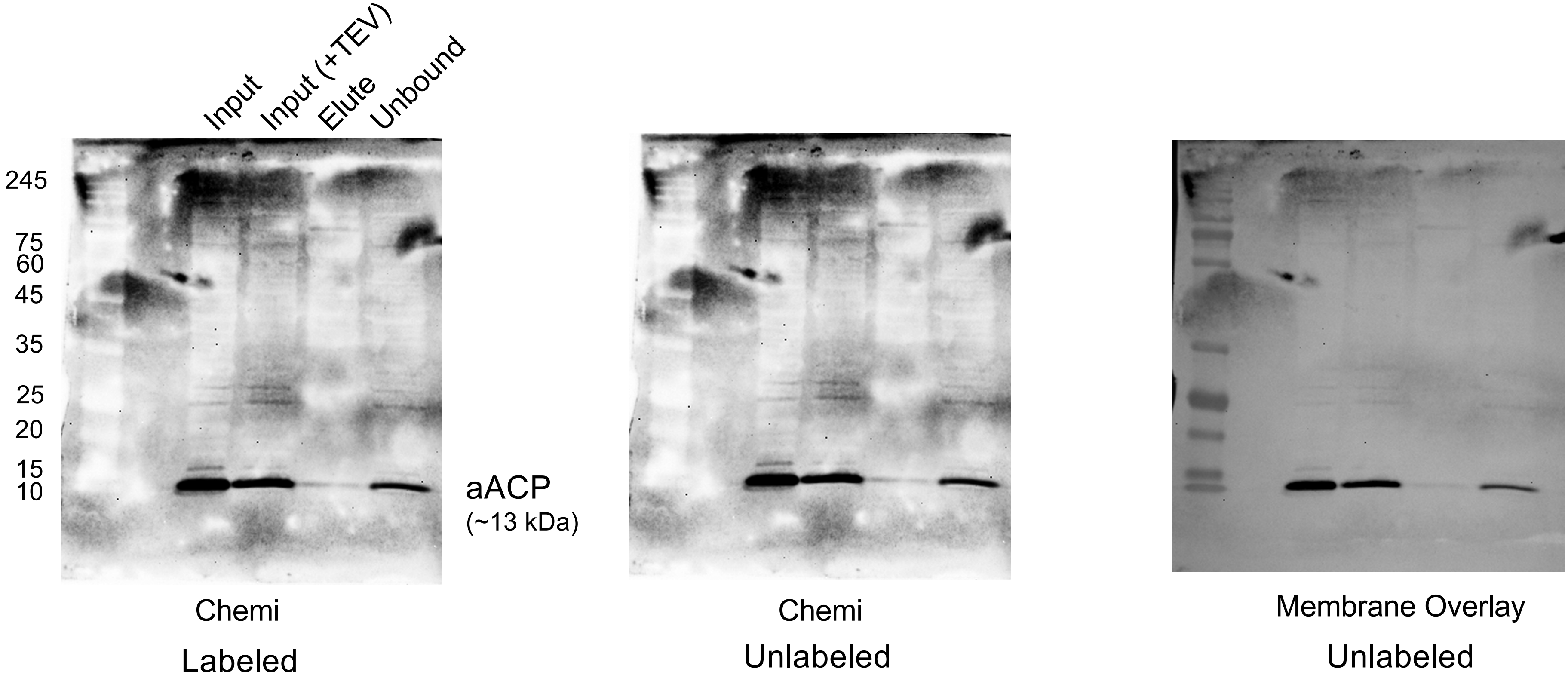


Ni-NTA pulldown (PKII 6xHis)
TEV protease-treated

Figure 4 – source data 3

Uncropped Western Blots of immunoprecipitation of aACP S95A and PKII in *E. coli*

*Replicate 1 – Probed for PKII*

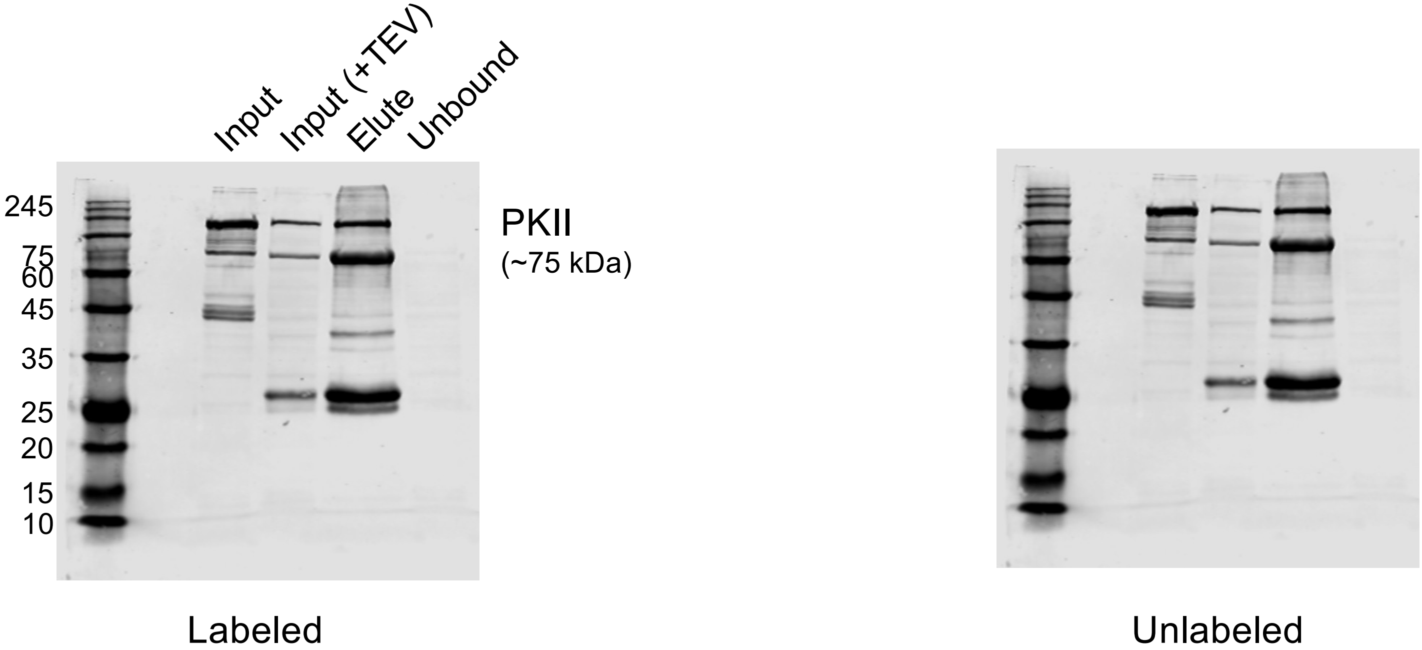


Ni-NTA pulldown (PKII 6xHis)
TEV protease-treated

Figure 4 – source data 3

Uncropped Western Blots of immunoprecipitation of aACP S95A and PKII in *E. coli*

*Replicate 1 – Probed for aACP*

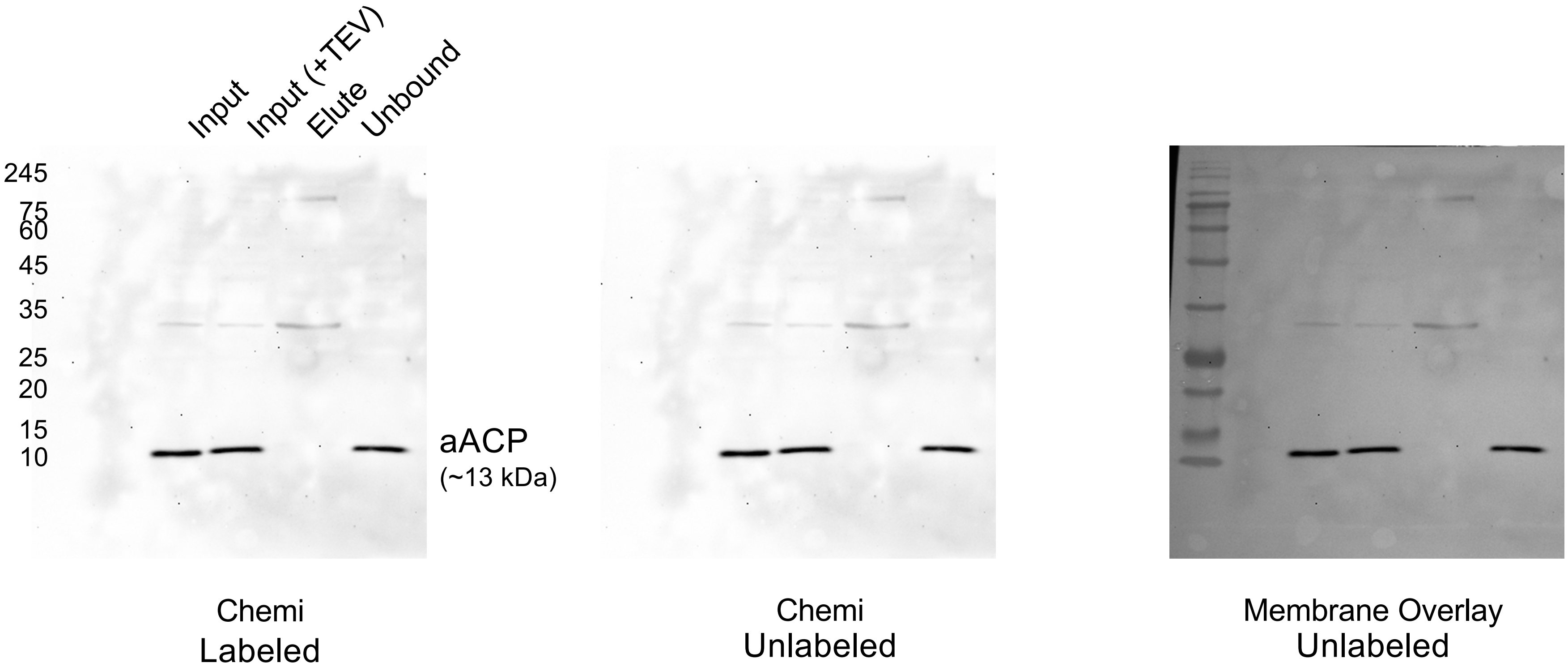


Ni-NTA pulldown (PKII 6xHis)
TEV protease-treated

Figure 4 – source data 3

Uncropped Western Blots of immunoprecipitation of aACP S95A and PKII in *E. coli*

*Replicate 2 – Probed for PKII*

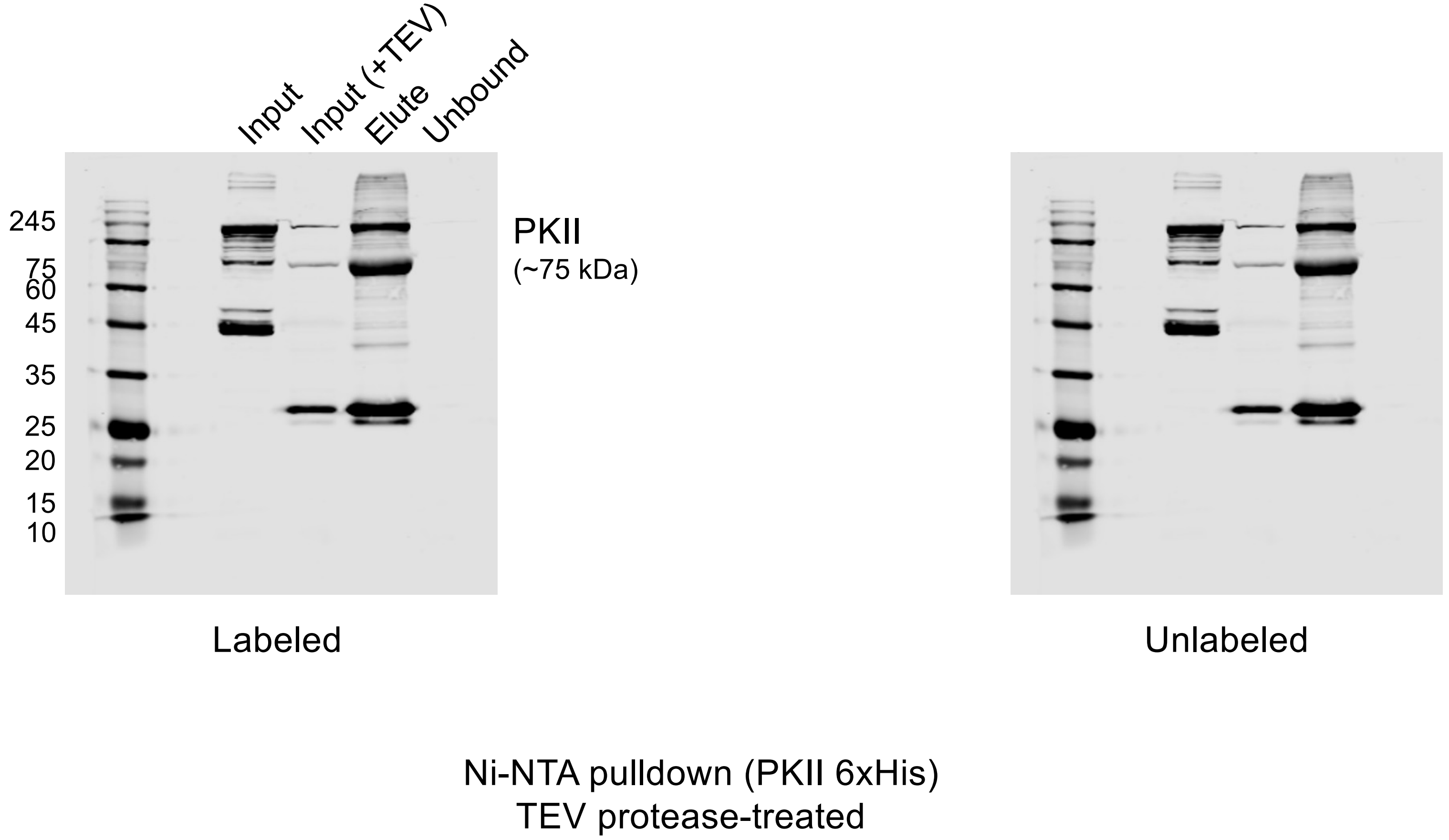

Figure 4 – source data 3

Uncropped Western Blots of immunoprecipitation of aACP S95A and PKII in *E. coli*

*Replicate 2 – Probed for aACP*

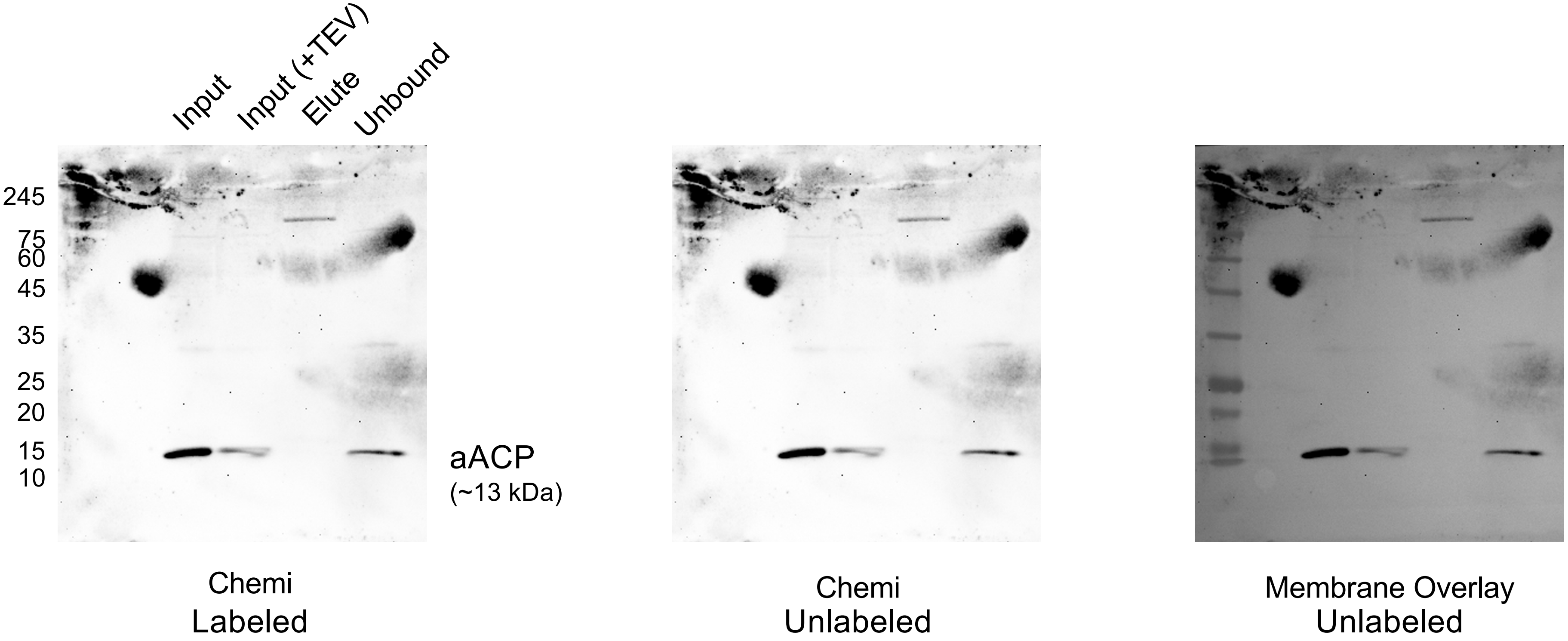


Ni-NTA pulldown (PKII 6xHis)
TEV protease-treated

Figure 4 – source data 3
Uncropped Western Blots of immunoprecipitation of aACP S95A and PKII in *E. coli*
*Replicate 3 – Probed for PKII*

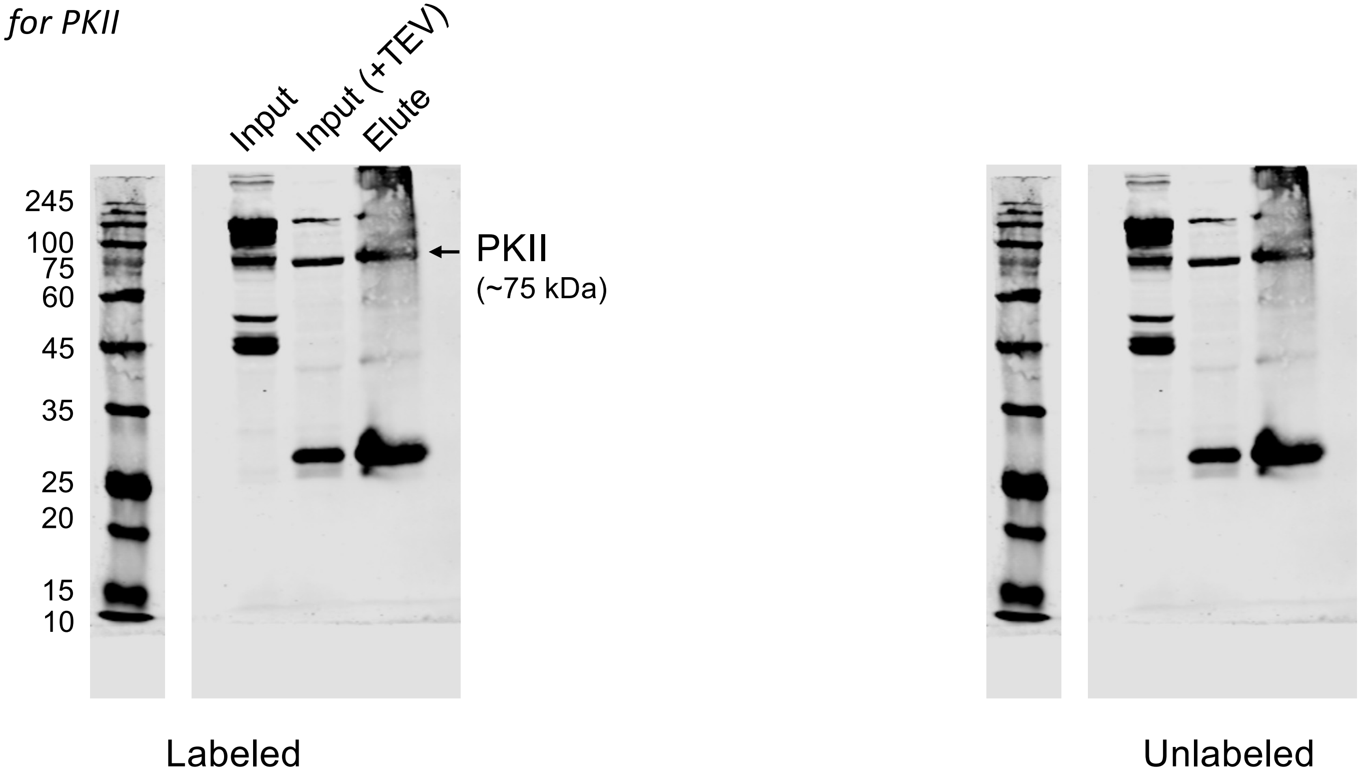


Ni-NTA pulldown (PKII 6xHis)
TEV protease-treated

Figure 4 – source data 3

Uncropped Western Blots of immunoprecipitation of aACP S95A and PKII in *E. coli*

*Replicate 3 – Probed for aACP*

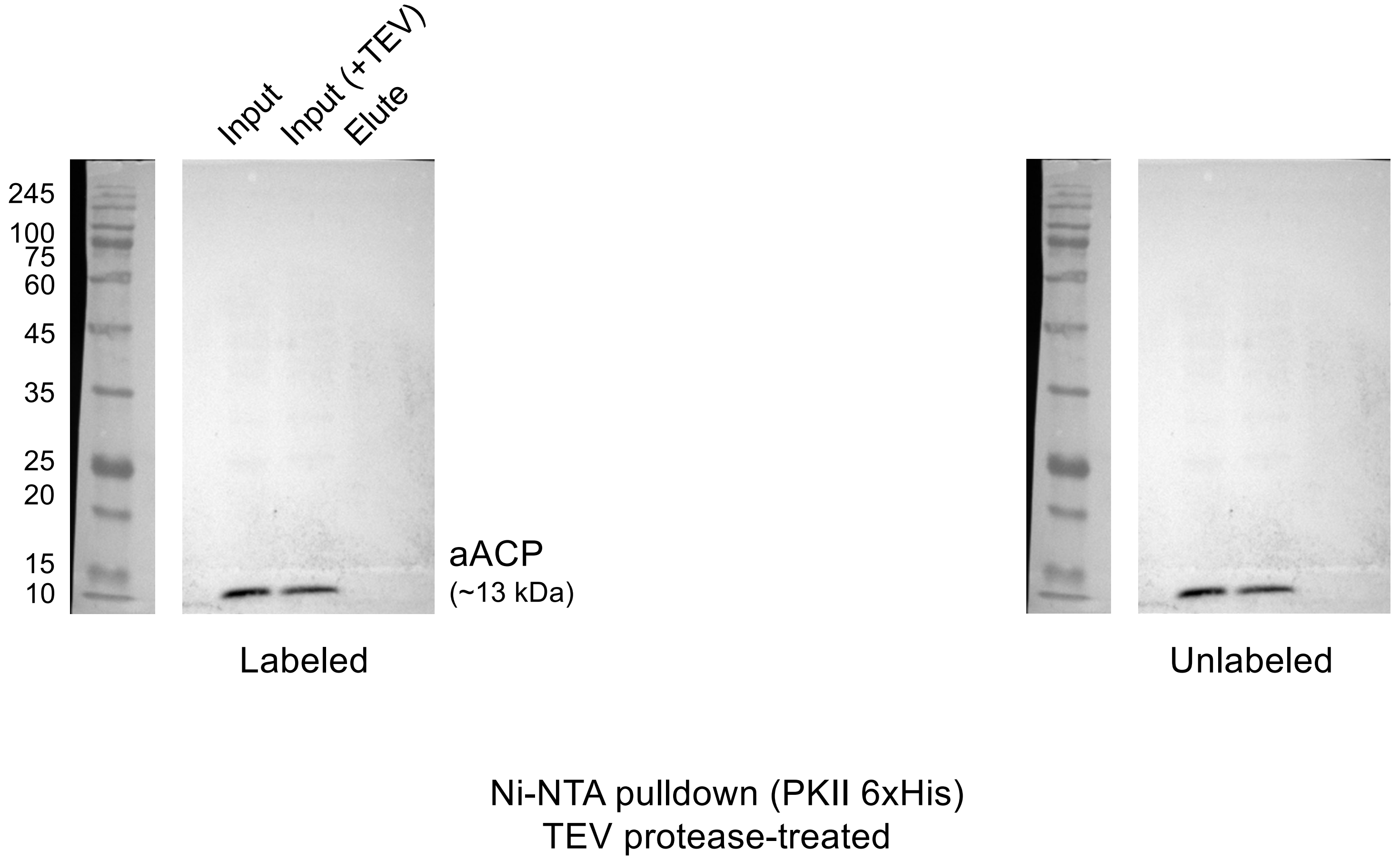

Figure 5 – source data 1
Uncropped Western Blots of PKII levels upon ACP-KD
*Replicate 1 – Probed for aACP*

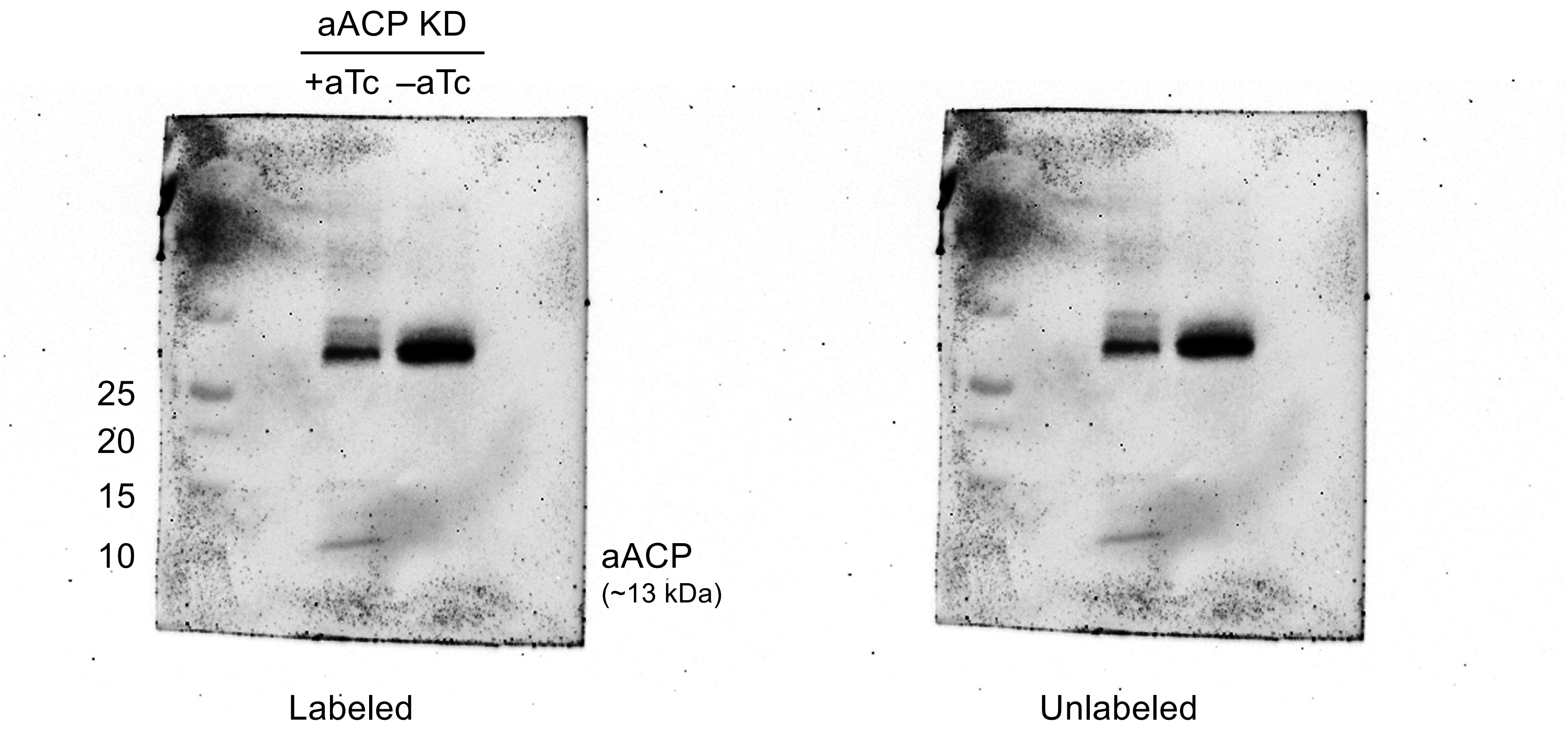

Figure 5 – source data 1

Uncropped Western Blots of PKII levels upon ACP-KD

*Replicate 1 – Probed for PKII*

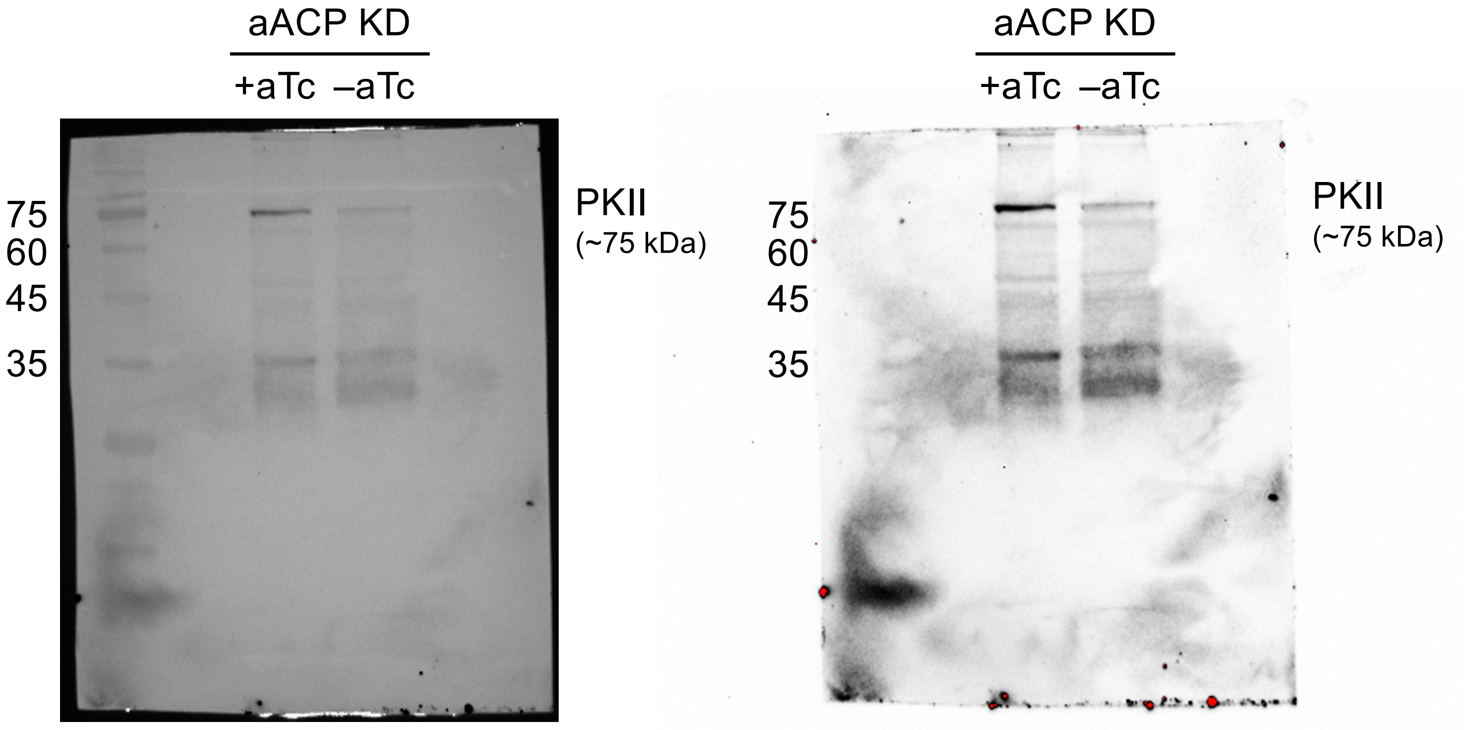


Membrane Overlay

Chemi

Labeled

Figure 5 – source data 1

Uncropped Western Blots of PKII levels upon ACP-KD

*Replicate 1 – Probed for PKII*

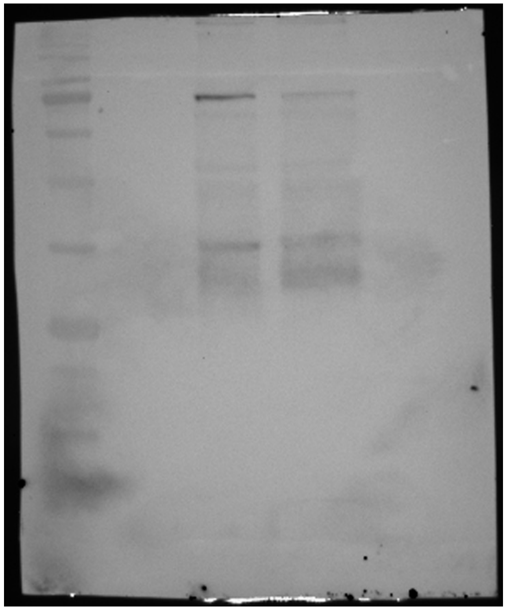

Membrane Overlay

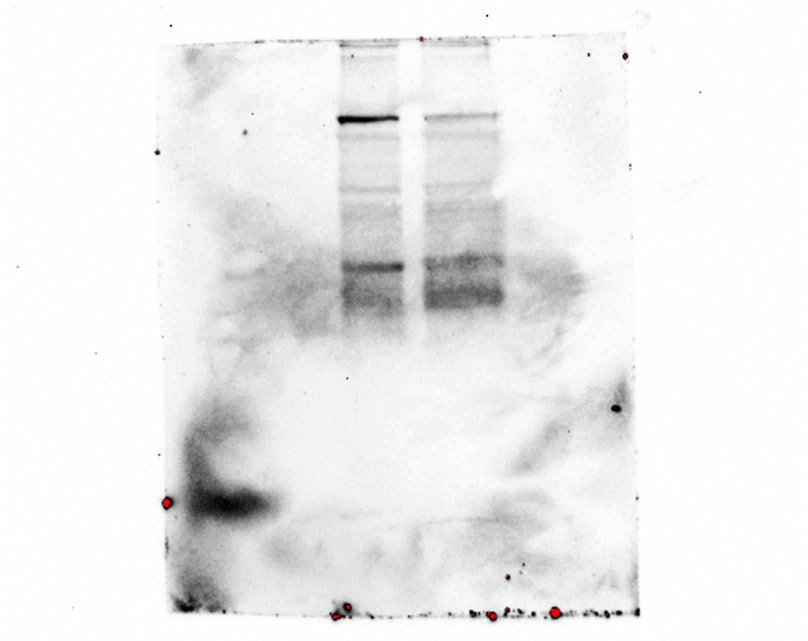

Chemi

Unlabeled

Figure 5 – source data 1

Uncropped Western Blots of PKII levels upon ACP-KD

*Replicate 1 – Probed for EF1-beta*

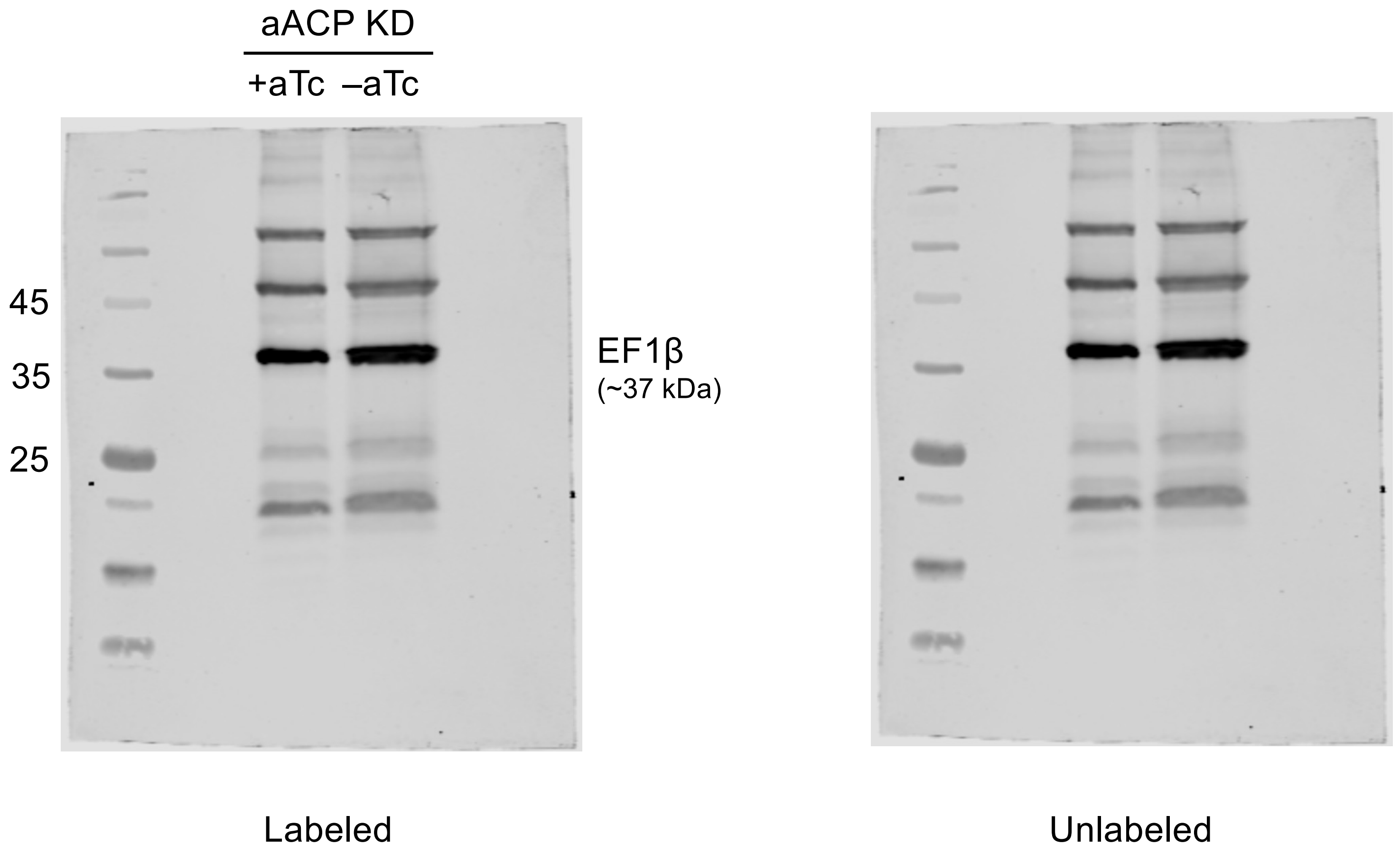

Figure 5 – source data 1
Uncropped Western Blots of PKII levels upon ACP-KD
*Replicate 1 – Ponceau S (Loading Control)*

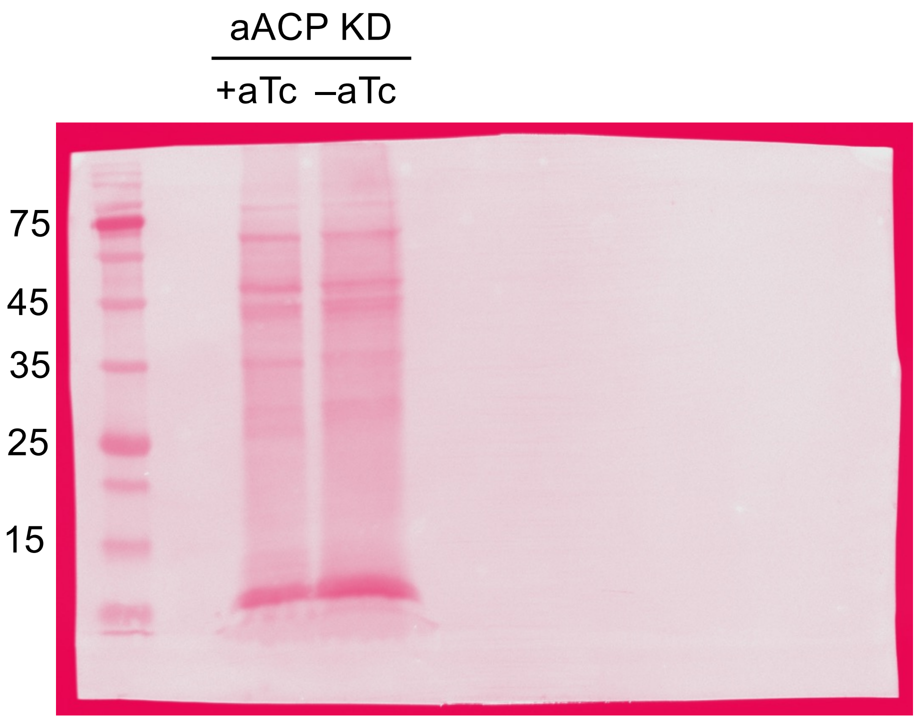


Labeled

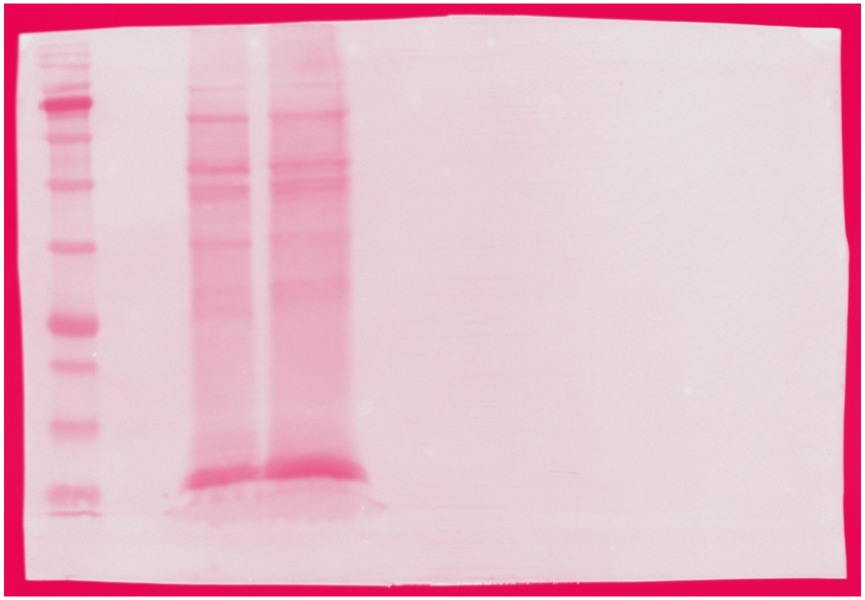

Unlabeled

Figure 5 – source data 1

Uncropped Western Blots of PKII levels upon ACP-KD

*Replicate 2 – Probed for PKII*

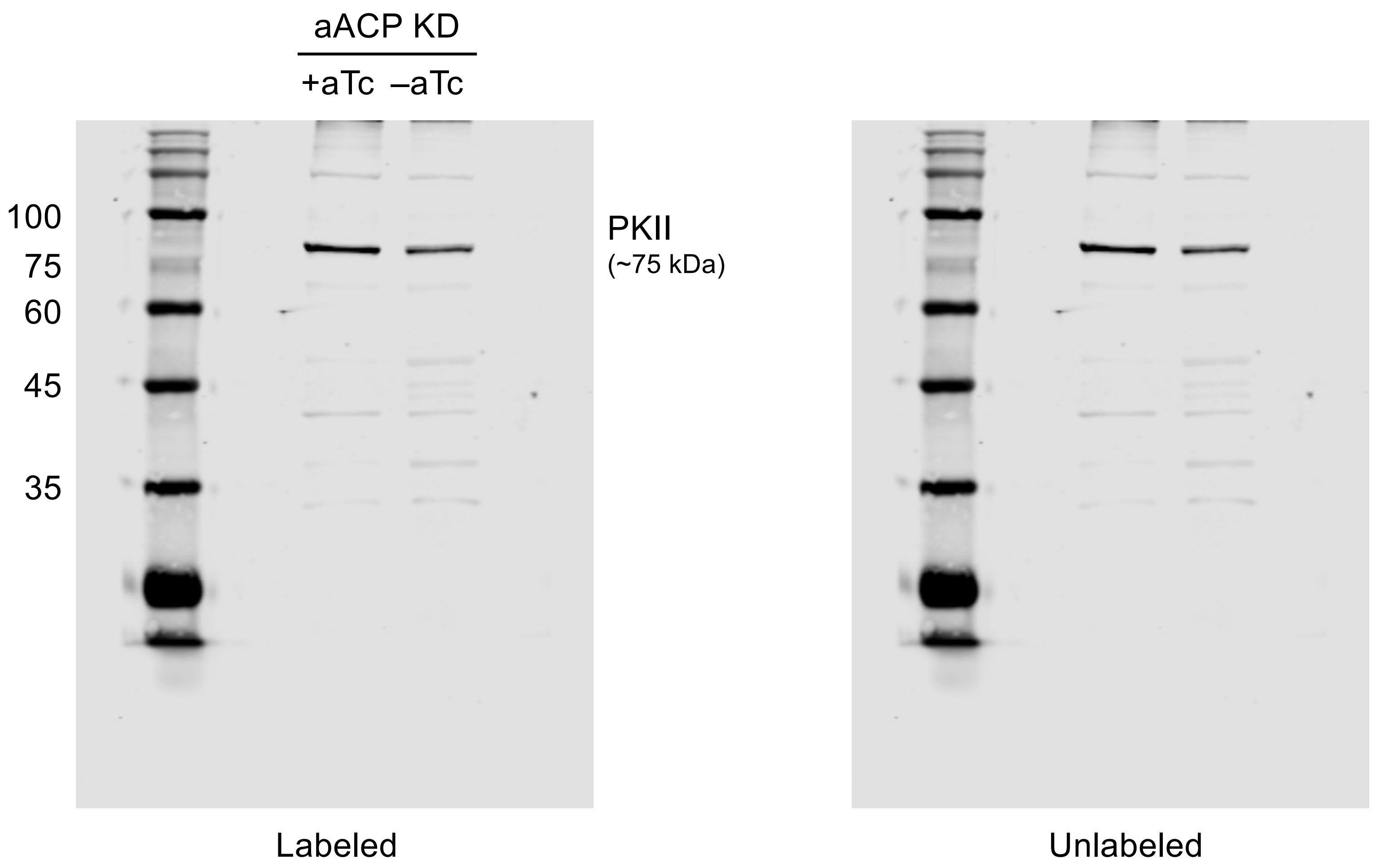

Figure 5 – source data 1

Uncropped Western Blots of PKII levels upon ACP-KD

*Replicate 2 – Probed for EF1-beta*

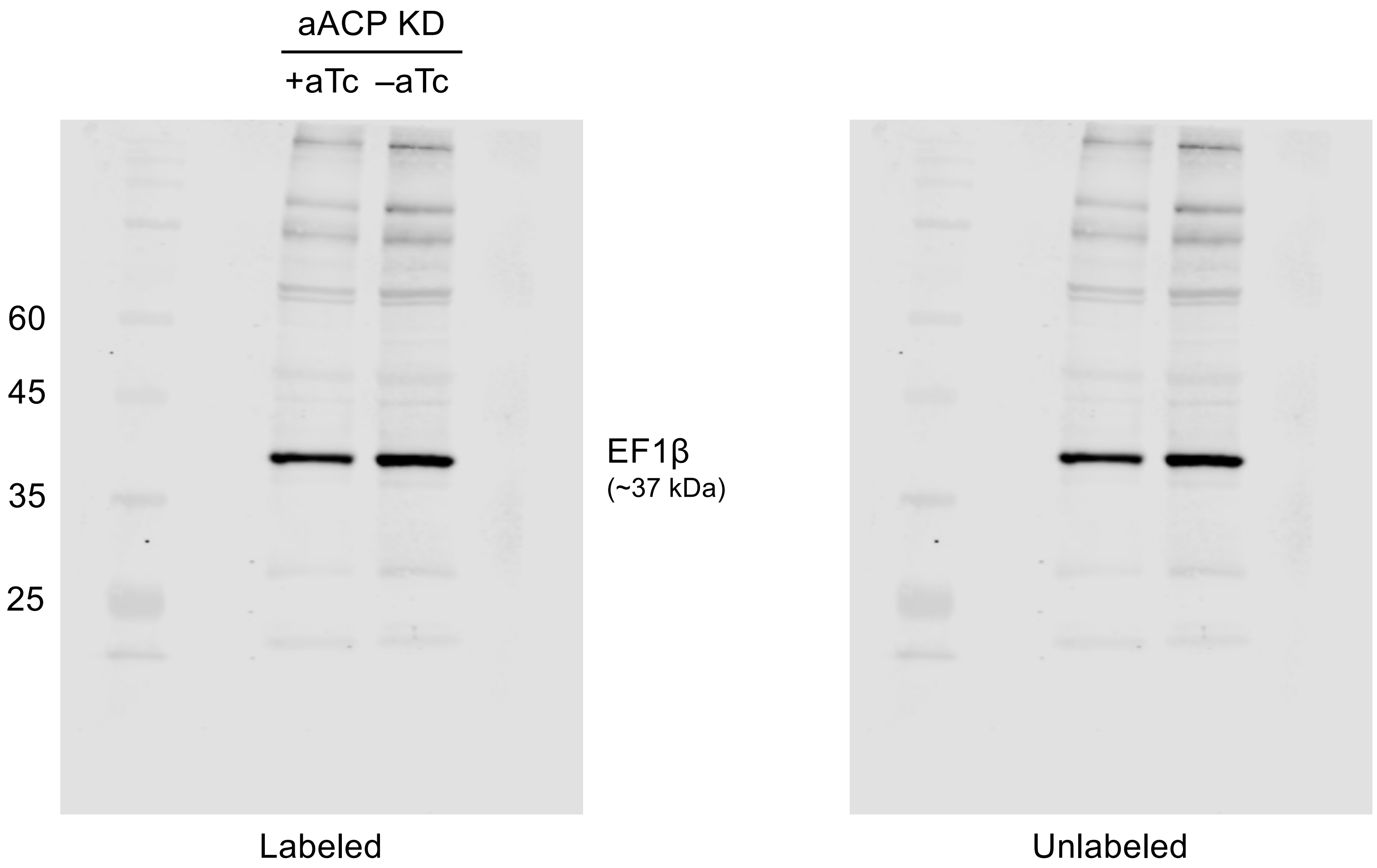

Figure 5 – source data 1
Uncropped Western Blots of PKII levels *upon ACP-KD*
*Replicate 3*

| | +aTc | | -aTc +IPP | |
|---|---|---|---|---|
| Ladder | Boiled | Sonicated | Boiled | Sonicated |

**Labeled**

75kDa
50kDa
37kDa
25kDa

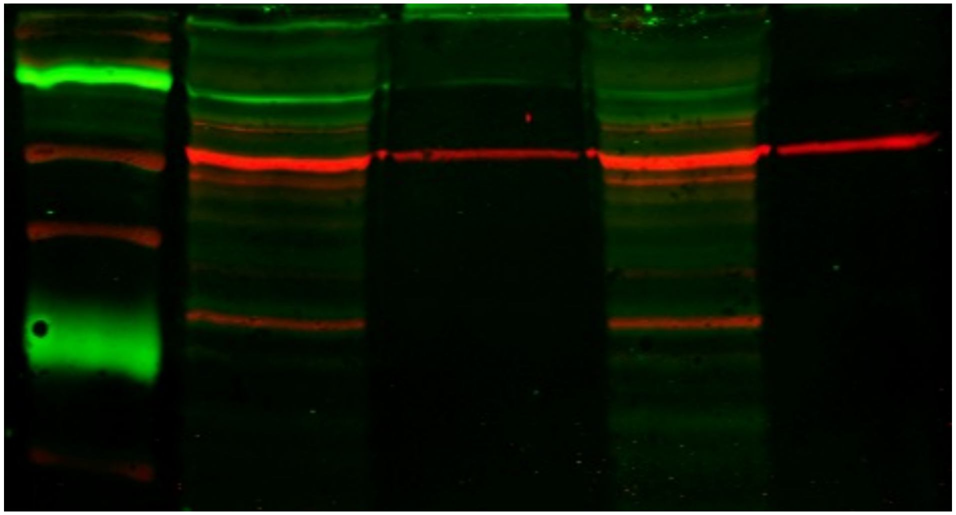

- PKII-3xMyc (~75kDa)
- EF1a (~50kDa)

Mouse anti-Myc
anti-Mouse 800

Rabbit anti-Ef1a
anti-Rabbit 680

**Unlabeled**

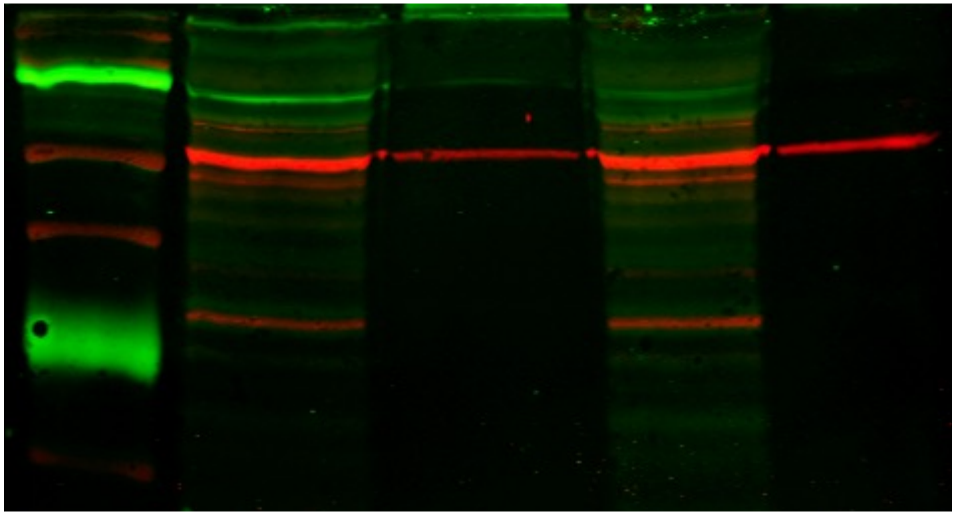
